# Sulfinyl Aziridines as Stereoselective Covalent Destabilizing Degraders of the Oncogenic Transcription Factor MYC

**DOI:** 10.1101/2025.02.24.639755

**Authors:** Hannah T. Rosen, Kelvin Li, Erin L. Li, Brynne Currier, Scott M. Brittain, Francisco J. Garcia, Diana C. Beard, Sandra Haenni-Holzinger, Dustin Dovala, Jeffrey M. McKenna, Markus Schirle, Thomas J. Maimone, Daniel K. Nomura

**Author notes:** Authors contributed equally to the work (Hannah T. Rosen, Kelvin Li).

## Abstract

While MYC is a significant oncogenic transcription factor driver of cancer, directly targeting MYC has remained challenging due to its intrinsic disorder and poorly defined structure, deeming it “undruggable.” Whether transient pockets formed within intrinsically disordered and unstructured regions of proteins can be selectively targeted with small molecules remains an outstanding challenge. Here, we developed a bespoke stereochemically-paired spirocyclic oxindole aziridine covalent library and screened this library for degradation of MYC. Through this screen, we identified a hit covalent ligand KL2-236, bearing a unique sulfinyl aziridine warhead, that engaged MYC *in vitro* as pure MYC/MAX protein complex and *in situ* in cancer cells to destabilize MYC, inhibit MYC transcriptional activity and degrade MYC in a proteasome-dependent manner through targeting intrinsically disordered C203 and D205 residues. Notably, this reactivity was most pronounced for specific stereoisomers of KL2-236 with a diastereomer KL4-019 that was largely inactive. Mutagenesis of both C203 and D205 completely attenuated KL2-236-mediated MYC degradation. We have also optimized our initial KL2-236 hit compound to generate a more durable MYC degrader KL4-219A in cancer cells. Our results reveal a novel ligandable site within MYC and indicate that certain intrinsically disordered regions within high-value protein targets, such as MYC, can be interrogated by isomerically unique chiral small molecules, leading to destabilization and degradation.

## Introduction

Among “undruggable” targets that have been identified, transcription factors have represented a particularly challenging class of targets for drug discovery due to the presence of large regions of intrinsic disorder and poorly defined structures for many members. Whether these unstructured regions of transcription factors have ligandable sites that can be pharmacologically targeted in a selective manner is often unclear ^1^.

MYC has been a particularly elusive target for direct targeting due to its high degree of intrinsic disorder and poorly defined structure ^2–4^. Several therapeutic strategies have indirectly targeted MYC or the MYC pathway. MYC is a nuclear transcription factor that drives the expression of genes required for cell growth, metabolism, and survival, and it is one of the most frequently amplified oncogenes in human malignancies ^2,5^. Through required heterodimerization with MAX, MYC binds E-box sequences to activate its targets. Because of MYC’s key role in tumor development, extensive efforts have focused on developing agents that either block MYC directly or interfere with pathways that control it ^2,3,6,7^. Such strategies include impeding MYC transcription with bromodomain and extraterminal (BET) family inhibitors or CDK7/9 inhibitors^8^, reducing MYC translation via mTORC1 or AKT/PI3K inhibitors^2,4^, preventing MYC-MAX dimer formation^2,4^, and inhibiting BRD4 with inhibitors such as JQ1 and GSK525762 that disrupt BRD4 binding at acetylated chromatin regions necessary for MYC transcription. Small-molecule microarray approaches have also uncovered MAX/MAX homostabilizers that sequester MAX away from MYC, thereby impairing MYC’s transcriptional function,^9^ and small-molecule inhibitors have been developed that disrupt the WDR5 and MYC protein interaction ^10–13^. Nevertheless, directly targeting MYC remains challenging due to its predominantly disordered structure and lack of prominent binding pockets, earning it the reputation of being “undruggable.”

Over the past several decades, covalent chemoproteomic approaches such as activity-based protein profiling (ABPP) have arisen as powerful approaches for uncovering unique, cryptic, allosteric, or shallow ligandable sites that can be accessed with covalently-acting small molecules through balancing reactivity with binding affinity ^14–25^. There have also been several successes in using covalent chemistry to target transcription factors, including against a palmitoylation site cysteine in the Hippo pathway transcription factor TEAD^26^, a cysteine within the Wnt pathway transcription factor CTNNB1^22^, and most recently, a cysteine in FOXA1 to rewire its transcriptional specificity ^23^. Recently, we have also uncovered a covalent ligand EN4 that targets an allosteric cysteine, C171 (or C186, depending on the MYC isoform and associated protein sequence) to cause destabilization of MYC and subsequent inhibition of MYC/MAX binding to DNA and MYC transcriptional activity in cells, leading to downregulation of MYC target genes, and anti-proliferative and anti-tumorigenic effects ^21^. Some of these covalent ligands, including EN4, have been shown to target intrinsically disordered cysteines within these proteins ^21^. However, whether there is an actionable binding pocket recognized by these molecules versus whether the binding occurs mainly through reactivity-driven binding has been less clear. One strategy to address this key question of molecular recognition versus reactivity is the use of stereochemically matched compound libraries composed of either enantio- and/or diastereomerically matched compounds that allow the direct comparisons between these stereoisomers to determine if stereoselective interactions are dominating the bioconjugation of the corresponding protein target with the chiral ligand. Previous studies from Cravatt, Vinogradova, and others have extensively exemplified such studies ^18,23,27–29^.

In this study, we explored the possibility of using sulfinyl aziridines as covalent small molecules that can stereoselectively modulate MYC levels and thus inhibit MYC transcriptional activity in cells. To this end, we synthesized a small library of covalent ligands bearing electrophilic aziridine-derived warheads of variable reactivity coupled with stereochemical pairing. Through this effort, we identified and optimized a covalent ligand that directly engages MYC in a stereoselective manner and, as a result, leads to MYC destabilization and proteasome-dependent degradation in cells.

## Results

### Synthesis and Screening of a Stereochemically Paired Covalent Ligand Library

We synthesized a small library of stereochemically well-defined aziridines bearing tunable sulfonyl and sulfinyl warhead activating groups surrounding a conserved spirocyclic oxindole core–a privileged motif in drug discovery **(Figure 1a)** ^30–35^. While oxindole-based sulfonyl aziridines have been shown to react with indoles, thiols, alcohols, and amines in a flask, their utility and limitations (toxicity, GSH stability, etc.) are unexplored in cellular, proteome-wide contexts as required for covalent drug discovery ^36,37^. This fact, together with the observation that their sulfinyl congeners are even less interrogated in similar contexts, prompted us to synthesize and interrogate these novel chiral probes in a cellular setting with the knowledge that aziridines are recognized electrophiles with a range of reactivities toward numerous amino-acid side-chains^36–39^.

**Figure 1.**
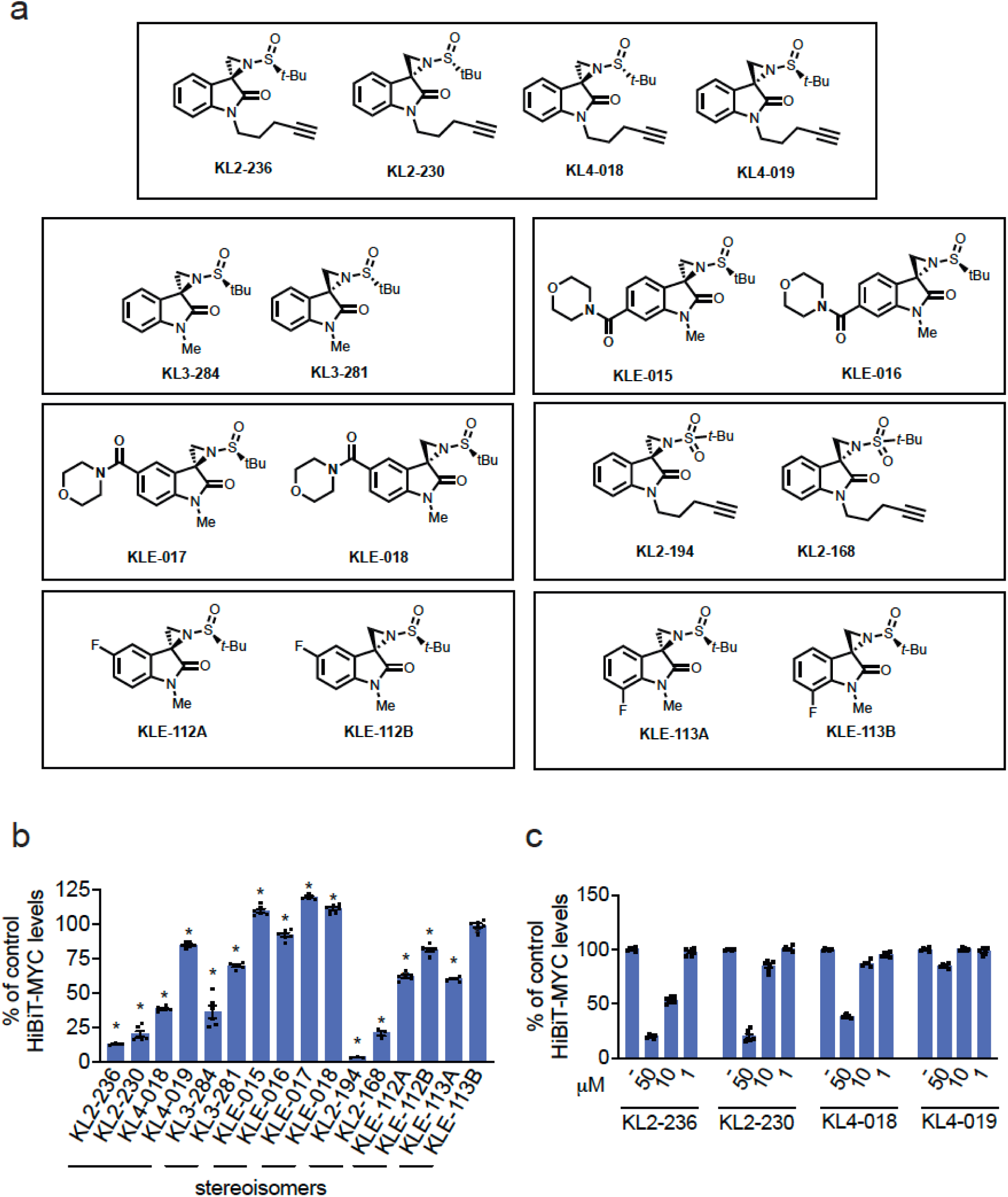
Screening a covalent ligand library for stereoselective MYC degradation. **(a)** Structures of members of a stereochemically-defined sulfinyl or sulfonyl oxindole aziridine covalent ligand library. **(b)** HiBiT-MYC HEK293-LgBiT cellular screen for MYC degraders. HiBiT-MYC HEK293-LgBiT cells were treated with DMSO vehicle or covalent compound (50 μM) for 24 h, and MYC levels were detected with the Nano-Glo HiBiT Lytic Detection System. **(c)** Dose-response of KL2-236 and its isomers in HiBiT-MYC cells. HiBiT-MYC HEK293-LgBiT cells were treated with DMSO vehicle, KL2-236, KL2-230, KL4-018, or KL4-019 for 24 h, and MYC levels were detected with the Nano-Glo HiBiT Lytic Detection System. Data in **(b,c)** are presented as individual replicate values and average ± sem percent of DMSO vehicle-treated controls. Significance is expressed as *p<0.05 compared to DMSO vehicle-treated controls.

We screened this bespoke covalent library in HEK293 cells expressing a HiBiT tag on the endogenous loci of MYC to identify molecules that lowered HiBiT-MYC levels **(Figure 1b).** We identified several molecules that significantly lowered HiBiT-MYC levels by >50 % – KL2-236, KL2-230, KL4-018, KL3-284, KL2-168, and KL2-194 **(Figure 1b).** While substituents appended to the oxindole ring generally decreased activity (i.e., KLE-017/KLE-018, KLE-015/KLE-016, KLE-112A/KLE-112B, KLE-113A/KLE-113B), modifications to the oxindole nitrogen were tolerated (see KL3-284). Interestingly, enantiomeric sulfonyl aziridine-containing compounds KL2-194 and KL2-168 showed only modest differences in their ability to attenuate HiBiT-MYC levels and were also significantly cytotoxic, thus limiting their further use **(Figure 1b, Figure S1a)**. In contrast, however, KL2-236, which possessed a sulfinyl activating group of attenuated reactivity, showed interesting patterns of stereoselective and dose-responsive HiBiT-MYC loss compared to its three stereoisomers **(Figure 1c)**. While KL2-236 is the most active stereoisomer in the series, its enantiomer KL2-230 also showed comparable activity. A diastereomer of these compounds, KL4-108 showed attenuated activity; however, its enantiomer KL4-019 was largely inactive. This pattern of stereoselective activity also held for oxindoles KL3-284 and KL3-281, KLE-112A and KLE-112B, and KLE-113A and KLE-113B. None of the other compounds screened, including the four stereoisomers KL2-236, KL2-230, KL4-018, or KL4-019, showed more than 25 % impairment in cell viability **(Figure S1a)**.

We next tested the chemoselectivity of the four sulfinyl aziridine stereoisomers alongside the sulfonyl aziridine KL2-194 with an artificial peptide bearing 15 of the 20 amino acids, including most nucleophilic amino acids, to assess amino acid reactivity preferences. We found that these compounds only reacted with cysteine and not with any other nucleophilic amino acid, including glutamic/aspartic acids, serines, threonines, lysines, or methionines in this experiment **(Figure S1b).** As expected, KL2-194, which possesses a more highly-oxidized sulfur atom, was more reactive than the four stereoisomers KL2-236, KL2-230, KL4-018, and KL4-019 **(Figure S1b-S1c)**. The higher reactivity of the sulfonyl aziridine KL2-194 compared to the sulfinyl aziridine counterparts is consistent with the higher degree of cytotoxicity observed with KL2-194. We also tested the *in vitro* stability of KL2-236 and KL4-019 with glutathione (GSH) and observed a comparable GSH half-life of 50.6 and 59.4 minutes, respectively. These values are longer than previously reported *in vitro* GSH stability of clinically approved covalent drugs such as afatinib and neratinib with GSH half-lives of ∼30 minutes ^40^. Overall, while our cellular data showed pronounced bioactivity differentiation between KL2-236 and its diastereomer KL4-019, these two compounds showed similar inherent reactivity with cysteine in a linear artificial peptide and with GSH. Moreover, our data also demonstrated that these sulfonyl aziridines were preferentially cysteine reactive.

Given that oxindole KL2-236 showed the best activity in reducing MYC without deleterious cytotoxicity and possessed a diastereomeric negative control compound (KL4-019), we prioritized downstream biological studies using this molecule. Confirming that HiBiT-MYC loss was through proteasome-mediated degradation rather than transcriptional downregulation, we observed significant attenuation of KL2-236-mediated HiBiT-MYC loss upon pre-treatment of cells with the proteasome inhibitor bortezomib **(Figure 2a)**. We further confirmed the proteasome-dependent degradation of endogenous MYC with KL2-236, but not with inactive stereoisomer KL4-019, in wild-type HEK293 cells **(Figure 2b-2c)**. We also demonstrated proteasome-mediated loss of MYC with KL2-236 treatment in more cancer-relevant PSN-1 pancreatic cancer cells with MYC amplification **(Figure 2d)**^41^.

**Figure 2.**
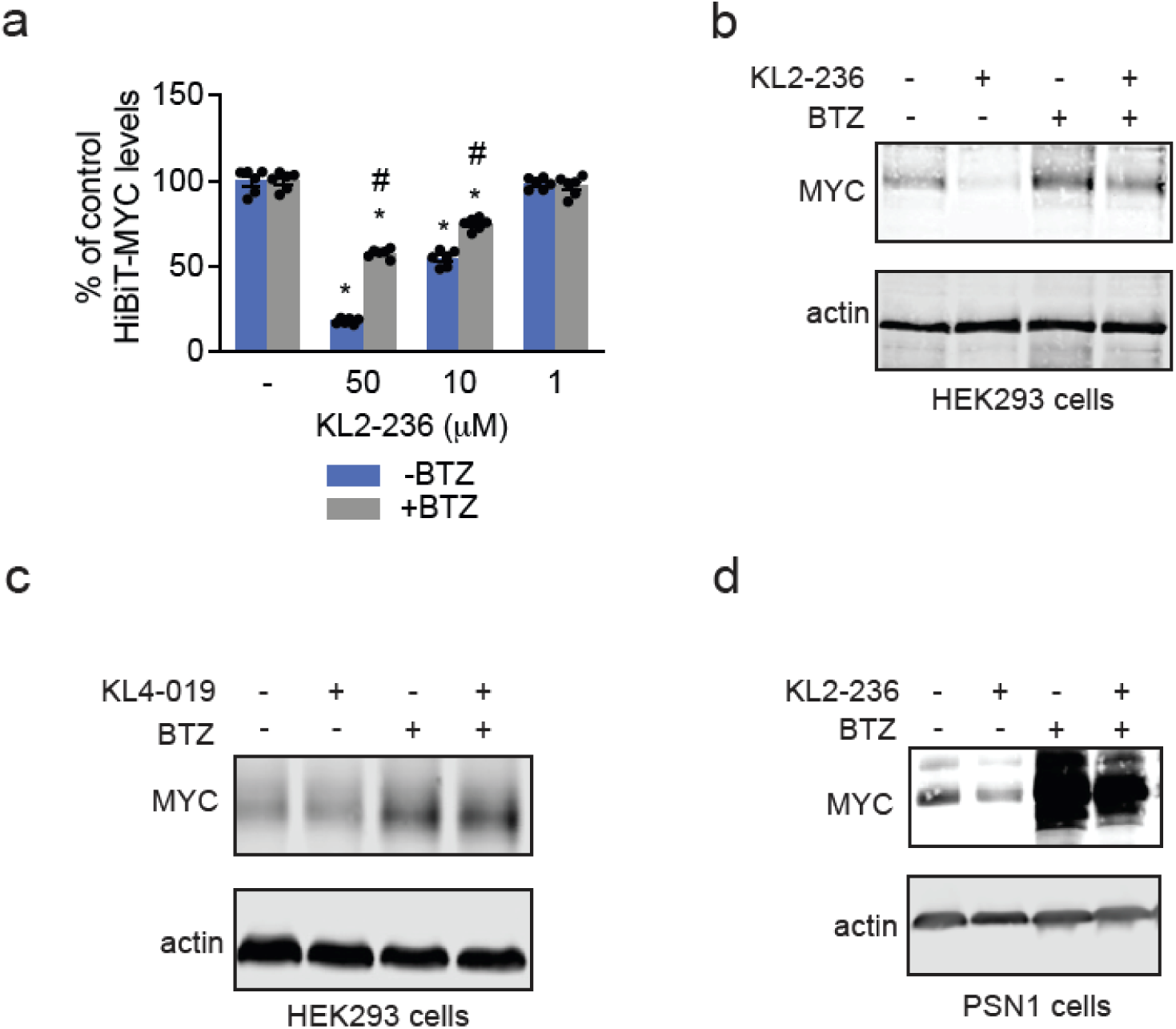
Characterization of KL2-236 and its Stereoisomers. **(a)** Proteasome-dependence of HiBiT-MYC loss. HiBiT-MYC HEK293-LgBiT cells were pre-treated with proteasome inhibitor bortezomib (500 nM) for 1 hr before treatment of DMSO vehicle or KL2-236 for 24 h, and MYC levels were detected by luminescence. **(b,c)** MYC degradation in HEK293 cells. HEK293 cells were treated with DMSO vehicle or bortezomib (500 nM) for 1 hr before treatment of DMSO vehicle or KL2-236 **(b)** or KL4-019 (50 μM) **(c)** for 24 h and MYC and loading control actin levels were assessed by Western blotting. **(d)** MYC degradation in PSN1 pancreatic cancer cells. PSN1 cells were treated with DMSO vehicle or bortezomib (500 nM) for 1 hr before treatment with DMSO vehicle or KL2-236 for 2 h, and MYC and loading control actin levels were assessed by Western blotting. Blots in **(b,c,d)** represent n=3 biologically independent replicates per group. Bar graph in **(a)** is individual replicate values and average ± sem. Significance in **(a)** is shown as *p<0.05 compared to respectively vehicle-treated controls and #p<0.05 compared to the KL2-236-treated group for each concentration.

### Characterizing KL2-236 Interactions with MYC

We next sought to determine whether KL2-236 directly interacted with MYC *in vitro* or in cancer cells. Given that KL2-236 had an alkyne handle for directly conjugating analytical handles through copper-catalyzed azide-alkyne cycloaddition (CuAAC), we exploited this feature to characterize its interactions using ABPP-based approaches. We first demonstrated direct covalent labeling of pure human full-length MYC in the MYC/MAX protein complex in a dose-responsive manner, visualized through CuAAC-mediated conjugation of a rhodamine fluorophore and gel-based ABPP **(Figure 3a)**. Strikingly, stereoselective labeling was also observed between KL2-236 and KL4-019, even at this pure MYC/MAX protein complex level **(Figure 3b)**.

**Figure 3.**
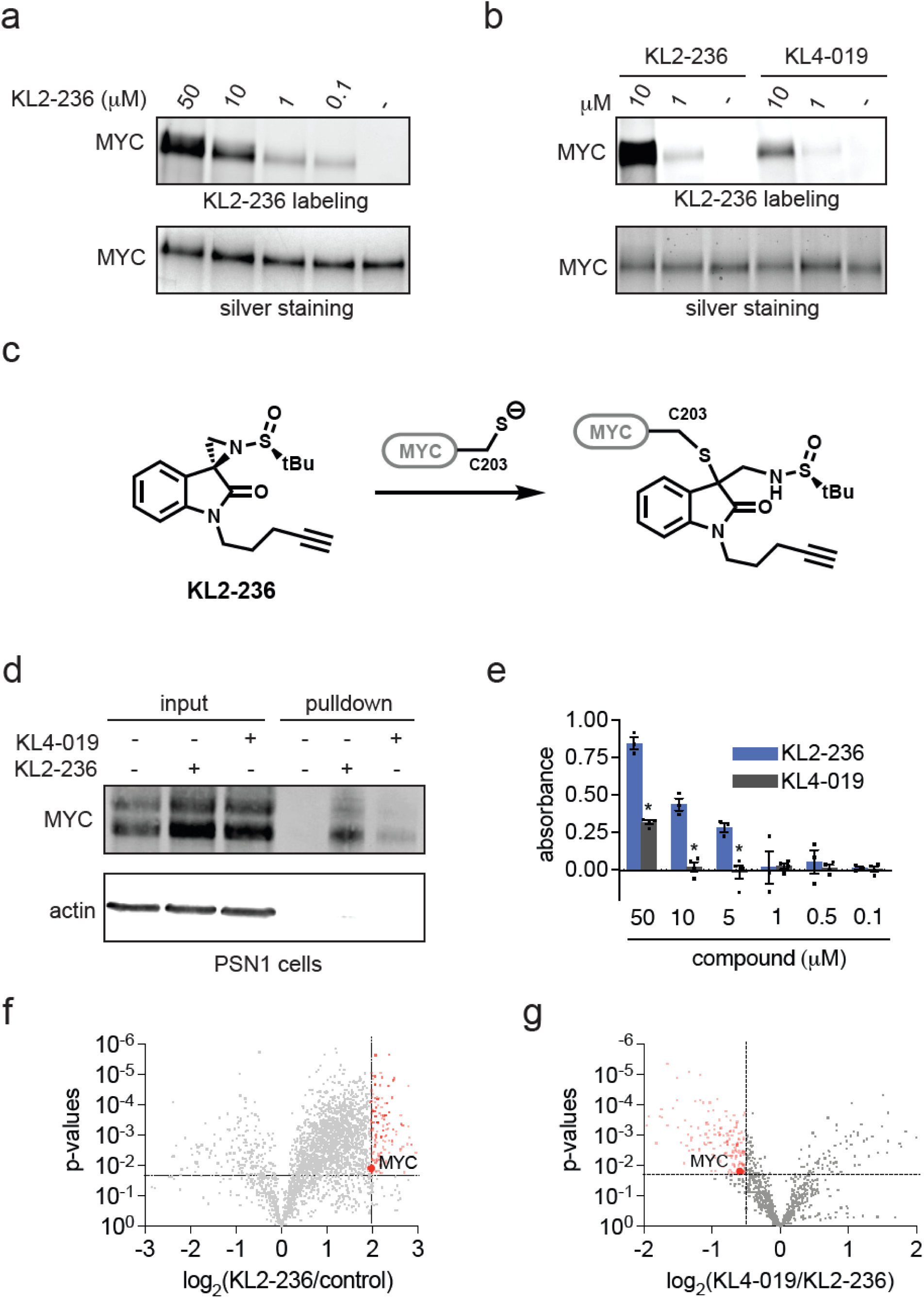
Characterization of KL2-236 Interactions with MYC. (a,b) Gel-based ABPP of KL2-236 and KL4-019 with pure human MYC/MAX protein. Pure MYC/MAX human protein complex was treated with DMSO vehicle, KL2-236, or KL4-019 for 1 h, after which probe-modified protein was subjected to CuAAC with an azide conjugated rhodamine, separated by SDS/PAGE and visualized by in-gel fluorescence. In **(b)**, KL2-236 and KL4-019 were treated at 50 μM. In **(a,b)**, gels were silver stained to account for MYC protein loading. **(c)** Reaction of KL2-236 with C203 of MYC from KL2-236 labeling studies wherein KL2-236 (50 μM, 1 hr) was incubated with pure human MYC/MAX protein (1 μM), followed by digestion of the complex with pepsin, and analysis of probe-modified peptides by LC-MS/MS as shown in **Figure S1.** Shown are the possible KL2-236 adducts on MYC C203. **(d)** KL2-236 versus KL4-019 engagement of MYC in PSN1 cells. PSN1 cells were pretreated with BTZ (500 nM) for 1 hour before treatment with DMSO vehicle, KL2-236 (50 μM) or KL4-019 (50 μM) for 2 h, after which probe-modified proteins were subjected to CuAAC with an azide-functionalized biotin, after which probe-modified proteins were enriched with avidin beads, eluted, and input and pulldown eluate was separated by SDS/PAGE and MYC and negative control protein actin were detected by Western blotting. **(e)** ELISA-ABPP assessment of KL2-236 versus KL4-019 cellular engagement of MYC in DLD-1 MYC-HiBiT expressing cells cells. DLD-1 MYC HiBiT expressing cells were treated with DMSO vehicle, KL2-236, or KL4-019 for 1 h after which probe-modified proteins were appended with a biotin handle by CuAAC, and HiBiT-MYC was captured and immobilized on plate using a HiBiT antibody and then compound-engaged HiBiT-MYC was detected by streptavidin HRP. **(f)** KL2-236 versus KL4-019 chemoproteomic profiling PSN1 cells. PSN1 cells were pretreated with BTZ (500 nM) for 1 hour before treatment with DMSO vehicle, KL2-236 (50 μM) or KL4-019 (50 μM) for 2 h, after which probe-modified proteins were subjected to CuAAC with an azide-functionalized biotin, after which probe-modified proteins were enriched with avidin beads, eluted, and input and pulldown eluate was tryptically digested and analyzed and quantified by LC-MS/MS. Gels, blots, and data in **(a,b,d,e,f,g)** represent n=3-4 biologically independent replicates per group. Data for **(f,g)** can be found in **Table S1.**

To understand where our molecule was binding in MYC, we assessed the KL2-236 modification site in the MYC/MAX pure protein complex through mass spectrometry (MS) analysis of KL2-236-modified MYC pepsin digests. The only modification found was on cysteine C203 **(Figure S1b; Figure 3c)**. While C203 was likely the primary site of modification, based on the MS/MS spectra and the chemoselectivity of KL2-236 for cysteine with our artificial peptide (**Figure S1b)**, we could also not rule out modification on the neighboring glutamic acid E202 **(Figure S2).** These data showed direct and covalent KL2-236 binding to MYC *in vitro*.

We next assessed direct MYC target engagement in living cells. Treating PSN-1 cells with KL2-236, appending a biotin enrichment handle through CuAAC, and avidin-enriching KL2-236-modified proteins, we observed significant and stereoselective MYC enrichment with KL2-236 treatment over both the less active diastereomer KL4-019 and vehicle-treated controls **(Figure 3d).** Using an orthogonal enzyme-linked immunosorbent assay (ELISA)-ABPP approach wherein we treated MYC-HiBiT expressing DLD-1 cells with KL2-236 or KL4-019, appended a biotin handle by CuAAC, captured and immobilized HiBiT-MYC using a HiBiT antibody, and reading out compound-engaged MYC-HiBiT by streptavidin-HRP, we also observed dose-responsive and significant stereoselective KL2-236 engagement of MYC-HiBiT over KL4-019 **(Figure 3e; Figure S3a)**. Through quantitative chemoproteomic profiling of KL2-236-enriched targets, we identified 85 proteins enriched by >4-fold with p<0.001, including MYC **(Figure 3f; Table S1)**. MYC and 161 other proteins were also enriched by KL2-236 more significantly than KL4-019 among 970 total proteins quantified **(Figure 3g, Table S1).** These data demonstrate direct stereoselective engagement of MYC with KL2-236 over isomer KL4-019 in cells.

To further demonstrate MYC engagement in cells by KL2-236, we performed a cellular thermal shift assay (CETSA) in PSN1 cells. Instead of observing the stabilization of MYC thermal stability, we found a significant destabilization of MYC thermal stability with KL2-236 treatment in PSN1 cells with a 6.4 °C shift in the temperature at which 50 % of MYC was stable (T_m_) from 51.8 °C to 45.4 °C **(Figure 4a-4b)**. This destabilization of MYC by KL2-236 was reminiscent of our previously discovered covalent destabilizing MYC degrader EN4 and our destabilizing CTNNB1 degrader NF764, indicating that KL2-236 was degrading MYC through a similar destabilizing degradation mechanism compared to traditional PROTACs and molecular glue degraders ^21,22,42,43^.

**Figure 4.**
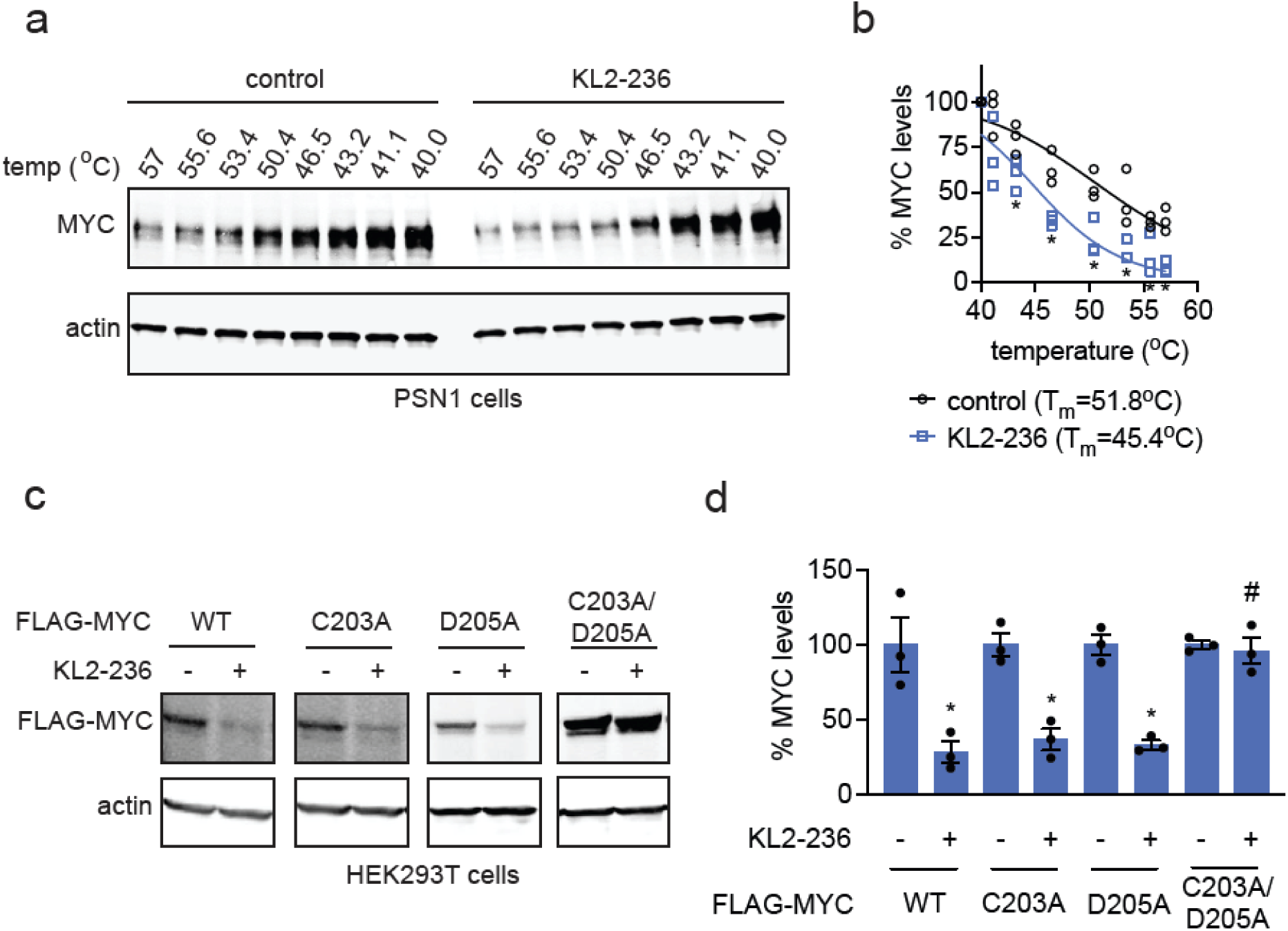
Understanding Mechanism Through Which KL2-236 Degrades MYC. (a,b) Cellular Thermal Shift Assay (CETSA) of KL2-236 on MYC and negative control actin in PSN1 cells. PSN1 cells were treated with DMSO vehicle or KL2-236 (50 μM) for 4 h. Cells were then heated at the designated temperatures, cells were harvested and lysed, insoluble proteins were pelleted, and soluble proteins were separated by SDS/PAGE and MYC and negative control actin levels were detected by Western blotting **(a)** and quantified **(b)**. **(c,d)** FLAG-MYC degradation with KL2-236 treatment. HEK293T cells stably expressing FLAG-MYC wild-type (WT), C203A, D205A, or C203A/D205A were treated with DMSO vehicle or KL2-236 (50 μM) for 24 h after which proteins were separated by SDS/PAGE and FLAG-MYC and loading control actin were detected (**c)** and quantified **(d)** by Western blotting. Blots in **(a,c)** represent n=3 biologically independent replicates per group. Data in **(b)** shows individual replicate values. Data in **(d)** shows individual replicate and average ± sem values. Significance in **(b,d)** expressed *p<0.05 compared to vehicle-treated controls and #p<0.05 compared to KL2-236 treated FLAG-MYC WT control groups in **(d)**.

To assess whether KL2-236 engagement of C203 was responsible for MYC degradation, we determined whether mutagenesis of C203 to alanine would attenuate MYC loss. Disappointingly, KL2-236 still degraded MYC C203A to the same extent as the wild-type protein in HEK293T cells **(Figure 4c-4d)**. We observed that C203 in MYC is neighbored by two acidic residues, E202 and D205. We postulated that one of these acidic residues could activate the C203 thiol, making it more nucleophilic and hyper-reactive and that KL2-236 could also potentially react with one of these acidic residues in the absence of C203. Alternatively, one of these acidic residues could be required for protonation of the chiral sulfoxide and subsequent positioning of the electrophile for covalent bond formation. Mutagenesis of E202 or D205 alone also did not attenuate MYC degradation from KL2-236 treatment **(Figure 4c-4d; Figure S2)**. However, we observed complete attenuation of MYC degradation with the MYC D205A/C203A double mutant **(Figure 4c-4d)**. We did not observe this rescue with the MYC E202A/C203A mutant **(Figure S3b)**. Overall, our results suggest that in addition to C203, D205 is involved in the reactivity and activity of KL2-236 with MYC.

### Impact of KL2-236 on MYC Transcriptional Activity

Thus far, we have shown that KL2-236 stereoselectively engages MYC *in vitro* and in cells at C203 and potentially through D205 to destabilize and degrade MYC in a proteasome-dependent manner. We next wanted to understand whether KL2-236 impairs MYC transcriptional activity and downregulates MYC target genes in cancer cells. KL2-236, but not the inactive isomer KL4-019, dose-responsively inhibited MYC luciferase reporter transcriptional activity in HEK293T cells **(Figure 5a).** KL2-236 treatment in PSN1 cancer cells significantly modulates>280 transcript levels **(Figure 5b; Table S2).** Consistent with KL2-236 action on inhibiting and degrading MYC, gene enrichment analysis of significantly modulated genes revealed the most significant pathway affected with the lowest normalized enrichment score as MYC targets, followed by IFNα response, E2F targets, oxidative phosphorylation, and IFNγ response **(Figure 5c-5d; Table S2)**. Our data collectively demonstrated that KL2-236 engages and degrades MYC, inhibits MYC transcriptional activity, and downregulates MYC target genes in cells.

**Figure 5.**
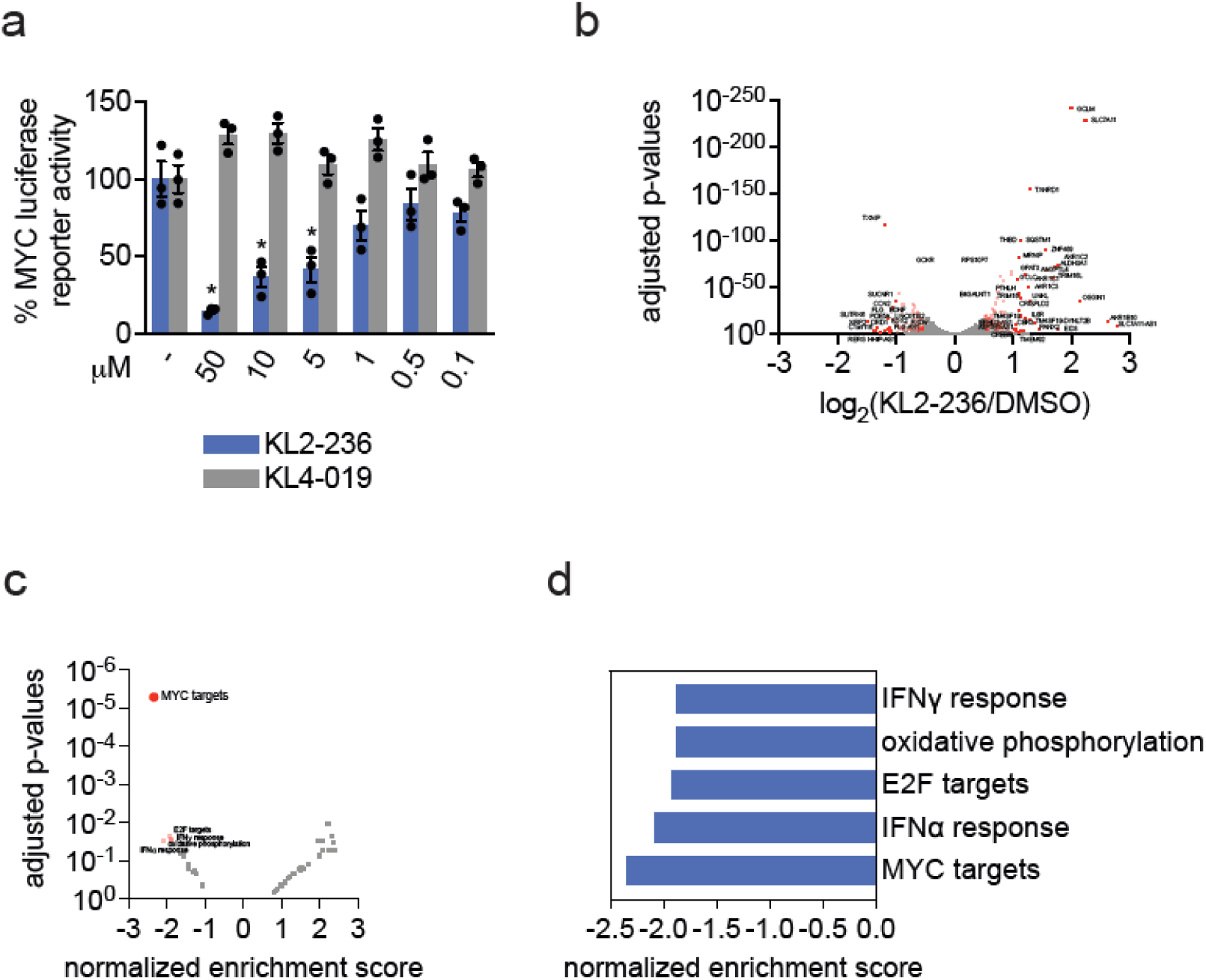
Functional Effects of KL2-236 on MYC Transcriptional Activity. **(a)** MYC luciferase reporter transcriptional activity in HEK293T cells. HEK293T cells expressing an MYC luciferase reporter were treated with DMSO vehicle, KL2-236, or KL4-019 for 24 h, after which MYC transcriptional activity was read out by luminescence. **(b)** RNA sequencing of KL2-236 in PSN1 cells. PSN1 cells were treated with DMSO vehicle or KL2-236 (50 μM) for 2 h. Resulting RNA from treated cells were sequenced and quantified. Shown in red are significantly altered transcripts (p<0.05, <log_2_ -0.5 or > log_2_ 0.5). Data are in **Table S2. (c,d)** Gene enrichment analysis shows significantly altered pathways and normalized enrichment scores, detailed in **Table S2**. Data in **(a-d)** are from n=3 biologically independent replicates per group. Data in **(a)** shows individual replicate and average ± sem values. Significance in **(a)** expressed *p<0.05 compared to vehicle-treated controls.

### Improving the Durability of MYC Degradation

While we showed compelling results with KL2-236 of a covalent molecule that stereoselectively engages MYC directly in cells to destabilize, degrade, and inhibit MYC, KL2-236 only shows acute loss of MYC in the first 2-6 hours of treatment with recovery of MYC protein levels by 12 h in PSN1 cancer cells where MYC turnover is rapid **(Figure 6a-6b)**. Indeed, this feature of MYC (rapid turnover) presents a known challenge for small molecule drug discovery efforts ^44^. Given the tolerability of modification to the oxindole nitrogen (**Figure 1**), we have found that an N-arylated analog of KL2-236, namely KL4-219A, shows more potent and durable degradation of MYC with continued MYC loss after 24 h of treatment in PSN1 cells **(Figure 6c-6e; Figure S4a)**. Like KL2-236, we also observed diminished HiBiT-MYC and parental MYC degradation and inhibition of MYC transcriptional reporter activity with the diastereomeric compound KL4-219B, compared to KL4-219A **(Figure 6d, 6f; Figure S4b-S4c).** The loss of MYC conferred by KL4-219A was attenuated upon pre-treatment of cells with a proteasome inhibitor, confirming proteasome-dependence of MYC loss **(Figure S4b-S4c).** Interestingly, these analogs, which induce more long-lasting degradation, have shorter *in vitro* GSH half-lives (19.3 and 26.3 minutes for KL4-219A and KL4-219B, respectively) as compared to KL2-236 and KL4-019. Thus, the sustained MYC degradation observed with KL4-219A may be related to other physicochemical parameters (permeability, potency, etc) as opposed to warhead stability. Previous studies have implicated that electrophiles may cause cell stress and stress granules that could complicate the interpretation of protein degradation ^45^. We show that neither KL2-236 nor KL4-219A causes stress granule formation at concentrations used in this study compared to the positive control sodium arsenite **(Figure S5)**. Chemoproteomic profiling showed 65 proteins that KL2-236 enriched that were significantly outcompeted by KL4-219A pre-treatment among >968 proteins quantified, demonstrating relatively selective cellular MYC engagement with KL4-219A **(Figure 6g, Table S3)**. Quantitative proteomic profiling of KL4-219A demonstrated remarkable proteome-wide selectivity for MYC degradation **(Figure 6h; Table S4)**. Overall, we put forth an optimized stereoselective and proteome-wide selective MYC degrader with KL4-219A.

**Figure 6.**
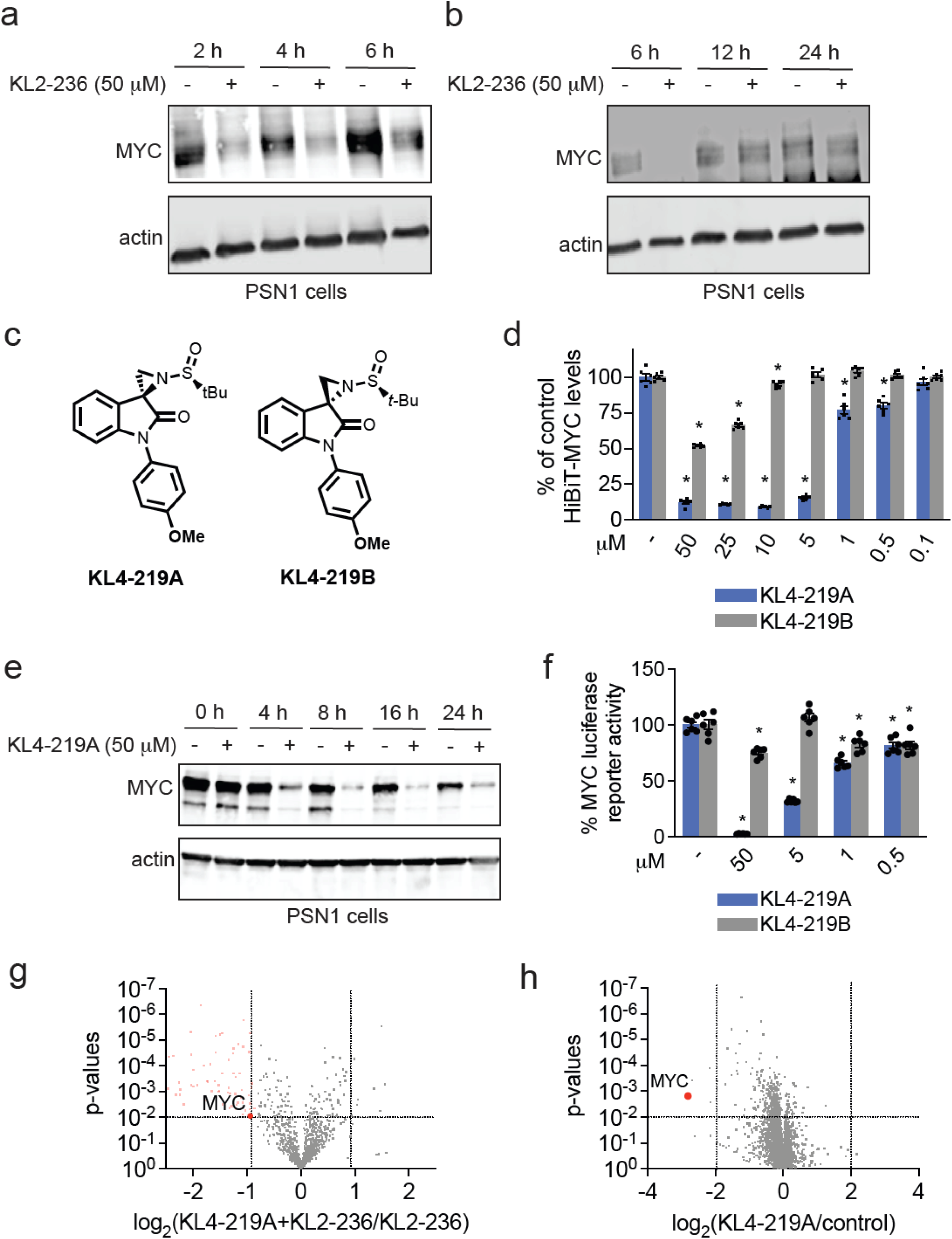
More Durable Analog of KL2-236: KL4-219A. (a,b) Time-course of KL2-236 degradation of MYC in PSN1 cells. PSN1 cells were treated with DMSO vehicle or KL2-236 (50 μM) for 24 h. Proteins were separated by SDS/PAGE and MYC and loading control actin levels were detected by Western blotting. **(c)** Structure of KL4-219A and diastereomer KL4-219B. **(d)** HiBiT-MYC HEK293-LgBiT cell dose-response with KL4-219A/B. HiBiT-MYC HEK293-LgBiT cells were treated with DMSO vehicle, KL4-219A, or KL4-219B (50 μM) for 24 h, and MYC levels were detected with the Nano-Glo HiBiT Lytic Detection System. **(e)** Time-course of KL4-219A-mediated MYC degradation in PSN1 cells. PSN1 cells were treated with a DMSO vehicle or KL4-219A (50 μM) at designated times, and MYC and loading control actin levels were assessed by Western blotting. **(f)** MYC luciferase reporter transcriptional activity in HEK293T cells. HEK293T cells expressing an MYC luciferase reporter were treated with DMSO vehicle, KL4-219A, KL4-219B for 24 h, after which MYC transcriptional activity was read out by luminescence. **(g)** Chemoproteomic profiling of KL4-219A. PSN1 cells were pre-treated with proteasome inhibitor bortezomib (500 nM) for 30 min prior to treatment with DMSO vehicle or KL4-219A (100 μM) for 1 h and then treatment with KL2-236 (25 μM) for 4 h, after which resulting cell lysates were subjected to CuAAC mediated appendage of a azide-functionalized biotin handle, probe-modified proteins were enriched, tryptically digested, and analyzed by TMT-based quantitative proteomics. Shown in red are proteins that were enriched by KL2-236 that were significantly (p<0.001) outcompeted by KL4-219A (log_2_ < -0.9). Data is shown in **Table S3. (h)** Quantitative proteomic profiling of KL4-219A in PSN1 cells. PSN1 cells were treated with DMSO vehicle or KL4-219A (50 μM) for 8h. Protein level changes were assessed by TMT-based quantitative proteomic methods. Shown in red is MYC. Data is shown in **Table S4.** Data or blots in **(a,b,e,f,g,h)** represent n=3-6 biologically independent replicates per group. The bar graphs in **(d,f)** show individual replicate values and average ± sem values from n=6 biologically independent replicates per group. Significance is shown as *p<0.05 compared to vehicle-treated controls.

## Discussion

Our findings address a longstanding challenge in chemical biology and oncology by demonstrating that MYC, a notoriously “undruggable” oncogenic transcription factor with extensive intrinsic disorder, can indeed be directly and stereoselectively targeted. Through a focused covalent library of stereochemically defined electrophiles, we discovered the hit compound KL2-236, which engages MYC at C203 with contribution from the neighboring D205 with distinct selectivity to its isomers to trigger proteasome-mediated degradation of MYC. While the nature of the interactions between these small molecules and MYC remains unclear at the molecular level, this study further highlights the power of chiral, covalent molecules to interrogate structurally undefined proteins. We also introduce the sulfinyl aziridine as a robust, protein-reactive electrophilic warhead with desirable stability, modulatable reactivity, and opportunities for direct at-warhead stereochemical tuning. The engagement of KL2-236 relative to KL4-019, even at the pure protein complex level, also further underscores the importance of expanding stereochemical space in covalent warhead design for targeting intrinsically disordered protein regions beyond that of just enantiomers, as a greater number of stereocenters may be required for observable compound differentiation ^18,23^. Spirocyclic oxindole sulfonyl aziridines react with nucleophiles under mild aqueous conditions owing to hydrogen-bond activation of the sulfonyl group with water^36,37^. This could suggest context-specific, two-point activation for the even less reactive sulfinyl aziridines reported herein whereby acidic protein residues position or activate the chiral sulfoxide before covalent bond formation occurs. Such a mechanism would be sensitive to local chiral environments found in dynamic structures within intrinsically disordered regions, in accordance with our stereochemical findings.

Because MYC is a central driver of growth and metabolism in various cancers, even modest decreases in MYC protein levels can have broad anticancer effects. By mapping the binding sites and confirming their importance in MYC degradation through mutagenesis studies, we uncovered a cooperative role for C203 and D205 in KL2-236-mediated MYC degradation, suggesting that residue adjacency and local electrostatics can significantly influence small-molecule reactivity within disordered protein regions. Consistent with this mechanism, KL2-236 not only degraded MYC but also demonstrably reduced its transcriptional activity— underscoring the central role of structural destabilization in achieving functional inhibition of an intrinsically disordered transcription factor.

Moreover, we improved upon the liability of KL2-236 in only achieving acute and transient MYC degradation through the development of KL4-219A, which exhibits enhanced degradation durability. Such iterative optimization exemplifies the power of rational medicinal chemistry approaches for improving the potency and temporal profile of covalent degraders. These differences in durability are likely driven by structural modifications that influence MYC engagement, covalent bond formation, proteasomal processing, and the stability and half-life of the molecule.

Further characterization of KL4-219A and more potent, selective, and metabolically stable analogs, particularly in physiologically relevant animal models, will be critical for translating these proof-of-concept findings into potential therapeutic interventions. Delineating how the surrounding protein environment in living systems affects compound reactivity and specificity will deepen our understanding of covalent ligand-protein interactions in intrinsically disordered regions. By demonstrating that intrinsically disordered proteins harbor exploitable cryptic sites, this work opens new vistas for targeting one of the most recalcitrant classes of oncogenic transcription factors.

## Materials and Methods

### Chemicals

Bortezomib (C835570) was purchased from Cayman. The Supporting Information describes the synthesis and characterization of the covalent library.

### Cell Culture

c-MYC HiBiT cells were commercially purchased from Promega (CS302396) in a HEK293-LgBiT background. HEK293T and PSN1 cells were obtained from UC Berkeley’s Biosciences Divisional Services Cell Culture Facility. LgBiT cells and HEK293T cells were cultured in DMEM (Corning 10-013-CV) with 10% Fetal Bovine Serum (FBS) (Corning 35-010-CV). PSN1 cells were cultured in RPMI1640 (Gibco 11875093) with 10% FBS. Cells were maintained at 37°C with 5% CO_2_.

### Screening and Testing of Covalent Ligands with HiBiT-MYC Cells

Covalent ligand screen and dose responses were conducted using the Nano-Glo HiBiT Lytic Detection System (Promega N3040) and CellTiter-Glo 2.0 Assay (G9242). c-MYC-HiBiT cells were seeded at 25,000 cells per well in a 96-well plate (Corning 3917) in 99µL of media and were left to adhere overnight. Cells were then pre-treated or treated with 1 µL of DMSO stock solutions containing varying concentrations of the compound under the conditions described. The Nano-Glo HiBiT Lytic Detection System and Cell Titer Glo 2.0 were used per the manufacturer’s instructions (1:1 media to reagent). Plates were incubated in the dark for 10 minutes before their luminescence readout on the Tecan Spark plate reader (30086376).

### Western Blotting

Cells were seeded overnight in 6 well plates (800,000 cells/well) or 6cm dishes (1,200,000 cells/dish). Media was aspirated, replaced with media containing DMSO or compound, and treated for the indicated length. Cells were then washed twice with PBS before harvesting by scraping. Cells were collected and pelleted, and PBS was aspirated. Pelleted cells were lysed using RIPA buffer (ThermoScientific 89900) containing protease inhibitor cocktail (Pierce A32955) and Benzonase (Millipore 70746-3 at 25-29 unit/mL) at 37°C for 10 minutes.

Sample protein concentrations were normalized using BCA Protein Assay (Pierce 23225) to run 20-50 µg per well depending on the desired protein target. Samples were boiled for 7 minutes at 95°C after the addition of 4x reducing Laemmli SDS sample loading buffer and run on precast 4-20% Criterion TGX gels (Bio-Rad) in 1x TGX buffer (Bio-Rad 1610734). Proteins were resolved by SDS/PAGE and transferred to 0.2 µm nitrocellulose membrane using the Bio-Rad Trans-Blot Turbo Transfer system (Bio-Rad, 1704150 and 1704271).

Membranes were blocked with 5% Bovine Serum Albumin (BSA) (bioWORLD 22070008-6) in Tris-buffered saline containing Tween 20 solution (TBST)(ThermoFisher J77500.K8) for 1 hour at room temperature. Blots were then probed with primary antibody solution overnight at 4°C. Antibodies used in this study were c-MYC (Abcam, AB32072), β-Actin (Cell Signaling Technology, 8H10D10 or Cell Signaling Technology, D6A8), and DDDDK tag (Cell Signaling Technology, 9A3) were diluted 1:1000 in 5% BSA in TBST. Membranes were then washed 3 times with TBST, followed by 1 hour of incubation with a secondary antibody solution in the dark at room temperature. Secondary antibodies used in this study include IRDye680RD Goat anti-Mouse IgG (LicorBio, 926-68070) and IRDye800CW Goat anti-Rabbit IgG (LicorBio 926-32211) diluted 1:10,000 in 5% BSA in TBST solution. Before imaging, membranes were washed an additional 3 times with TBST. Western blots were visualized using an Odyssey DLx Imager (LICORbio) and ChemiDoc MP Imaging System (Bio-Rad).

### Pure Protein Labeling with Covalent Ligands/Probes

MYC/MAX pure protein (0.5 µg/50 µL in PBS) was treated with DMSO vehicle or KL2-236 or KL4-019 at 37 °C for 30 minutes. A master mix of the click reagents was prepared such that each replicate would receive 0.25 µL of Rhodamine Azide (5mM stock in DMSO, Click Chemistry Tools, AZ109-5), 1 µL of copper (II) sulfate (50mM stock in water, Sigma-Aldrich, 203165), 3 µL of TBTA (1.7 mM in 4:1 tBuOH/DMSO, TCI Chemicals, T2993) and 1 µL of TCEP (50 mM in water freshly prepared, Thermo Scientific, 20491). Samples were vortexed and incubated at room temperature for 1 hour, after which 30 µL of 4x reducing Laemmli SDS sample loading buffer was added to each replicate and boiled at 95 °C for 10 minutes. Samples were cooled to room temperature, loaded into a precast 4-20% Criterion TGX gel (Bio-Rad), and run in 1x TGX buffer (Bio-Rad 1610734). Proteins were resolved by SDS/PAGE. Probe-labeled proteins were analyzed by in-gel fluorescence using the ChemiDoc MP Imaging system (Bio-Rad). Gels were silver-stained using the Pierce Silver Stain Kit (Thermo Scientific, 24612) per manufacturer instructions and imaged on the ChemiDoc MP Imaging System (Bio-Rad).

### Peptide Mapping Using Online Pepsinolysis LC-MS/MS

MYC(1-439)/MAX(2-160) (1 μM) protein was labeled with 50 μM KL2-236 in 20 mM HEPES, pH 7.0, 300 mM NaCl buffer for 1 hour at room temperature. The mixture as then directly injected on to LC-MS system.

Peptide mapping was performed using tandem online pepsin digestion and reversed-phase separation coupled to an Orbitrap Q Exactive mass spectrometer. Samples were injected onto an immobilized-pepsin column (Waters Enzymate BEH Pepsin, 2.1 mm × 30 mm) with resulting peptic peptides collected and separated by reversed-phase (C18) (Agilent ZORBAX Extend-C18, 1.0 mm × 150 mm, 3.5 μm particle size). The eluting peptides were then analyzed using data-dependent tandem MS acquisition (DDA). Samples (10 μL, 10 pmol) were injected at 0.25 mL/min in 0.1% formic acid in water for 4 min with resulting peptic peptides collected on the C18 trap cartridge. The pepsin column is then switched from the flow stream and peptides are eluted at 0.12 mL/min using a gradient of 8–40% B (0–14 min), followed by 70% B (2 min) and equilibration back to 2% B (Mobile Phase A = 0.1% formic acid, B = acetonitrile with 0.1% formic acid). Tandem MS spectra were acquired using the following DDA settings (Full MS Scan: 70,000 Resolution, AGC Target 3e6, *m*/*z* 370–1200 ddMS2 Settings; 17,500 resolution, AGC Target 1e5, max IT 50 ms, Isol window 2.0 *m*/*z*, NCE 30, Charge Exclusion >5, Min AGC Targe 8e4, and dynamic exclusion off).

### Expression and Purification of MYC/MAX

MYC (residues 1-439, isoform 1) without any purification tags was cloned into a pE-derived vector for expression in *E. coli*. MAX (residues 2-160) with an N-terminal STREP tag, followed by a TEV cleavage sit,e was cloned into a pE-derived vector for expression in *E. coli*.

MYC was expressed in BL21 cells as inclusion bodies by induction with 1 mM IPTG at an OD_600_ of 0.6-0.8 at 37 °C for 4 h. Cells were lysed with a high pressure homogenizer (Emulsiflex C3, Avestin) in lysis buffer (50 mM Tris/HCl pH 8.0, 5 mM DTT, 5 mM Benzamidine-HCl, 5 mM EDTA) and inclusion bodies were collected by centrifugation at 16,000 x *g* at 4 °C for 30 min. Inclusion bodies were washed once with lysis buffer and collected by centrifugation at 16,000 x *g* at 4 °C for 30 min. Inclusion bodies were washed a second time with water and collected by centrifugation at 16,000 x *g* at 4 °C for 30 min. Protein concentration in inclusion bodies was determined by HPLC analysis after dissolving in 6 M guanidine chloride. Inclusion bodies were dissolved in inclusion body buffer (25 mM HEPES/NaOH pH 7.6, 15 % glycerol, 300 mM NaCl, 0.1 mM EDTA, 1 mM DTT, 7 M Urea), clarified by centrifugation at 25,000 x *g* at 4 °C for 30 min and the supernatant was flash frozen in small aliquots at a final protein concentration of 5 mg/mL.

MAX was expressed in BL21 cells by induction with 0.15 mM IPTG at an OD_600_ of 2.2 at 18 °C for 16 h. Cells were lysed with a high pressure homogenizer (Emulsiflex C3, Avestin) in lysis buffer (50 mM Tris/HCl pH 7.5, 300 mM NaCl, one mM TCEP, one mM EDTA, 0.1 % Triton-X100) and the supernatant was cleared by centrifugation at 40,000 x *g* at 4 °C for 40 min. The supernatant was loaded on a prepacked StrepTrap HP column (Cytiva), washed with 5 CV of wash buffer (50 mM Tris/HCl pH 7.5, 300 mM NaCl) and bound protein was eluted with 5 CV elution buffer (50 mM Tris/HCl pH 7.5, 300 mM NaCl, 2.5 mM d-desthiobiotin).

The STREP-tag was removed by cleavage with TEV protease at RT for 4 h. MAX was further purified by anion exchange on a HiTrapQ column. The salt concentration in the TEV-cleaved sample was reduced to 25 mM NaCl by dilution with 50 mM Tris/HCl pH 8.3. The sample was loaded onto the HiTrapQ HP column, washed with 5 CV of HiTrapQ A buffer (50 mM Tris/HCl pH 8.3, 25 mM NaCl) and eluted with a gradient from 0-100 % HiTrapQ buffer B (50 mM Tris/HCl pH 8.3, 1000 mM NaCl) over 20 CV. Peak fractions were pooled, concentrated with a 10 kDa MWCO spin concentrator (Millipore), and further purified by size exclusion chromatography on a Superdex 200 16/60 PG column equilibrated in SEC buffer (50 mM Tris/HCl pH 7.5, 150 mM NaCl). Peak fractions were pooled and concentrated to >5 mg/mL with a 10 kDa MWCO spin concentrator (Millipore).

MYC/MAX was obtained by refolding MYC from inclusion bodies in presence of purified MAX. All steps were carried out at 4 °C. MYC/MAX were mixed at equal molar ratios and the sample was diluted to a final protein concentration of 0.25 mg/mL in refolding buffer (25 mM HEPES/NaOH pH 7.6, 15 % glycerol, 300 mM NaCl, 0.1 mM EDTA, 1 mM DTT, 6 M Urea). The complex was refolded by a stepwise dialysis (3 M Urea, 1 M Urea, 0.5 M Urea) with 10x buffer excess for each step over 1 h to 1.5 h. The final buffer exchange in 25 mM HEPES/NaOH pH 7.6, 15% glycerol, 300 mM NaCl, 0.1 mM EDTA, 1 mM DTT was performed over night with 30x buffer excess. The sample was further purified on a Heparin HP column, washed with 10 CV of Heparin HP A buffer (25 mM Tris/HCl, pH 8, 200 mM NaCl), 10% Heparin buffer B (25 mM Tris/HCl, pH 8, 2000 mM NaCl) and eluted with a gradient from 10-100 % Heparin buffer B (25 mM Tris, pH 8, 2000 mM NaCl) over 10 CV. The sample was concentrated using a 10 kDa MWCO spin concentrator (Millipore) and further purified by SEC on a Superdex75 16/60 column in SEC buffer (20 mM HEPES/NaOH pH7.0, 300 mM NaCl, 2 mM TCEP). Peak fractions containing a 1:1 molar ratio of MYC:MAX based on LC/MS analysis were pooled, flash frozen in liquid nitrogen and stored at -80 °C until use.

### Pulldown of MYC with Covalent Ligand/Probes for Western Blotting Detection

Pelleted cells were lysed by probe sonication on ice in PBS containing a protease inhibitor cocktail (Pierce A32955). Samples were centrifuged at 10,000 g at 4 °C for 10 minutes to remove cellular debris. Lysate protein concentrations were measured using BCA Protein Assay (Pierce 23225) and normalized to 5 mg/mL in 500 µL of PBS with protease inhibitor cocktail. A master mix of the click reagents was prepared such that each replicate would receive 10 µL of biotin picolyl azide (10 mM stock in DMSO, Sigma-Aldrich, 900912), 10 µL of copper (II) sulfate (50 mM stock in water, Sigma-Aldrich, 203165), 30 µL of TBTA (1.7 mM in 4:1 tBuOH/DMSO, TCI Chemicals, T2993) and 10 µL of TCEP (50mM in water freshly prepared, ThermoScientific, 20491). Samples were vortexed and incubated on a rotator at room temperature for 1 hour. Proteins were pelleted by centrifugation (6500 g) at room temperature for 5 minutes. The supernatant was removed, 500 µL of cold methanol was added, and samples underwent three cold methanol washes with centrifugation to pellet proteins and sonication for resuspension. Pelleted proteins were redissolved in 200 µL of PBS with 1.2% SDS (w/v) by sonication. Samples were heated to 90°C for 5 minutes, and 5µL of the sample was removed and saved for input (diluted to 60 µL and 20 µL of 4x reducing Laemmli SDS sample loading buffer). 55 µL (per sample) of streptavidin-agarose beads (ThermoFisher, 20353) were washed with PBS using a Micro-Bio Spin Column (Bio-Rad, 7326204) and vacuum manifold. 500 µL of PBS was added to the dissolved proteins, and then the washed beads were added to each sample using two washes of 250 µL of PBS. Samples were incubated on a rotator at 4°C overnight. Samples were then put in a 37°C bath to redissolve SDS for 5 minutes, and then beads were pelleted by centrifugation at 1400 g for 5 min. The supernatant was removed, and beads were washed with 0.2% SDS in PBS (w/v) for 10 minutes on a rotator at room temperature. The supernatant was removed, and pelleted beads were moved to Micro-Bio Spin Columns using two washes of PBS. Beads were washed on a vacuum manifold three times with PBS and then three times with water. The beads were then moved to screw cap Eppendorf tubes with PBS, centrifuged, and the supernatant removed. 30 µL of 1x reducing Laemmli SDS sample loading buffer was added to each sample and then boiled at 95 °C for 15 minutes to elute proteins from beads. Samples and corresponding inputs were run on precast 4-20% Criterion TGX gels (Bio-Rad) in 1x TGX buffer (Bio-Rad 1610734). Proteins were resolved by SDS/PAGE and transferred to 0.2 µm nitrocellulose membrane using the Bio-Rad Trans-Blot Turbo Transfer system (Bio-Rad, 1704150 and 1704271). Membranes were blocked with 5% Bovine Serum Albumin (BSA) (bioWORLD 22070008-6) in Tris-buffered saline containing Tween 20 solution (TBST)(ThermoFisher J77500.K8) for 1 hour at room temperature. Blots were then probed with primary antibody solution overnight at 4 °C. Antibodies used in this study were c-MYC (Abcam, AB32072) and B-Actin (Cell Signaling Technology, 8H10D10), were diluted 1:1000 in 5% BSA in TBST. Membranes were then washed 3 times with TBST, followed by 1 hour of incubation with a secondary antibody solution in the dark at room temperature. Secondary antibodies used in this study include IRDye680RD Goat anti-Mouse IgG (LicorBio, 926-68070) and IRDye800CW Goat anti-Rabbit IgG (LicorBio 926-32211) diluted 1:10,000 in 5% BSA in TBST solution. Before imaging, membranes were washed an additional 3 times with TBST. Western blots were visualized using an Odyssey DLx Imager (LICORbio).

### Pulldown TMT Proteomics Experiment with Covalent Ligand/Probes

Cells were seeded at 15 million cell/dish in 15 cm dishes (1 dish per replicate). All cells were pre-treated with 500 nM of Bortezomib for 1 hour. Cells were then treated with compound at 50µM final concentration and incubated for an additional 4 hours. Cells were then washed twice with PBS, scraped, and collected. Cells were pelleted by centrifugation, and the supernatant was removed. Cells were lysed by probe sonication on ice in PBS containing a protease inhibitor cocktail (Pierce A32955). Samples were centrifuged at 10,000 g at 4 °C for 10 minutes to remove cellular debris. Lysate protein concentrations were measured using BCA Protein Assay (Pierce 23225) and normalized. A master mix of the click reagents was prepared such that each replicate would receive 10 µL of biotin picolyl azide (10 mM stock in DMSO, Sigma-Aldrich, 900912), 10 µL of copper (II) sulfate (50 mM stock in water, Sigma-Aldrich, 203165), 30 µL of TBTA (1.7 mM in 4:1 tBuOH/DMSO, TCI Chemicals, T2993) and 10 µL of TCEP (50 mM in water freshly prepared, ThermoScientific, 20491). Samples were vortexed and incubated on a rotator at room temperature for 1 hour.

To each sample, 100% acetonitrile was added to the final concentration of 80% and mixed by inversion to precipitate proteins. Proteins were pelleted by centrifugation at 21,000 g at 4 °C for 10 minutes. The supernatant was removed, 500 µL of cold methanol was added, and samples underwent three cold methanol washes with centrifugation to pellet proteins and sonication for resuspension. After the final wash, the pellets were left to air dry for 10 minutes. Proteins were redissolved in 1 mL of PBS with 1.2% SDS (w/v) by sonication. Samples were then heated to 90°C for 5 minutes, and 5 mL of PBS was transferred to 15 mL tubes. 170 µL of streptavidin-agarose beads were added to each sample and incubated at 4°C overnight with constant rotation. Samples were then put in a 37°C bath to redissolve SDS for 5 minutes, and then beads were pelleted by centrifugation at 1400 g for 5 min. The supernatant was removed, and beads were washed with 0.2% SDS in PBS (w/v) for 10 minutes on a rotator at room temperature. The supernatant was removed, and pelleted beads were moved to Micro-Bio Spin Columns using two washes of PBS. Beads were washed on a vacuum manifold four times with PBS and then four times with water. The beads were then moved to screw cap Eppendorf tubes with two 250 µL washes of 6 M Urea in PBS. To each sample, 25 µL of DTT (30 mg/mL in water) was added and incubated at 65 °C for 20 minutes, gently mixing every 5 minutes. Samples were cooled to room temperature, and then 25 µL of iodoacetamide (400 mM in water) was added and incubated at 37 °C for 30 minutes with constant agitation. Samples were then diluted with PBS, centrifuged, the supernatant removed, rewashed with PBS, and then washed once with 500 µL of 50 mM TEAB (ThermoScientific 90114), and the supernatant removed. Beads were resuspended in 100 µL of 50 mM TEAB, and 4 µL of sequencing grade trypsin (0.5 mg/mL, Promega, V5111) was added to each sample. Samples were digested at 37°C overnight with constant agitation. Samples (beads and liquid) were then moved to a Micro-Bio Spin Column and centrifuged at 2000 g for 2 min at RT to collect flow through. Quantification of peptides was done using the Thermo Scientific Pierce Quantitative Colorimetric Peptide Assay (ThermoScientific, 23275). 100 µL of each sample was for TMT labeling. TMT labels (ThermoScientific, 90061, or A58332) were equilibrated with anhydrous acetonitrile, and 20 µL of each tag was added to a corresponding replicate. The reaction was incubated at room temperature for 1 hour on a rotating mixer and then quenched with 5% hydroxylamine for 15 minutes at room temperature. Peptides were dried down by Vacufuge (Eppendorf, 022820168) at 30 °C until dry. Samples were then resuspended in 50 µL of 0.1% Trifluoroacetic acid (TFA) in water and combined. Using the previously recorded concentrations, 100 µg of peptides were then fractionated according to the manufacturer’s protocol using the Thermo Fisher High pH Reverse Phase Fractionation Kit (ThermoScientific, 84868). Following fractionation, samples were dried down using Vacufuge at 30°C and then resuspended in 25 µL of 0.1% formic acid in water by vortexing and bath sonication. Samples were centrifuged at 20,000 g for 10 minutes at 4 °C and the supernatant were transferred into an LCMS vial with a glass insert (ThermoScientific, 6PME03C1SP) and capped. Samples were analyzed by LC-MS/MS.

### ELISA-ABPP

For capture plate preparation, the HiBiT capture antibody (Promega N7200) was diluted down to 2.5 µg/mL in ELISA coating buffer A (Invitrogen CB07100) and 100 µL were dispensed into each well of a Nunc MaxiSorp flat bottom 96-well plate (Thermo 473768) and incubated overnight at 4C. The following day, the plate was washed once with 200µL PBST (Boston BioProducts IBB-171) and blocked with 200 µL SuperBlock (Thermo 37515) for 1 hour at room temperature. Blocking buffer was removed, plate was washed three times with 200 µL PBST, and coated plate was used immediately for ELISA-ABPP.

For the ELISA-ABPP method, 96-well cell culture plates (Corning 3596) were coated with 100 µL of 0.1 mg/mL poly-D-lysine (Sigma Aldrich P7280-5MG) in PBS for 3 hours at room temperature followed by two 200 µL washes with PBS. DLD1 cells expressing HiBiT-MYC were seeded at 75,000 cells per well in complete media (RPMI, 10% FBS, 1% Penn/Strep, 10 µg/mL blasticidin) and incubated overnight (37 °C, 5% CO_2_) to allow cells to attach. The next day, the media was removed and replaced with 75 µL complete media followed with 25 µL of a 4x stock of the oxindoles in complete media and incubated for 1 hour at 37C, 5%CO_2_. After incubation, the media was removed, and cells were washed once with 200 µL warm cell imaging solution (Thermo A59688DJ) followed with addition of 100 µL lysis buffer consisting of 1% n-dodecyl-β-D-maltopyranoside (Anatrace D310-25GM) in cell imaging solution containing cOmplete EDTA-free protease inhibitors (Sigma Aldrich 4693132001) and lysed for 1 hour at room temperature with constant mixing. During lysis, reporter biotin click reagents were prepared by dissolving 100 mg THPTA (Vector 1010-5G) into 5 mL of water, 100 mg ascorbate (Sigma Aldrich 11140-5G) into 5mL water, 20mg copper sulfate (Sigma Aldrich 451657-10G) into 5 mL water, and 125µL 10 mM biotin picolyl azide (Vector 1167-100MG) into 5 mL of water. The individual click components were combined and 50 µL of the click mix was added to each well containing protein lysate and incubated for 1 hour at room temperature. The click reaction was quenched with the addition of 12.5 µL 0.5 M EDTA (Sigma Aldrich 324506-100ML) and centrifuged at 3000xg for 15 minutes at 4 °C. After centrifugation, 130 µL of clarified lysate was transferred to “Capture Plate” and incubated overnight at 4C with gentle mixing. After incubation, capture plates were washed five times with 200 µL PBST. Following final wash, 50 µL of streptavidin-HRP (CST 3999S) was added at a 1:1000 dilution and incubated for 1 hour at room temperature. After incubation, capture plate were washed five times with 200µL PBST followed with addition of 135µL of TMB substrate (Thermo N301) with a 5 minute incubation at room temperature with gentle mixing. Following incubation, the reaction was quenched with addition of 50 µL TMB stop solution (Thermo N600) and strong shaking for a few seconds followed by reading absorbance at 450 nm on PHERAstar FS (BMG Labtech).

### Generating Cells Expressing FLAG-Wild-Type or Mutant MYC

Human MYC constructs were customized and purchased from GenScript. The customized construct was in a pGenLenti backbone and MYC had a C-terminal DYKDDDDK (FLAG) tag. All mutant constructs were ordered from GenScript, which maintained a pGenLenti backbone and C-terminal DYKDDDDK tag.

These constructs were then used to generate stable FLAG-MYC-expressing cells using lentiviral infection. Before transfection, HEK293T cells were conditioned in complete media containing heat-inactivated FBS. For lentivirus production, FLAG-tagged wild type c-MYC or FLAG-tagged MYC mutant plasmids (GenScript), psPAX2 (Addgene, 12260) and pMD2.G (Addgene, 12259) were transfected into HEK293T cells using Lipofectamine 2000 (ThermoFisher, 11668027) in Opti-MEM (Gibco, 31985062). The virus-containing medium was collected and filtered (0.45 µm PES) after 48 hours. The virus was then used to infect new HEK293T with a 1:1000 dilution of Polybrene (Sigma-Aldrich, TR-1003-G). After 48 hours, infected cells were selected with puromycin (Abcam, ab141453) (1.5 µg/mL for HEK293T cells. After 4 days of the selection, cells were recovered by replacing puromycin-containing media with fresh media.

### Luciferase Reporter assay

MYC luciferase reporter assay was purchased from Qiagen (CCS-012L) and performed according to the manufacturer’s protocol with minor alterations. Before transfection, HEK293T cells were seeded into a 96-well plate (Corning 3917) at 30,000 cells/well in 100 µL of media. Negative control, positive control, or MYC luciferase reporter construct were diluted in Opti-MEM medium (1 µL DNA into 25 µL Opti-MEM per well). Attractene (Qiagen, 301005) was diluted in Opti-MEM (1 µL Attractene into 25 µL Opti-MEM per well). DNA and diluted Attractene were gently mixed in a 1:1 ratio and then incubated at room temperature for 20 minutes undisturbed. 50 µL of DNA-Attractene mixture was added to each well in triplicate. 24 hours post-transfection, media was carefully aspirated from each well, and 75 µL of media containing either DMSO or compound was added. After 24 hours of compound treatment, Dual-Glo luciferase assay (Promega, E2920) was used according to the manufacturer’s protocol for luminescent readout. Firefly and Renilla luminescence were read on a Tecan Spark plate reader. Background luminescence levels were subtracted using a blank control, and then Firefly: Renilla was calculated for each well.

### Cellular Thermal Shift Assay (CETSA)

PSN1 cells were harvested by scraping from a 10 cm dish after 1 hour at 50 µM compound treatment. Cells were resuspended in PBS containing protease inhibitor cocktail and 50 µM compound and then aliquoted into eight 0.2 mL PCR strips with 100 µL per tube. PCR strips were designated a temperature via a gradient program on Bio-Rad’s T100 Thermal cycler (Bio-Rad, 1861096). Samples were heated at their respective temperatures (57 °C, 55.6 °C, 53.4 °C, 50.4 °C, 46.5 °C, 43.2 °C, 41.1 °C, 40 °C) for 3 minutes and then at 25 °C for 3 minutes. Immediately following, cells were snap-lysed using liquid nitrogen (3 freeze-thaw cycles). Cell debris, along with any precipitated and aggregated proteins, were removed by centrifugation at 20,000 g for 20 minutes at 4 °C. 80 µL of supernatant was transferred to new PCR strips, 26.6 µL of 4x reducing Laemmli SDS sample loading buffer was added to each sample and boiled at 95°C for 10 minutes. 12 µL of each sample with buffer were loaded per well and run on precast 4-20% Criterion TGX gels (Bio-Rad) in 1x TGX buffer (Bio-Rad 1610734). Proteins were resolved by SDS/PAGE and transferred to 0.2µm nitrocellulose membrane using the Bio-Rad Trans-Blot Turbo Transfer System (Bio-Rad, 1704150 and 1704271). Membranes were blocked with 5% Bovine Serum Albumin (BSA) (bioWORLD 22070008-6) in Tris-buffered saline containing Tween 20 solution (TBST) (Thermo Fisher J77500.K8) for 1 hour at room temperature. Blots were then probed with primary antibody solution overnight at 4 °C. Antibodies used in this study were c-MYC (Abcam, AB32072) βανδ -Actin (Cell Signaling Technology, 8H10D10), diluted 1:1000 in 5% BSA in TBST. Membranes were washed 3 times with TBST, followed by 1 hour of incubation with a secondary antibody solution in the dark at room temperature. Secondary antibodies used in this study include IRDye680RD Goat anti-Mouse IgG (LicorBio, 926-68070) and IRDye800CW Goat anti-Rabbit IgG (LicorBio 926-32211) diluted 1:10,000 in 5% BSA in TBST solution. Before imaging, membranes were washed an additional 3 times with TBST. Western blots were visualized using an Odyssey DLx Imager (LICORbio).

### RNA Extraction

PSN-1 cells were seeded at 1,200,000 cells/dish in a 6 cm dish and allowed to adhere overnight. Aspirate media, replace it with media containing DMSO or compound, and treat it for 2 hours. After treatment, cells were washed twice with PBS before harvesting by scraping. Cells were collected into a 1.7 mL Eppendorf tube and centrifuged to collect pellet cells. PBS supernatant was aspirated, and the cell pellet was further processed. The Monarch Total RNA Miniprep kit (New England Biolabs, T2010S) was used to harvest total RNA and was followed according to the manufacturer’s protocols.

### RNA Sequencing

Library preparation and sequencing was performed by the QB3-Berkeley Genomics core labs (cite). Total RNA quality, as well as poly-dT enriched mRNA quality, were assessed on an Agilent 2100 Bioanalyzer. Libraries were prepared using the KAPA mRNA Hyper Prep kit (Roche KK8581). Truncated universal stub adapters were ligated to cDNA fragments, which were then extended using 10 cycles of PCR using unique dual indexing primers into full length Illumina libraries. Library quality was checked on an AATI (now Agilent) Fragment Analyzer and transferred to the Vincent J. Coates Genomics Sequencing Laboratory (GSL), another QB3-Berkeley Core Research Facility at UC Berkeley.

### Analysis of RNA Seq Data

RNA libraries were sequenced, generating 30-40 million 150bp paired end reads per sample (average 36 million reads). Fastq files were processed using the Exon Quantification Pipeline (v2.5)^46^, in which reads were aligned to the human genome (GRCh38) using STAR (version 2.7.3a)^47^ with an average of 32.9 million reads mapped per sample (89-92%), and gene counts were generated. Prior to differential expression analysis, DESeq2 (v1.44.0)^48^ normalized gene read counts (vsn; v3.72.0)^49^ were visualized, and one outlier sample was removed from subsequent analysis due to high read count variability. When downstream analysis was repeated with this outlier included, results were consistent with the outlier-removed analysis, and conclusions were not changed. Differential expression analysis was performed using DESeq2 comparing KL2-236 versus DMSO. Genes with an adjusted p-value of less than 0.05 were considered to be significantly differentially expressed. To examine gene set enrichments, gene lists were ranked using -log10(p-value)*sign(log2 Fold Change), and MSigDB Hallmark gene sets (v7.5.1)^50^ were tested for gene set enrichment using fGSEA (v1.30.0)^51^. Results were visualized using ggplot2 (v3.5.1)^52^.

### Assessment of Stress Granules

Cells were seeded in 4-well glass bottom cell culture dishes at 100,000 cells/well (Greiner Bio, 627870). The treated cells were fixed with 4% paraformaldehyde in PBS (Cell Signaling Technology, 47746P) for 15 minutes at room temperature and then blocked with Blocking Buffer (Cell Signaling Technology, 12411) for one hour at room temperature. Samples were further incubated with primary antibodies in Antibody Dilution Buffer (Cell Signaling Technology, 12378S) either overnight at 4°C or at room temperature for 1 hour. Following primary antibody incubation, cells were washed 3 times with PBS and then incubated in secondary antibody in Antibody Dilution Buffer for 1 hour at room temperature. Antibodies used in this experiment include G3BP1 (BD Biosciences, 611126), Phalloidin-iFluor 594 Conjugate (Abcam, ab176757) and Anti-mouse IgG (H+L), F(ab’)2 Fragment (Alexa Fluor 488 Conjugate) (Cell Signaling Technology, 4408S). Upon completion of immunostaining, a few drops of ProLong Gold Antifade reagent with DAPI (Invitrogen P36941) were added to each well. Images were captured using a Zeiss LSM880 FCS microscope with a 63x oil objective.

## Acknowledgment

We thank the members of the Nomura Research Group and Novartis BioMedical Research for critically reading the manuscript. This work was also supported by Novartis Biomedical Research, the National Science Foundation Molecular Foundations for Biotechnology (MFB) grant (2127788), the UC Berkeley Molecular Therapeutics Initiative (MTI), Bakar Fellows Award, the Mark Foundation for Cancer Research ASPIRE Award, and the National Institutes of Health (R35CA263814, R01CA240981, UM1CA29410). We also thank Hasan, Lund, and the UC Berkeley NMR facility in the College of Chemistry (CoC-NMR) for spectroscopic assistance. Instruments in the College of Chemistry NMR facility are partly supported by NIH S10OD024998. K. L. acknowledges the National Science Foundation for a pre-doctoral graduate fellowship. RNA Sequencing was performed at the QB3 Genomics Facility at UC Berkeley, Berkeley, CA RRID:SCR_022170 and was supported by NIH S10 OD018174 Instrumentation Grant. We also recognize the RCNR Biological Imaging Facility at the University of California, Berkeley and Denise Schichnes for their microscopy facilities, expertise and support.

## Author Contributions

DKN, TJM, HTR, KL conceived of the project. HTR, KL, ELL, BC, SMB, FJG, DCB, SHH, DD, DKN performed experiments, analyzed data, and interpreted results. HTR, KL, SMB, DD, JMM, DCB, FJG, JMM, MS, TJM, and DKN wrote the paper.

