## Supporting Information for "Sulfinyl Aziridines as Stereoselective Covalent Destabilizing Degraders of the Oncogenic Transcription Factor MYC"

* Authors contributed equally to the work

**Supporting Table Legends**

**Table S1. Chemoproteomic Profiling of KL2-236 and KL4-019.** Chemoproteomic profiling of KL2-236 versus KL4-019 in PSN1 cells. PSN1 cells were pretreated with BTZ for 1 hr before treatment with DMSO vehicle, KL2-236 (50 μM) or KL4-019 (50 μM) for 4 h, after which probe-modified proteins were subjected to CuAAC with an azide-functionalized biotin, after which probe-modified proteins were enriched with avidin beads, eluted, pulldown eluate was tryptically digested and analyzed and quantified by LC-MS/MS. Data are from n=3 biologically independent replicates per group.

**Table S2. RNA Sequencing of KL2-236 Treatment in PSN1 Cells.** RNA sequencing of KL2-236 in PSN1 cells. PSN1 cells were treated with DMSO vehicle or KL2-236 (50 μM) for 2 h. Resulting RNA from treated cells were sequenced and quantified. Gene enrichment analysis shows significantly altered pathways and normalized enrichment scores.

**Table S3. Chemoproteomic Profiling of KL4-219A.** PSN1 cells were pre-treated with proteasome inhibitor bortezomib (500 nM) for 30 min prior to treatment with DMSO vehicle or KL4-219A (100 μM) for 1 h and then treatment with KL2-236 (25 μM) for 4 h, after which resulting cell lysates were subjected to CuAAC mediated appendage of an azide-functionalized biotin handle, probe-modified proteins were enriched, tryptically digested, and analyzed by TMT-based quantitative proteomics.

**Table S4. Quantitative Proteomic Profiling of KL4-219A.** Quantitative proteomic profiling of KL4-219A in PSN1 cells. PSN1 cells were treated with DMSO vehicle or KL4-219A (50 μM) for 8h. Protein level changes were assessed by TMT-based quantitative proteomic methods. Data are from n=3 biologically independent replicates/group.

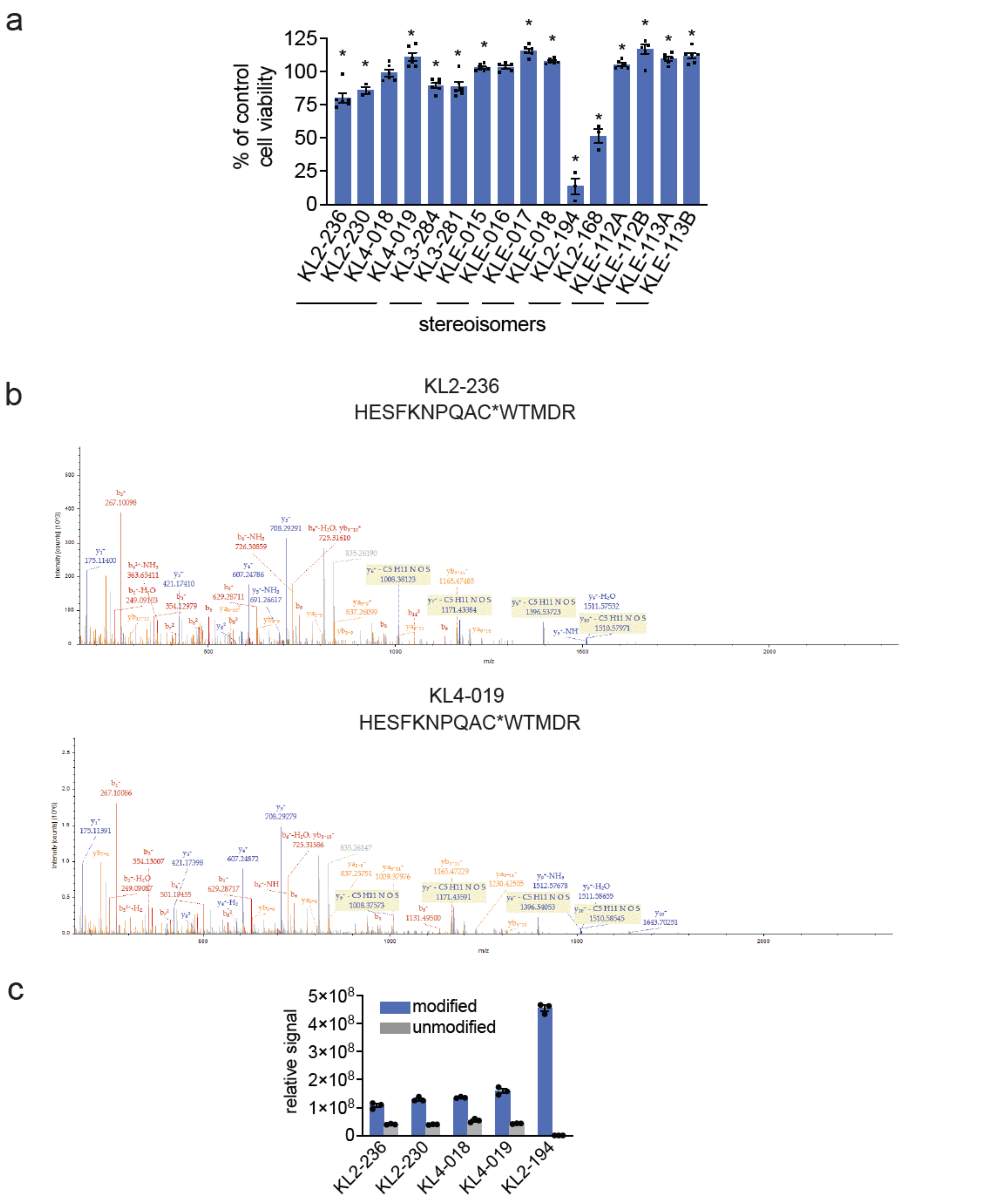


**Figure S1. Effects of Covalent Screening Compounds on Cell Viability and KL2-236 Adduct on MYC.** **(a)** Cell viability. LgBiT-MYC HEK293 cells were treated with DMSO vehicle or covalent compound (50 μM) for 24 h, and cell viability was assessed by CellTiter-Glo. **(b)** Reaction of KL2-236 and KL4-019 (50 μM) with artificial peptide HESFKNPQACWTMDR (1 μM, 1 hr). Shown are MS/MS spectra showing that only the cysteine was modified. **(c)** Relative reactivity of KL2-236, KL2-230, KL4-018, KL4-019, and KL2-194 (50 μM) with HESFKNPQACWTMDR (1 μM, 1 hr) where the mass spectrometry signal of the modified peptide was quantified by measuring area under the curve. Shown are the mass spectrometry signals for modified and unmodified peptides.

MYC C203
DLSAAASEC*IDPSVVFPYPL
m/z=485.63388

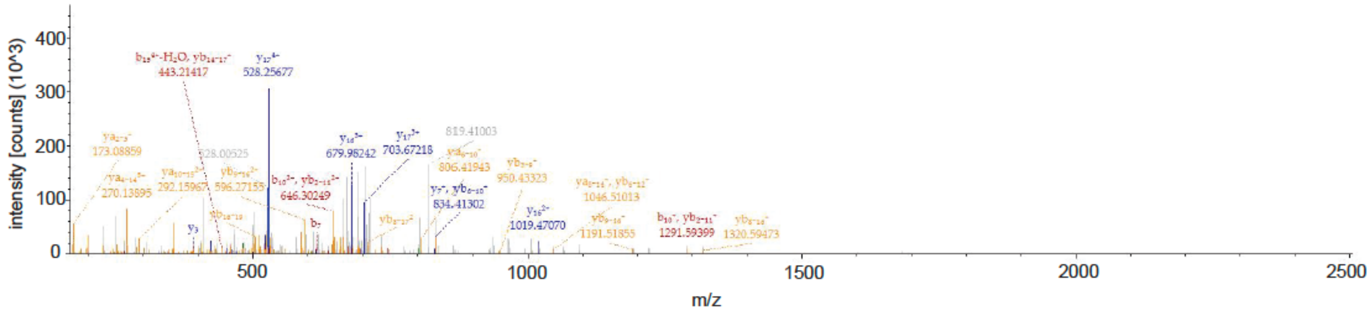


**Figure S2.** Reaction of KL2-236 with MYC based on incubation of KL2-236 (50 μM, 1 hr) with pure human MYC/MAX protein (1 μM), digestion of the complex with pepsin, and analysis of probe-modified peptides by LC-MS/MS**.** Shown is the MS/MS spectra of the KL2-236 modified peptide.

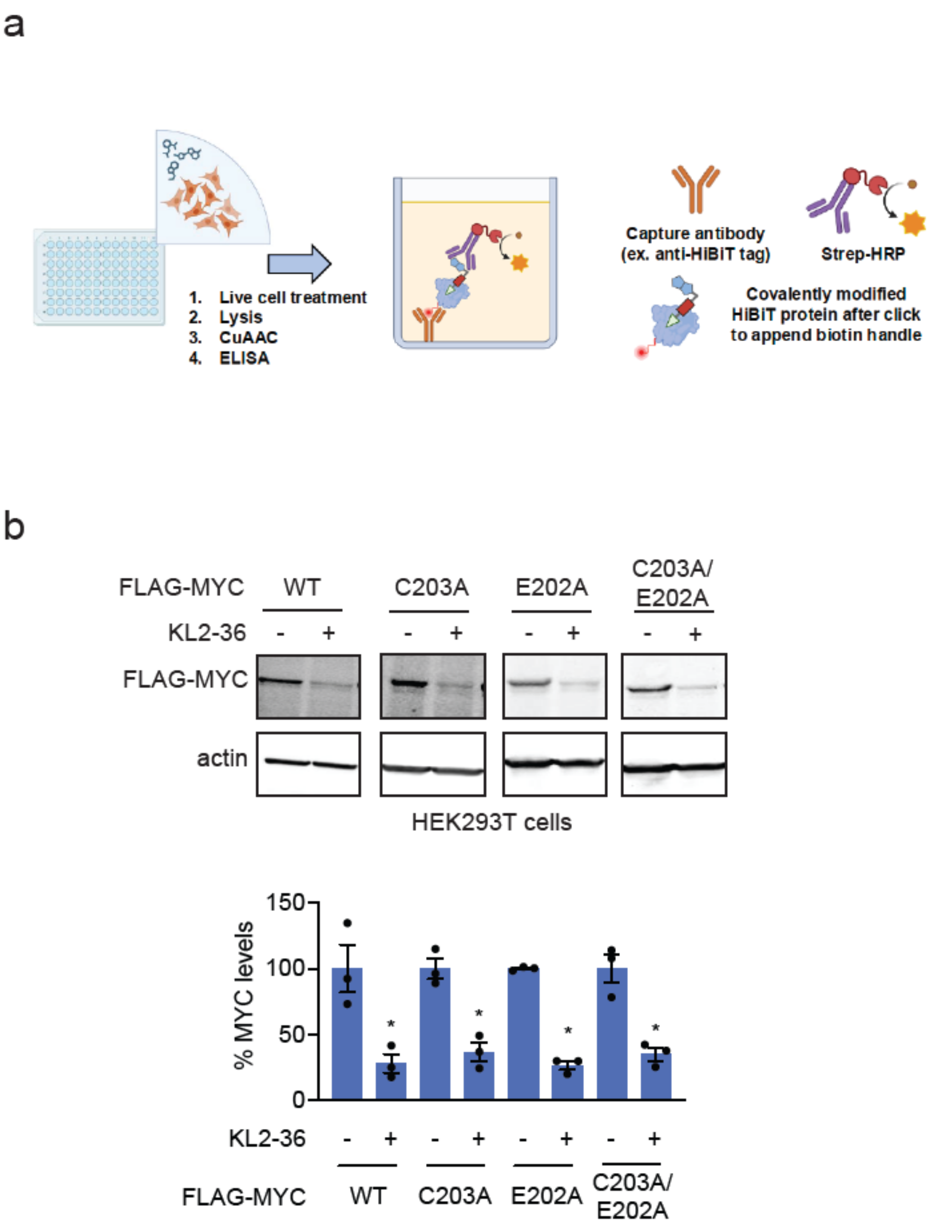


**Figure S3. ELISA-ABPP and FLAG-MYC degradation with KL2-236 treatment. (a)** Schematic of ELISA-ABPP based on data shown in **Figure 3e**. DLD-1 MYC HiBiT expressing cells were treated with DMSO vehicle, KL2-236, or KL4-019 for 1 h after which probe-modified proteins were appended with a biotin handle by CuAAC, and HiBiT-MYC was captured and immobilized on plate using a HiBiT antibody and then compound-engaged HiBiT-MYC was detected by streptavidin HRP. **(b)** HEK293T cells stably expressing FLAG-MYC wild-type (WT), C203A, E202A, C203A/E202A were treated with DMSO vehicle or KL2-236 (50 μM) for 24 h after which proteins were separated by SDS/PAGE and FLAG-MYC and loading control actin were detected and quantified by Western blotting. Blots represent n=3 biologically independent replicates per group. Data shows individual replicate values. Bar graph shows individual replicate and average ± sem values. Significance in expressed *p<0.05 compared to vehicle-treated controls.

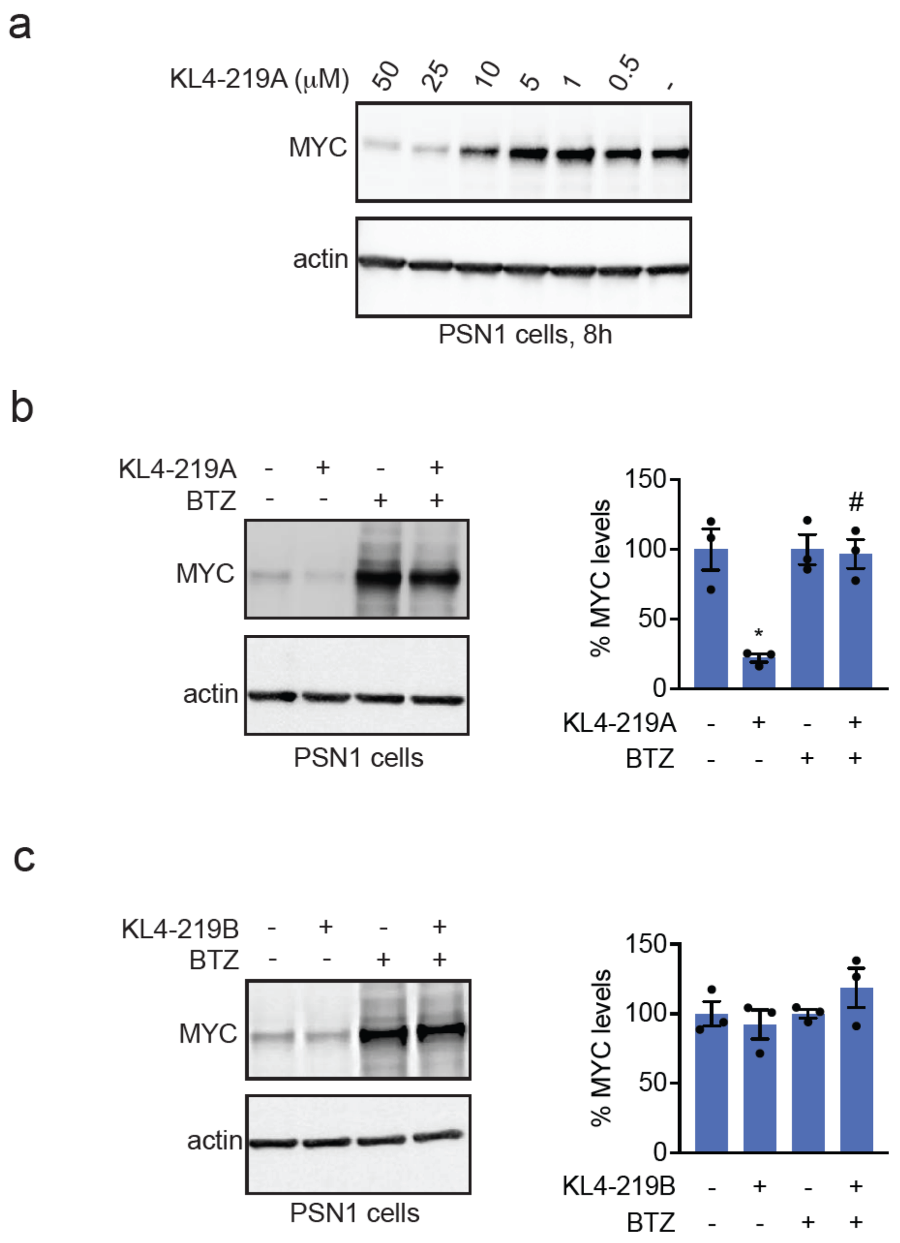


**Figure S4. Characterization of KL4-219A and KL4-219B in PSN1 cells. (a)** PSN1 cells were treated with a DMSO vehicle or KL4-219A for 8h, and MYC and loading control actin levels were assessed by Western blotting. **(b,c)** Proteasome-dependence of MYC degradation. PSN1 cells were pretreated with a DMSO vehicle or bortezomib (BTZ) (200 nM) for 1 hr prior to treatment of cells with a DMSO vehicle or KL4-219A or KL4-219B (25 μM) for 6h. MYC and loading control actin levels were assessed by Western blotting. Blots shown in **(a-c)** are representative of n=3 biologically independent replicates/group. Bar graphs in **(b,c)** show individual replicate values and average ± sem quantification of blots in **(b,c)**. Data in bar graphs have been normalized against average of DMSO control for DMSO and KL4-219A/B treatment groups and normalized against average of BTZ treatment for BTZ and BTZ and KL4-219A/B treatment groups. Statistical significance is shown as *p<0.05 compared to DMSO-treated controls and #p<0.05 compared to KL4-219A-treated groups.

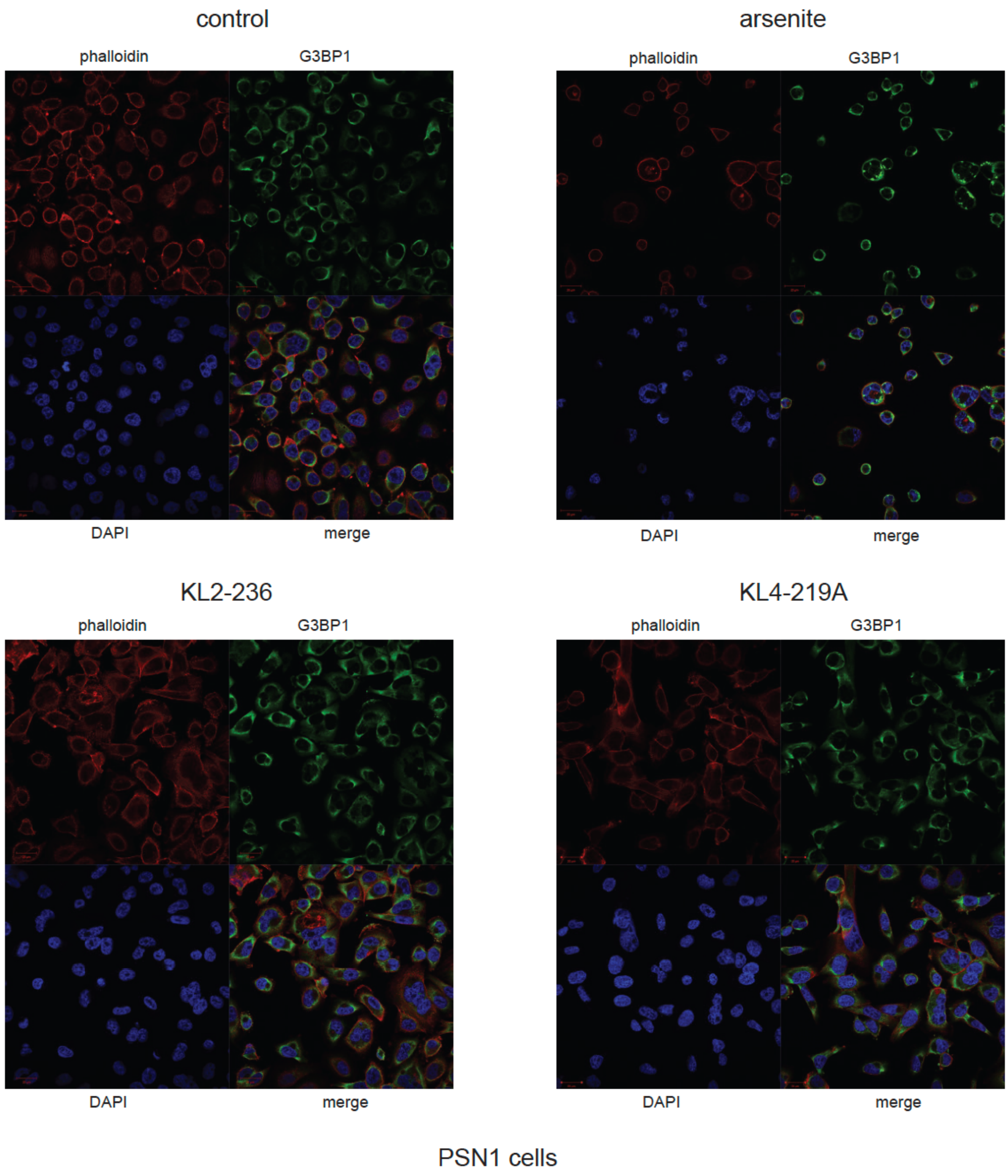


**Figure S5. Stress granule assessment.** PSN1 cells were treated with DMSO vehicle, KL2-236 (50 μM), KL4-219A (50 μM), or sodium arsenite (400 μM) for 4 h after which cells were fixed and visualized for actin marker phalloidin, G3BP1, or nuclear marker DAPI by microscopy. Shown are the individual channels and merged channels. Data are representative of n=3 biologically independent replicates per group.

### Synthetic Methods and Characterization

### General Procedures

All reactions were performed in flame- or oven-dried glassware under a positive pressure of nitrogen or argon, unless otherwise noted. Air- and moisture-sensitive liquids were transferred via syringe. When indicated, solvents or reagents were degassed by sparging with argon for 10 minutes in an ultrasound bath at 25 °C. Volatile solvents were removed under reduced pressure using rotary evaporation below 35 °C. Analytical and preparative thin-layer chromatography (TLC) were performed using glass plates pre-coated with silica gel (0.25-mm, 60-Å pore size, Merck TLC Silicagel 60 F254) impregnated with a fluorescent indicator (254 nm). TLC plates were visualized by exposure to ultraviolet light (UV) and then were stained by submersion in an ethanolic *p*-anisaldehyde solution, ceric ammonium molybdate solution, or potassium permanganate solution followed by brief heating on a hot plate. Flash column chromatography was performed with silica gel purchased from Fisher Scientific (Fisher Chemical™, 60 Å, 230-400 mesh, 40-63 µm). Synthetic procedures used to prepare oxindole sulfonyl and sulfinyl aziridines were adapted from the prior methods of Hajra and co-workers.[1]

Commercial solvents and reagents were used as received with the following exceptions. Anhydrous dichloromethane (DCM), dimethylformamide (DMF), and acetonitrile (MeCN) were obtained by passage through activated alumina columns. 1,4-dioxane (dioxane), triethylamine ($NEt_3$), and methanesulfonyl chloride (MsCl) were distilled over calcium hydride and stored under argon over activated 4Å molecular sieve beads.

Optical rotations were measured on a Perkin-Elmer 241 polarimeter at the D line (path length: 1 dm, cell volume: 1 mL, c in g/mL). The instrument was warmed up for 30 minutes before use. Optical rotations were conducted at room temperature, where room temperature is defined between 25 ºC to 30 ºC unless otherwise noted. Proton nuclear magnetic resonance ($^1$H NMR) spectra and carbon nuclear magnetic resonance ($^{13}$C NMR) spectra were recorded on Bruker NEO 500 (500 MHz/126MHz), Bruker AV 500 (500 MHz/126 MHz), Bruker AV 600 (600 MHz/151 MHz), and Bruker AV 700 (700 MHz/176 MHz) NMR spectrometers at 23 °C. Proton chemical shifts are expressed as parts per million (ppm, δ scale) and are referenced to residual solvent ($CHCl_3$: δ 7.26, $C_6D_5H$: δ 7.16), unless stated otherwise. Carbon chemical shifts are expressed as parts per million (ppm, δ scale) and are referenced to the solvent ($CDCl_3$: δ 77.16, $C_6D_6$: δ 128.06), unless stated otherwise. Data is represented as follows: chemical shift, multiplicity (s = singlet, d = doublet, dd = doublet of doublets, ddd, doublet of doublet of doublet, dt = triplet of doublets, t = triplet, hept = heptet, m = multiplet, br = broad), coupling constant (*J*) in Hertz (Hz), and integration. Infrared (IR) spectra were recorded on a Bruker Alpha FT-IR spectrometer as thin films and are reported in frequency of absorption ($cm^{-1}$). Only selected resonances are reported. High-resolution mass spectra were obtained by the QB3/chemistry mass spectrometry facility at the University of California, Berkeley using a Finnigan LTQFT mass spectrometer (Thermo Electron Corporation) with electrospray ionization (ESI).

**Compound Preparation and Characterization Data**

**General procedure A: Synthesis of *N*-sulfinyl spiroaziridines** (**KL2-236**, **KL2-230**, **KL4-018**, **KL4-019**, **KL3-284**, **KL3-281**, **KLE-015**, **KLE-016**, **KLE-017**, **KLE-108**, **KLE-112A**, **KLE-112B**, **KLE-113A**, **KLE-113B**). *i.* To a mixture of substituted isatin (0.250 mmol, 1 equiv.) and (*S*)-*tert*-butanesulfinamide (37.9 mg, 0.313 mmol, 1.25 equiv.) in anhydrous DCM (1 mL) at 20 °C was added $Ti(OR)_4$ (0.500 mmol, 2 equiv., R = Me, Et, or *i*-Pr) upon which the color of the mixture immediately darkened. The reaction vessel was sealed with Teflon tape and the reaction mixture was heated to 50 °C for 4 hours with stirring. The reaction was cooled to room temperature and diluted with EtOAc (3 mL). Excess $Ti(OR)_4$ was quenched by the addition of saturated *aq*. $NaHCO_3$ (2 mL), and the emulsion formed was vigorously stirred for 10 minutes at which point a white *aq*. suspension of $TiO_2$ separated. The mixture was filtered through a pad of celite and rinsed with EtOAc until colorless. The orange filtrate was washed with brine (10 mL), dried over $Na_2SO_4$, and concentrated in vacuo to yield the crude sulfinimine as a red oil that solidified upon standing. The crude sulfinimine was used immediately without further purification.

*ii.* A mixture of $Me_3SOI$ (110 mg, 0.500 mmol, 2 equiv.), $Cs_2CO_3$ (163 mg, 0.500 mmol, 2 equiv.), and powdered, activated 4Å molecular sieves (ca. 75 mg) in anhydrous MeCN (1 mL) was heated to 50 °C and stirred for 0.5 hours. After cooling to 20 °C, a solution of the crude sulfinimine (0.250 mmol, 1 equiv.) in anhydrous MeCN (1 mL) was added, and additional MeCN (2 x 0.5 mL) was used for quantitative transfer. The orange-colored reaction mixture was stirred at 20 °C for 1 hour during which the orange color gradually disappeared. The suspension, which was often beige-colored but sometimes green, blue, or purple, was filtered through a pad of celite, rinsing with EtOAc, to remove the 4Å molecular sieves. The filtrate was then washed with brine (5 mL), dried over $Na_2SO_4$, and concentrated in vacuo. The crude residue was purified by flash column chromatography on silica gel (20% →33% → 50% EtOAc–hexanes) then preparative TLC (15% EtOAc–toluene or 25% EtOAc–toluene) to yield (*S*,*S*)- and (*S*,*R*)-spiroaziridines respectively as white or yellow solids. The combined yield of the two diastereomers ranged from 40% to 60%, and the *d.r.* ranged from 3:1 to 4:1 (*S*,*S*):(*S*,*R*).

**General procedure B: Synthesis of *N*-sulfonyl aziridines** (**KL2-194** and **KL2-168**). To a solution of the *N*-sulfinyl aziridine (0.1 mmol, 1 equiv.) in DCM (1 mL) at 20 °C was added *m*-CPBA (77%, 67.2 mg, 0.300 mmol, 3 equiv.). The reaction mixture was stirred at 20 °C for 1 hour, then excess *m*-CPBA was quenched by addition of saturated *aq*. $Na_2S_2O_3$ (1 mL) and saturated *aq*. $NaHCO_3$ (1 mL). The biphasic mixture was vigorously stirred for 10 min., then the aqueous layer was extracted with EtOAc (3 x 2 mL). The combined organic layers were washed with brine (2 mL), dried over $Na_2SO_4$, then concentrated in vacuo. The crude residue was purified by two rounds of preparative TLC (33% EtOAc–hexanes then 15% EtOAc–toluene) to yield the *N*-sulfonyl aziridine as a white solid in 70% to 90% yield.

**General procedure C: Synthesis of morpholine amides** (**KLE-015**, **KLE-016**, **KLE-017**, **KLE-018**). *i.* To a solution of aryl ester (0.100 mmol, 1 equiv.) in a 1:9 mixture of $H_2O$:dioxane (1 mL) at 20 °C was added LiOH ·$H_2O$ (21.0 mg, 0.500 mmol, 5 equiv.). The reaction mixture was stirred at 60 °C for 1.5 h, allowed to cool to 20 °C, then diluted with EtOAc (2 mL). Excess LiOH was quenched by addition of *aq.* HCl (1.0 M, 1 mL), and the aqueous layer was extracted with EtOAc (5 x 1 mL). The combined organic layers were washed with brine (2 mL), dried over $Na_2SO_4$, then concentrated in vacuo to yield crude carboxylic acid as a yellow solid. The crude carboxylic acid was used immediately without further purification.

*ii.* To a mixture of the crude carboxylic acid and HOBt (14.9 mg, 0.110 mmol, 1.1 equiv.) in DCM (1 mL) at 20 °C was added morpholine (34.5 µL, 0.400 mmol, 4 equiv.) then DCC (22.7 mg, 0.110 mmol, 1.1 equiv.). The reaction mixture, which immediately turned cloudy, was stirred at 20 °C for 6 h, then filtered through a pad of celite and concentrated in vacuo. The crude residue was purified by two rounds of preparative TLC (EtOAc then 66% EtOAc–toluene + 5% MEOH) to yield the morpholine amides as colorless or slightly yellow oils or solids.

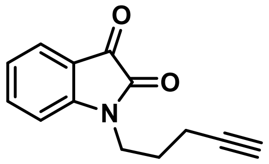

**SI-1. Requisite Isatin used for the preparation of KL2-236, KL2-230, KL4-018, KL4-019, KL2-194, and KL2-168** : $Cs_2CO_3$ (652 mg, 2.00 mmol, 2 equiv.) was added to a solution of 1H-Indole-2,3-dione (147 mg, 1.00 mmol, 1 equiv.) in anhydrous DMF (4 mL) at 20 °C. The deep purple-colored reaction mixture was stirred for five minutes, then pent-4-yn-1-yl methanesulfonate[2] (1.50 mmol, 1.5 equiv.) was added dropwise. The reaction mixture was stirred at 20 °C for 16 h, then diluted with EtOAc (10 mL). Excess $Cs_2CO_3$ was quenched with *aq.* HCl (1.0 M, 5 mL), and the aqueous layer was extracted with EtOAc (3 x 5 mL). The combined organic layers were washed with washed with 5% *aq*. LiCl (2 x 10 mL) and brine (10 mL) then dried over $Na_2SO_4$ and concentrated in vacuo. The crude residue was purified by flash column chromatography on silica gel (20%→33% EtOAc–hexanes) to yield the *N*-alkylated isatin **SI-1** (171.0 mg, 80.2%) as an orange solid. **$^1$H NMR** (700 MHz, $CDCl_3$) δ 7.63 – 7.57 (m, 2H), 7.12 (t, *J* = 7.5 Hz, 1H), 7.00 (d, *J* = 7.9 Hz, 1H), 3.86 (t, *J* = 7.3 Hz, 2H), 2.32 (td, *J* = 6.8, 2.6 Hz, 2H), 2.04 (t, *J* = 2.6 Hz, 1H), 1.94 (p, *J* = 6.9 Hz, 2H); **$^{13}$C NMR** (151 MHz, $CDCl_3$) δ 183.48, 158.44, 151.08, 138.53, 125.65, 123.88, 117.76, 110.23, 82.88, 69.86, 39.30, 26.12, 16.30; **IR** (thin film) $\nu_{max}$: 1736, 1610, 1469, 1353, 1170 cm$^{-1}$; **HRMS** (ESI-TOF) *m/z*: $[M+H]^+$ calcd for $C_{13}H_{11}NO_2$: 214.0863, found 214.0863.

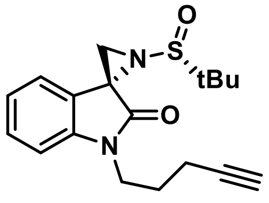

**KL2-230**: Prepared by general procedure A with **SI-1** (47.1 mg, 0.221 mmol), (*R*)-*tert*-butanesulfinamide, and Ti(O*i*-Pr)$_4$. **Yield**: 24.0 mg, 33%. **$^1$H NMR** (600 MHz, $CDCl_3$) δ

7.67 – 7.61 (m, 1H), 7.34 (td, *J* = 7.8, 1.2 Hz, 1H), 7.07 (td, *J* = 7.6, 1.0 Hz, 1H), 7.01 (dt, *J* = 7.9, 0.8 Hz, 1H), 3.93 – 3.84 (m, 2H), 3.37 (s, 1H), 2.78 (s, 1H), 2.30 (td, *J* = 6.9, 2.7 Hz, 2H), 2.02 (t, *J* = 2.7 Hz, 1H), 1.94 (p, *J* = 7.1 Hz, 2H), 1.31 (s, 9H); **$^{13}$C NMR** (126 MHz, $CDCl_3$) δ 171.90, 144.71, 129.64, 125.44, 122.81, 121.70, 108.99, 83.15, 69.53, 58.63, 44.30, 39.60, 31.33, 26.38, 22.59, 16.36; **IR** (thin film) $v_{max}$ 3249, 2953, 1727, 1611, 1468, 1372, 1173, 1078, 941, 765 cm$^{-1}$; **$[\alpha]_D^{25}$** = −217.6° (c 0.0013 g/ml, $CHCl_3$); **HRMS** (ESI-TOF) *m/z*: $[M+H]^+$ calcd for $C_{18}H_{23}N_2O_2S$: 331.1475, found 331.1475.

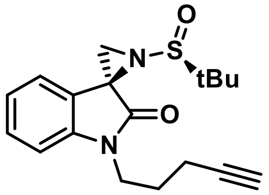

**KL2-236**: Prepared by general procedure A with **SI-1** (47.1 mg, 0.221 mmol), (*R*)-*tert*-butanesulfinamide, and Ti(O*i*-Pr)$_4$. **Yield**: 27.2 mg, 37%. **$^1$H NMR**, **$^{13}$C NMR**, and **IR** data matched that of **KL2-230**; **$[\alpha]_D^{25}$** = +240.5° (c 0.0037 g/ml, $CHCl_3$); **HRMS** (ESI-TOF) *m/z*: $[M+H]^+$ calcd for $C_{18}H_{23}N_2O_2S$: 331.1475, found 331.1475.

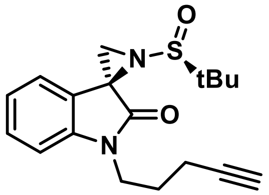

**KL4-018**: Prepared by general procedure A with **SI-1** (47.1 mg, 0.221 mmol), (*S*)-*tert*-butanesulfinamide, and Ti(O*i*-Pr)$_4$. **Yield**: 6.5 mg, 8.9%. **$^1$H NMR** (700 MHz, $CDCl_3$) δ 7.35 (ddd, *J* = 8.7, 6.0, 2.1 Hz, 1H), 7.12 – 7.05 (m, 2H), 6.99 (d, *J* = 7.9 Hz, 1H), 3.93 (dt, *J* = 14.4, 7.3 Hz, 1H), 3.86 (dt, *J* = 14.3, 7.2 Hz, 1H), 3.35 (s, 1H), 2.69 (s, 1H), 2.29 (td, *J* = 7.0, 2.7 Hz, 2H), 2.02 (t, *J* = 2.6 Hz, 1H), 1.94 (p, *J* = 7.1 Hz, 2H), 1.24 (s, 9H); **$^{13}$C NMR** (151 MHz, $CDCl_3$) δ 171.05, 143.60, 129.59, 125.31, 122.87, 121.98, 108.80, 83.18, 69.54, 57.89, 45.96, 39.75, 36.04, 26.57, 22.21, 16.37; **IR** (thin film) $v_{max}$: 3251, 2926, 1719, 1617, 1468, 1366, 1173, 938, 758 cm$^{-1}$; **$[\alpha]_D^{25}$** = −126.0° (c 0.001 g/ml, $CHCl_3$); **HRMS** (ESI-TOF) *m/z*: $[M+H]^+$ calcd for $C_{18}H_{23}N_2O_2S$: 331.1475, found 331.1476.

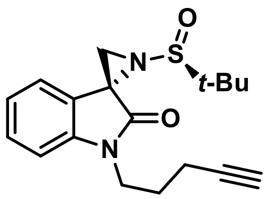

**KL4-019**: Prepared by general procedure A with **SI-1** (47.1 mg, 0.221 mmol), (*S*)-*tert*-butanesulfinamide, and Ti(O*i*-Pr)$_4$. **Yield**: 6.8 mg, 9.3%. **$^1$H NMR**, **$^{13}$C NMR**, and **IR** data matched that of **KL4-018**; **$[\alpha]_D^{25}$** = +117.0° (c 0.001 g/ml, $CHCl_3$); **HRMS** (ESI-TOF) *m/z*: $[M+H]^+$ calcd for $C_{18}H_{23}N_2O_2S$: 331.1475, found 331.1476.

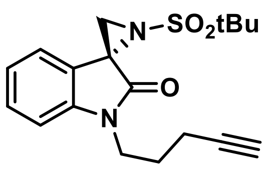

**KL2-168**: Prepared by general procedure B with **KL2-230** (38.2 mg, 0.116 mmol). **Yield**: 28.5 mg, 70.9%. **$^1$H NMR** (700 MHz, $CDCl_3$) δ 7.57 (br, 1H), 7.35 (td, *J* = 7.8,

1.2 Hz, 1H), 7.08 (td, $J$ = 7.6, 1.0 Hz, 1H), 6.99 (d, $J$ = 7.9 Hz, 1H), 3.89 (hept, $J$ = 7.2 Hz, 2H), 3.31 (s, 1H), 3.16 (s, 1H), 2.30 (tt, $J$ = 6.7, 2.6 Hz, 2H), 2.04 (t, $J$ = 2.7 Hz, 1H), 1.94 (p, $J$ = 7.1 Hz, 2H), 1.52 (s, 9H). **$^{13}$C NMR** (151 MHz, $C_6D_6$) δ 169.87, 144.91, 129.87, 125.82, 122.50, 121.52, 108.85, 83.17, 69.71, 61.17, 46.27, 41.24, 39.45, 26.32, 23.88, 16.20; **IR** (thin film) $\nu_{max}$: 3358, 2923, 2853, 1728, 1613, 1469, 1313, 1122, 958, 788 cm$^{-1}$; $[\alpha]_D^{25}$ = −17.1° (c 0.00082 g/ml, $CHCl_3$); **HRMS** (ESI-TOF) *m/z*: $[M+H]^+$ calcd for $C_{18}H_{23}N_2O_3S$: 347.1424, found 347.1426.

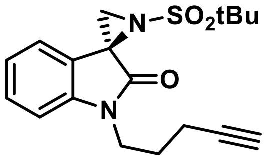


**KL2-194**: Prepared by general procedure B with **KL2-236** (31.2 mg, 0.0944 mmol). **Yield**: 28.1mg, 85.9%. **$^1$H NMR**, **$^{13}$C NMR**, and **IR** data matched that of **KL2-168**; $[\alpha]_D^{25}$ = +21.0° (c 0.001 g/ml, $CHCl_3$); **HRMS** (ESI-TOF) *m/z*: $[M+H]^+$ calcd for $C_{18}H_{23}N_2O_3S$: 347.1424, found 347.1426.

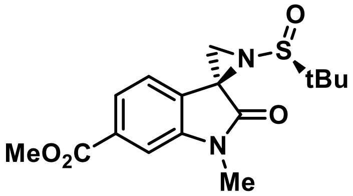


**SI-2**: Prepared by general procedure A with methyl 1-methyl-2,3-dioxoindoline-6-carboxylate[3] (50 mg, 0.228 mmol), (*S*)-*tert*-butanesulfinamide, and $Ti(OMe)_4$. **Yield**: 18.7 mg, 24%. **$^1$H NMR** (700 MHz, $CDCl_3$) δ 7.78 (dd, $J$ = 8.0, 1.5 Hz, 1H), 7.73 (d, $J$ = 7.9 Hz, 1H), 7.56 (d, $J$ = 1.5 Hz, 1H), 3.95 (s, 3H), 3.44 (s, 1H), 3.33 (s, 3H), 2.84 (s, 1H), 1.31 (s, 9H); **$^{13}$C NMR** (151 MHz, $CDCl_3$) δ 171.58, 166.63, 145.65, 131.51, 126.94, 125.21, 124.54, 109.28, 58.88, 52.57, 44.27, 31.75, 27.07, 22.58; **IR** (thin film) $\nu_{max}$: 2954, 2360, 1721, 1620, 1454, 1247, 1099, 938, 766 cm$^{-1}$; $[\alpha]_D^{25}$ = +308.0° (c 0.001 g/ml, $CHCl_3$); **HRMS** (ESI-TOF) *m/z*: $[M+H]^+$ calcd for $C_{16}H_{21}N_2O_4S$: 337.1217, found 337.1217.

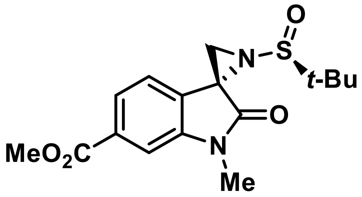


**SI-3**: Prepared by general procedure A with methyl 1-methyl-2,3-dioxoindoline-6-carboxylate (50 mg, 0.228 mmol), (*S*)-*tert*-butanesulfinamide, and $Ti(OMe)_4$. **Yield**: 4.4 mg, 5.7%. **$^1$H NMR** (700 MHz, $CDCl_3$) δ 7.81 (dd, $J$ = 7.7, 1.4 Hz, 1H), 7.55 (d, $J$ = 1.5 Hz, 1H), 7.15 (d, $J$ = 7.6 Hz, 1H), 3.95 (s, 3H), 3.40 (s, 1H), 3.34 (s, 3H), 2.75 (s, 1H), 1.24 (s, 9H); **$^{13}$C NMR** (151 MHz, $CDCl_3$) δ 170.71, 166.60, 144.44, 131.61, 130.27, 124.77, 121.66, 109.25, 58.08, 52.59, 45.86, 36.30, 27.09, 22.18; **IR** (thin film) $\nu_{max}$: 2954, 2360, 1718, 1624, 1453, 1246, 1084, 767 cm$^{-1}$; **[α]$_D^{25}$** = +78.7° (c 0.0015 g/ml, $CHCl_3$); **HRMS** (ESI-TOF) *m/z*: $[M+H]^+$ calcd for $C_{16}H_{21}N_2O_4S$: 337.1217, found 337.1219.

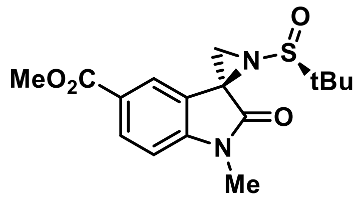

**SI-4**: Prepared by general procedure A with methyl 1-methyl-2,3-dioxoindoline-5-carboxylate[4] (94.0 mg, 0.429 mmol), (*S*)-*tert*-butanesulfinamide, and $Ti(OMe)_4$. **Yield**: 50.0 mg, 35%. **$^1$H NMR** (700 MHz, $CDCl_3$) δ 8.30 (s, 1H), 8.12 (dd, *J* = 8.2, 1.7 Hz, 1H), 6.97 (d, *J* = 8.2 Hz, 1H), 3.90 (s, 3H), 3.46 (s, 1H), 3.32 (s, 3H), 2.83 (s, 1H), 1.32 (s, 9H); **$^{13}$C NMR** (151 MHz, $CDCl_3$) δ 172.28, 166.80, 149.24, 132.36, 126.57, 125.08, 121.74, 108.48, 58.86, 52.40, 44.02, 31.52, 27.08, 22.65; **IR** (thin film) $v_{max}$: 2922, 2360, 1716, 1617, 1365, 1266, 1106, 934, 768 cm$^{-1}$; **[α]$_D^{25}$** = +42.0° (c 0.001 g/ml, $CHCl_3$); **HRMS** (ESI-TOF) *m/z*: $[M+H]^+$ calcd for $C_{16}H_{21}N_2O_4S$: 337.1217, found 337.1218.

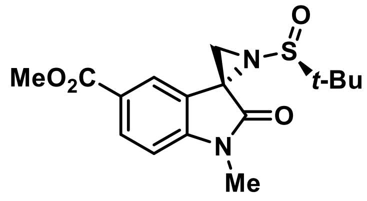

**SI-5**: Prepared by general procedure A with methyl 1-methyl-2,3-dioxoindoline-5-carboxylate (94.0 mg, 0.429 mmol), (*S*)-*tert*-butanesulfinamide, and $Ti(OMe)_4$. **Yield**: 15.5 mg, 10.7%. **$^1$H NMR** (700 MHz, $CDCl_3$) δ 8.10 (dd, J = 8.2, 1.7 Hz, 1H), 7.73 (s, 1H), 6.94 (d, J = 8.2 Hz, 1H), 3.92 (s, 3H), 3.40 (s, 1H), 3.32 (s, 3H), 2.74 (s, 1H), 1.26 (s, 9H); **$^{13}$C NMR** (151 MHz, $CDCl_3$) δ 171.31, 166.69, 148.11, 132.11, 125.39, 125.08, 122.99, 108.20, 58.14, 52.33, 45.50, 35.79, 27.09, 22.19; **IR** (thin film) $v_{max}$: 2953, 2360, 1715, 1623, 1367, 1265, 1099, 767 cm$^{-1}$; **[α]$_D^{25}$** = +190.6° (c 0.0027 g/ml, $CHCl_3$); **HRMS** (ESI-TOF) *m/z*: $[M+H]^+$ calcd for $C_{16}H_{21}N_2O_4S$: 337.1217, found 337.1219.

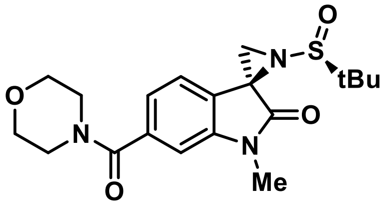

**KLE-015**: Prepared by general procedure C with **SI-2** (6.1 mg, 0.019 mmol). **Yield**: 2.8 mg, 38%. **$^1$H NMR** (500 MHz, $CDCl_3$) δ 7.68 (d, *J* = 7.7 Hz, 1H), 7.05 (dd, *J* = 7.7, 1.4 Hz, 1H), 7.02 (d, *J* = 1.4 Hz, 1H), 3.91 – 3.44 (br, 8H), 3.42 (s, 1H), 3.29 (s, 3H), 2.83 (s, 1H), 1.31 (s, 9H); **$^{13}$C NMR** (151 MHz, $CDCl_3$) δ 171.71, 169.75, 146.05, 136.80, 125.21, 123.53, 121.22, 107.97, 67.02, 58.83, 48.46, 44.13, 42.82, 31.40, 27.00, 22.59; **IR** (thin film) $v_{max}$: 2922, 2361, 1732, 1622, 1463, 1249, 1113, 765 cm$^{-1}$; **[α]$_D^{25}$** = +226.0° (c 0.001 g/ml, $CHCl_3$); **HRMS**

(ESI-TOF) *m/z*: $[M+H]^+$ calcd for $C_{19}H_{26}N_3O_4S$: 392.1639, found 392.1641.

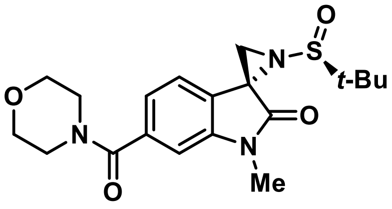

**KLE-016**: Prepared by general procedure C with **SI-3** (7.7 mg, 0.024 mmol). **Yield**: 6.7 mg, 71%. **$^1$H NMR** (500 MHz, $CDCl_3$) δ 7.12 (d, *J* = 7.9 Hz, 1H), 7.07 (dd, *J* = 7.5, 1.3 Hz, 1H), 7.00 – 6.96 (m, 1H), 3.91 – 3.42 (br, 8H), 3.37 (s, 1H), 3.30 (s, 3H), 2.73 (s, 1H), 1.24 (s, 9H); **$^{13}$C NMR** (151 MHz, $CDCl_3$) δ 170.83, 169.77, 144.80, 136.93, 126.91, 121.80, 121.37, 107.68, 67.02, 58.02, 45.79, 36.22, 27.03, 22.18; **IR** (thin film) $v_{max}$: 2923, 1724, 1624, 1463, 1247, 1113 $cm^{-1}$; **$[\alpha]_D^{25}$** = +120.0° (c 0.001 g/ml, $CHCl_3$); **HRMS** (ESI-TOF) *m/z*: $[M+H]^+$ calcd for $C_{19}H_{26}N_3O_4S$: 392.1639, found 392.1641.

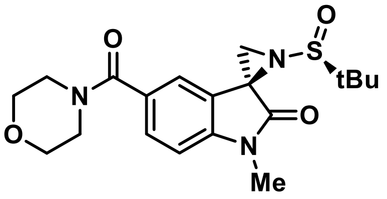

**KLE-017**: Prepared by general procedure C with **SI-4** (7.4 mg, 0.023 mmol). **Yield**, 3.1 mg, 34%. **$^1$H NMR** (500 MHz, $CDCl_3$) δ 7.72 (d, J = 1.6 Hz, 1H), 7.57 (dd, J = 8.1, 1.7 Hz, 1H), 6.98 (d, J = 8.1 Hz, 1H), 3.68 (br, 8H), 3.41 (s, 1H), 3.30 (s, 3H), 2.82 (s, 1H), 1.30 (s, 9H); **$^{13}$C NMR** (151 MHz, $CDCl_3$) δ 171.94, 169.91, 146.90, 130.35, 129.70, 124.51, 121.29, 109.04, 66.99, 58.79, 44.16, 31.23, 27.03, 22.59; **IR** (thin film) $v_{max}$: 2920, 1731, 1616, 1422, 1250, 1112, 923, 764 $cm^{-1}$; **$[\alpha]_D^{25}$** = +486.7° (c 0.00015 g/ml, $CHCl_3$); **HRMS** (ESI-TOF) *m/z*: $[M+H]^+$ calcd for $C_{19}H_{26}N_3O_4S$: 392.1639, found 392.1641.

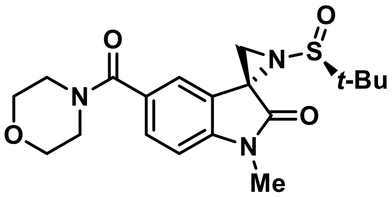

**KLE-018**: Prepared by general procedure C with **SI-5** (8.6 mg, 0.027 mmol). **Yield**: 2.8 mg, 26%. **$^1$H NMR** (700 MHz, $CDCl_3$) δ 7.44 (dd, *J* = 8.0, 1.7 Hz, 1H), 7.21 (br, 1H), 6.92 (d, *J* = 8.0 Hz, 1H), 3.70 (br, 8H), 3.36 (d, *J* = 11.9 Hz, 1H), 3.31 (s, 3H), 2.74 (s, 1H), 1.25 (s, 9H); **$^{13}$C NMR** (151 MHz, $CDCl_3$) δ 171.04, 170.02, 145.73, 129.76, 129.23, 125.48, 121.63, 108.27, 67.00, 58.14, 45.66, 36.06, 27.06, 22.23; **$[\alpha]_D^{25}$** = +54.0° (c 0.0005 g/ml, $CHCl_3$); **HRMS** (ESI-TOF) *m/z*: $[M+H]^+$ calcd for $C_{19}H_{26}N_3O_4S$: 392.1639, found 392.1639.

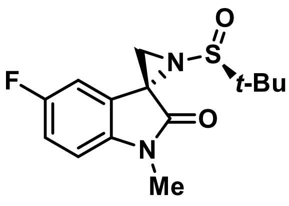

**KLE-112B**: Prepared by general procedure A with 5-fluoro-1-methylindoline-2,3-dione[5] (30 mg, 0.167 mmol), (*S*)-*tert*-butanesulfinamide, and Ti(O*i*-Pr)$_4$. **Yield**: 4.7 mg, 9.5%. **$^1$H NMR** (700 MHz, $CDCl_3$) δ 7.09 – 7.02 (m, 1H), 6.82 (m, 2H), 3.36 (s,

1H), 3.28 (s, 3H), 2.68 (s, 1H), 1.24 (s, 9H); **$^{13}$C NMR** (151 MHz, $CDCl_3$) δ 170.73, 159.51 (d, J = 241.5 Hz), 140.09 (d, J = 2.1 Hz), 126.89 (d, J = 8.2 Hz), 115.81 (d, J = 23.8 Hz), 109.97 (d, J = 25.7 Hz), 109.21 (d, J = 8.0 Hz), 57.97, 46.03 (d, *J* = 2.2 Hz), 35.97, 26.99, 22.14; **IR** (thin film) $\nu_{max}$: 2925, 1719, 1493, 1367, 1277, 1089, 945 cm$^{-1}$; **$[\alpha]^{25}_{D}$** = +231.8° (c 0.00073 g/ml, $CHCl_3$); **HRMS** (ESI-TOF) *m/z*: $[M+H]^+$ calcd for $C_{14}H_{18}FN_2O_2S$: 297.1068, found 297.1069.

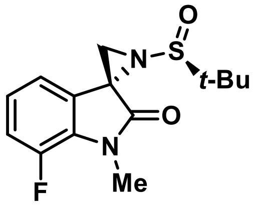

**KLE-113B**: Prepared by general procedure A with 7-fluoro-1-methylindoline-2,3-dione[6] (89.6 mg, 0.500 mmol), (*S*)-*tert*-butanesulfinamide, and Ti(O*i*-Pr)$_4$. **Yield**: 21.0 mg, 14.2%. **$^1$H NMR** (700 MHz, $CDCl_3$) δ 7.07 (ddd, *J* = 11.4, 8.4, 1.0 Hz, 1H), 7.00 (td, *J* = 7.9, 4.3 Hz, 1H), 6.85 (d, *J* = 7.4 Hz, 1H), 3.49 (d, *J* = 2.6 Hz, 3H), 3.36 (s, 1H), 2.67 (s, 1H), 1.24 (s, 9H); **$^{13}$C NMR** (151 MHz, $CDCl_3$) δ 170.70, 147.89 (d, J = 244.4 Hz), 130.73 (d, J = 9.1 Hz), 128.15, 123.54 (d, J = 6.5 Hz), 122.39 (d, J = 6.7 Hz), 117.56 (d, J = 19.3 Hz), 57.96, 45.92 (d, J = 3.3 Hz), 36.34, 29.43 (d, J = 5.5 Hz), 22.14; **IR** (thin film) $\nu_{max}$: 2925, 2361, 1728, 1479, 1371, 1238, 1098, 972, 731 cm$^{-1}$; **$[\alpha]^{25}_{D}$** = +142.5° (c 0.00027 g/ml, $CHCl_3$); **HRMS** (ESI-TOF) *m/z*: $[M+H]^+$ calcd for $C_{14}H_{18}FN_2O_2S$: 297.1068, found 297.1069.

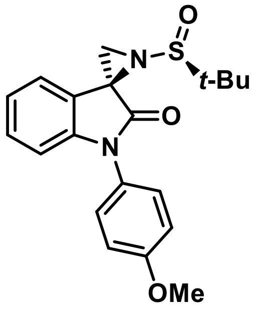

**KL4-219A**: Prepared by general procedure A with 1-(4-methoxyphenyl)indoline-2,3-dione[7] (16.6 mg, 0.0656 mmol), (*S*)-*tert*-butanesulfinamide, and Ti(O*i*-Pr)$_4$. **Yield**: 9.2 mg, 38%. **$^1$H NMR** (600 MHz, $CDCl_3$) δ 7.71 (d, *J* = 7.6 Hz, 1H), 7.37 – 7.32 (m, 2H), 7.26 (t, 1H), 7.10 (td, *J* = 7.6, 1.0 Hz, 1H), 7.07 – 7.01 (m, 2H), 6.84 (d, *J* = 7.9 Hz, 1H), 3.86 (s, 3H), 3.45 (s, 1H), 2.87 (s, 1H), 1.33 (s, 9H); **$^{13}$C NMR** (151 MHz, $CDCl_3$) δ 171.42, 159.47, 145.89, 129.53, 128.10, 126.86, 125.45, 123.21, 121.39, 115.08, 110.06, 58.75, 55.69, 44.50, 31.69, 22.63; **IR** (thin film) $\nu_{max}$: 2959, 2360, 1735, 1611, 1514, 1250, 1080, 938, 759 cm$^{-1}$; **$[\alpha]^{25}_{D}$** = +189.0° (c 0.001 g/ml, $CHCl_3$); **HRMS** (ESI-TOF) *m/z*: $[M+H]^+$ calcd for $C_{20}H_{23}N_2O_3S$: 371.1424, found 371.1424.

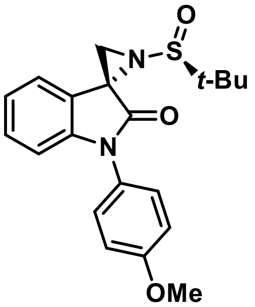


**KL4-219B**: Prepared by general procedure A with 1-(4-methoxyphenyl)indoline-2,3-dione (16.6 mg, 0.0656 mmol), (*S*)-*tert*-butanesulfinamide, and $Ti(O\textit{i}-Pr)_4$. **Yield**: 2.7 mg, 11%. **$^1$H NMR** (700 MHz, $CDCl_3$) δ 7.37 – 7.33 (m, 2H), 7.28 (td, *J* = 7.8, 1.4 Hz, 1H), 7.15 (s, 1H), 7.11 (t, *J* = 7.4 Hz, 1H), 7.05 – 7.00 (m, 2H), 6.87 (d, *J* = 7.9 Hz, 1H), 3.86 (s, 3H), 3.42 (s, 1H), 2.78 (s, 1H), 1.27 (s, 9H); **$^{13}$C NMR** (126 MHz, $CDCl_3$) δ 170.54, 159.37, 144.53, 129.48, 127.82, 126.85, 124.98, 123.30, 122.04, 114.95, 109.88, 57.94, 55.70, 46.20, 36.71, 22.22.; **IR** (thin film) $\nu_{max}$: 2924, 2361, 1727, 1615, 1514, 1251, 1091, 923, 751 cm$^{-1}$; $[\alpha]_D^{25}$ = +63.8° (c 0.00091 g/ml, $CHCl_3$); **HRMS** (ESI-TOF) *m/z*: $[M+H]^+$ calcd for $C_{20}H_{23}N_2O_3S$: 371.1424, found 371.1427.

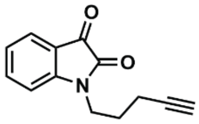

**SI-1**
(700 MHz, $CDCl_3$)

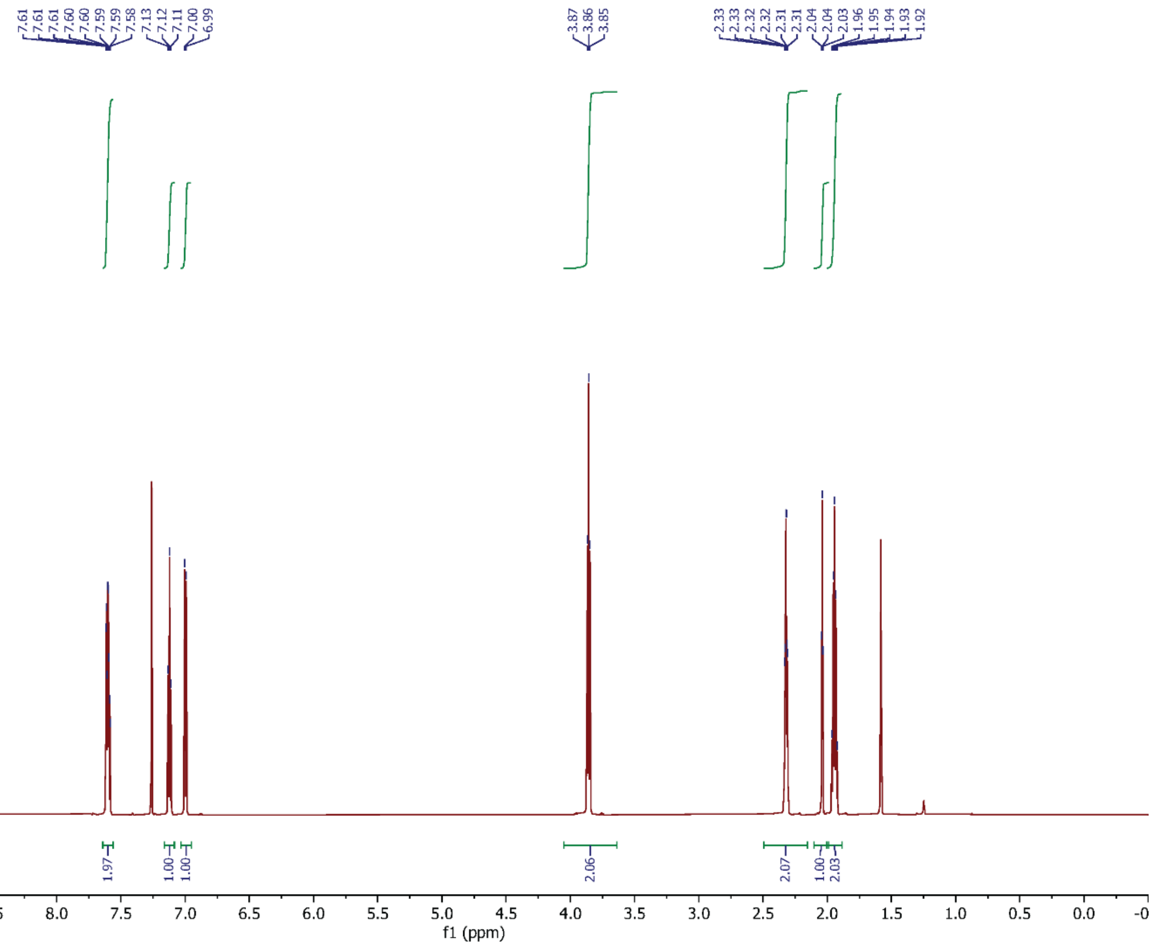


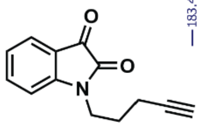

**SI-1**
(151 MHz, $CDCl_3$)

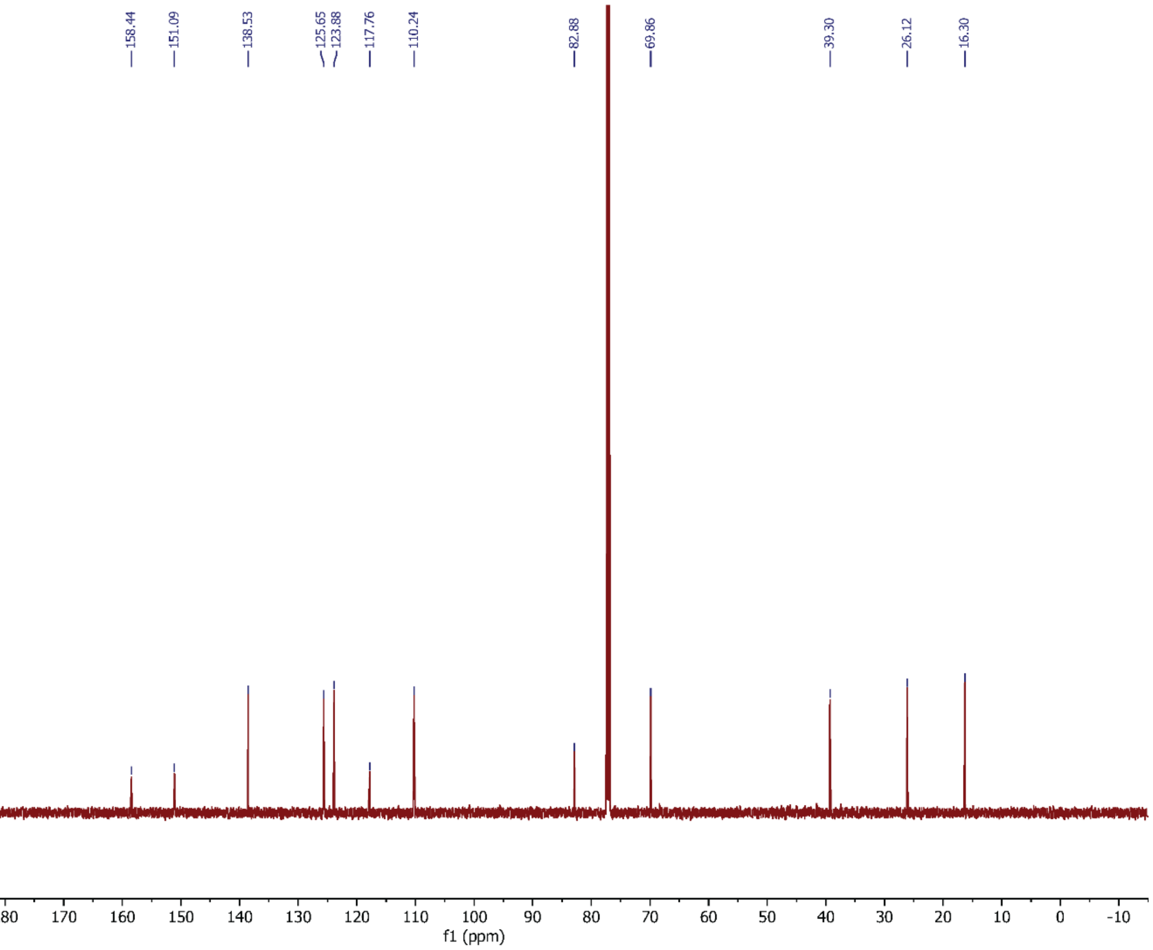

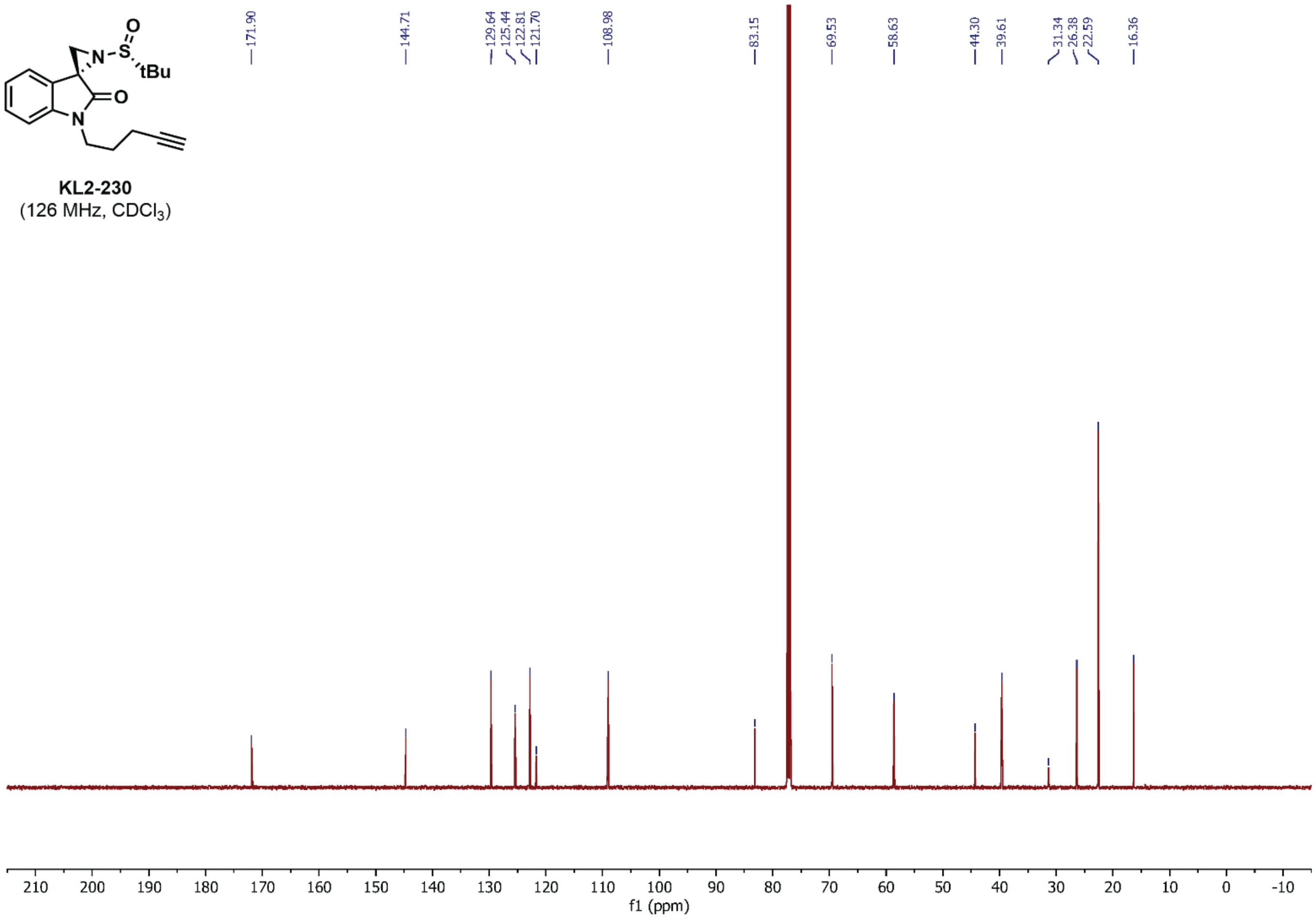


**KL2-230**
(126 MHz, $CDCl_3$)

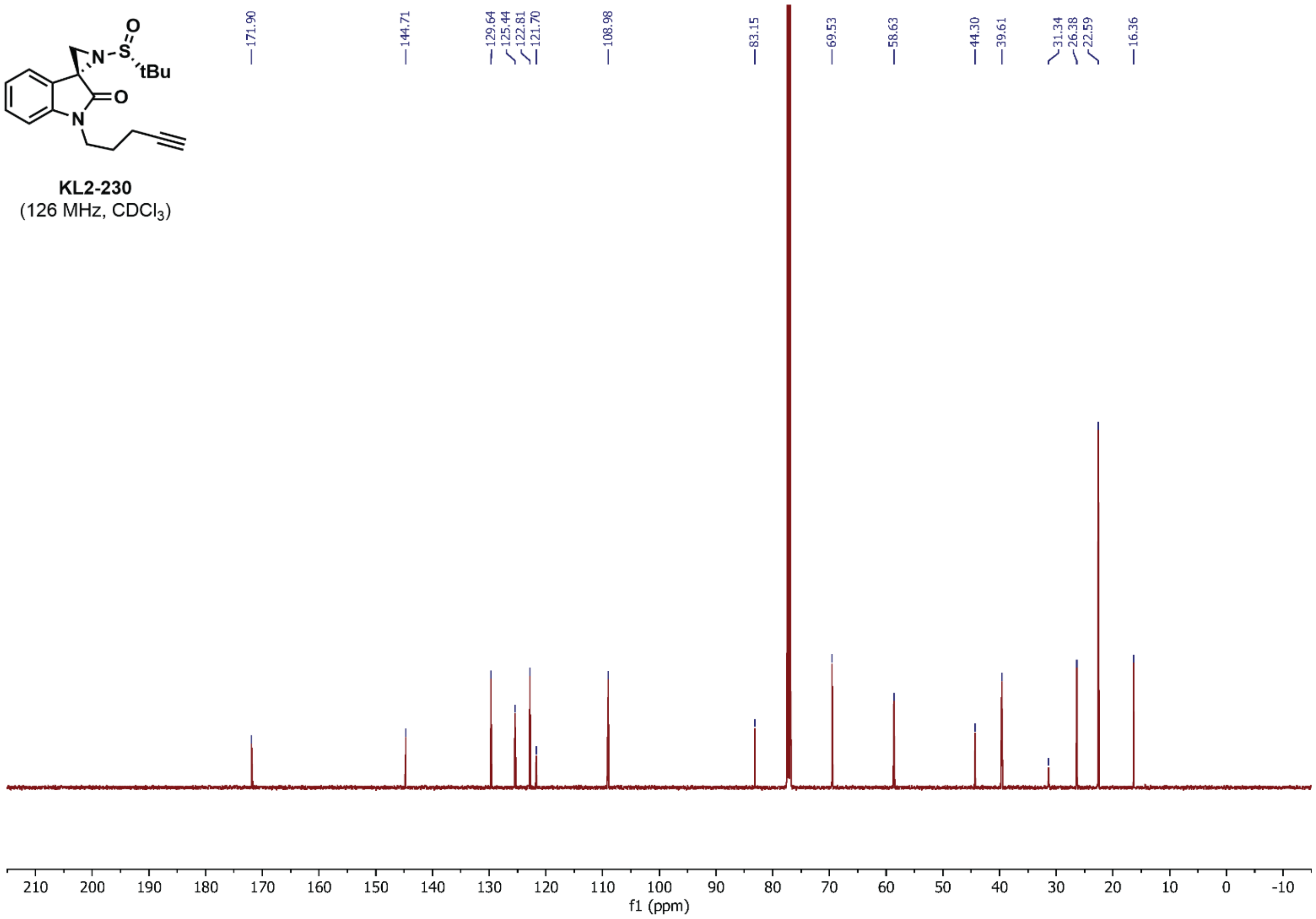


**KL2-230**
(126 MHz, $CDCl_3$)

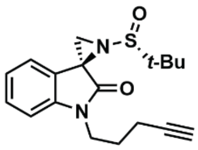


**KL4-018**
(700 MHz, $CDCl_3$)

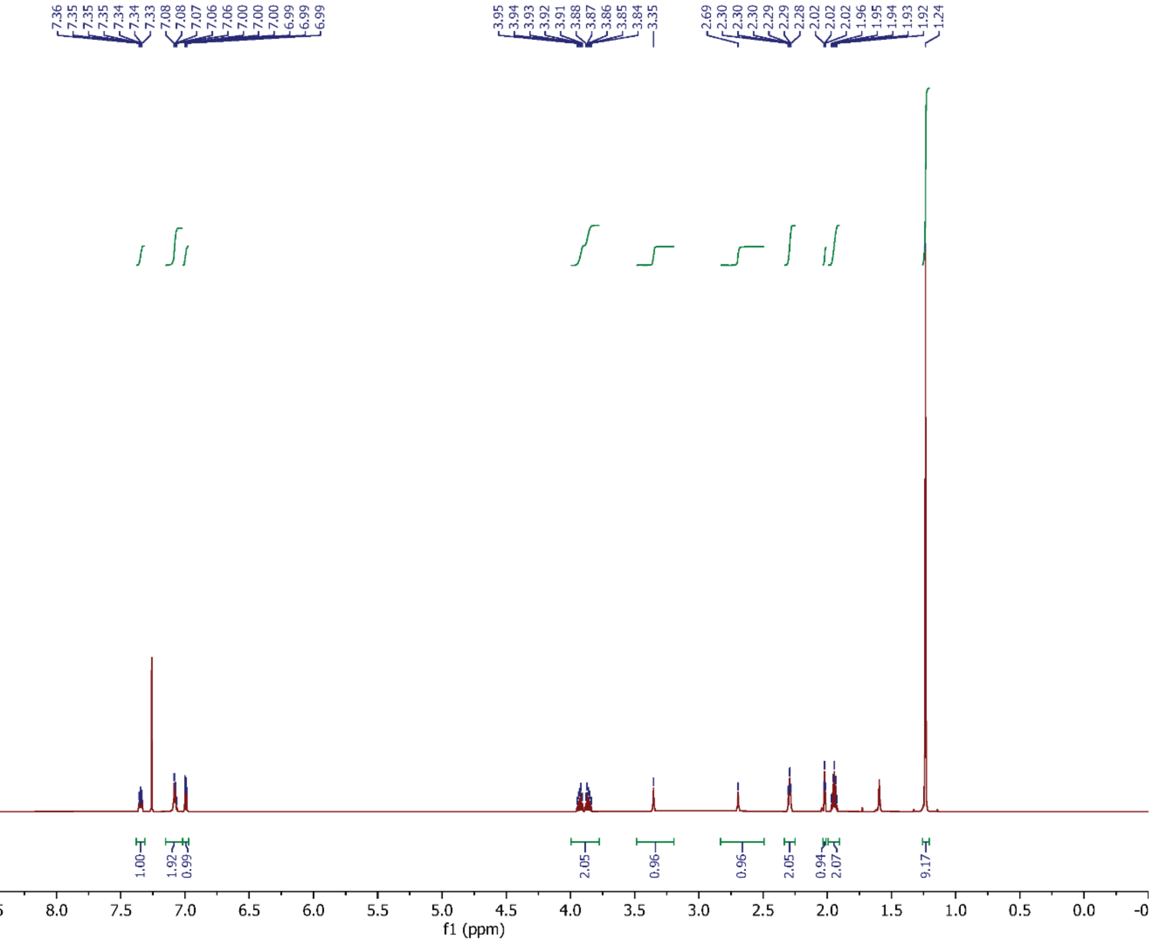


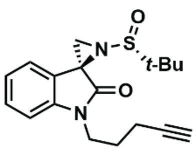


**KL4-018**
(151 MHz, $CDCl_3$)

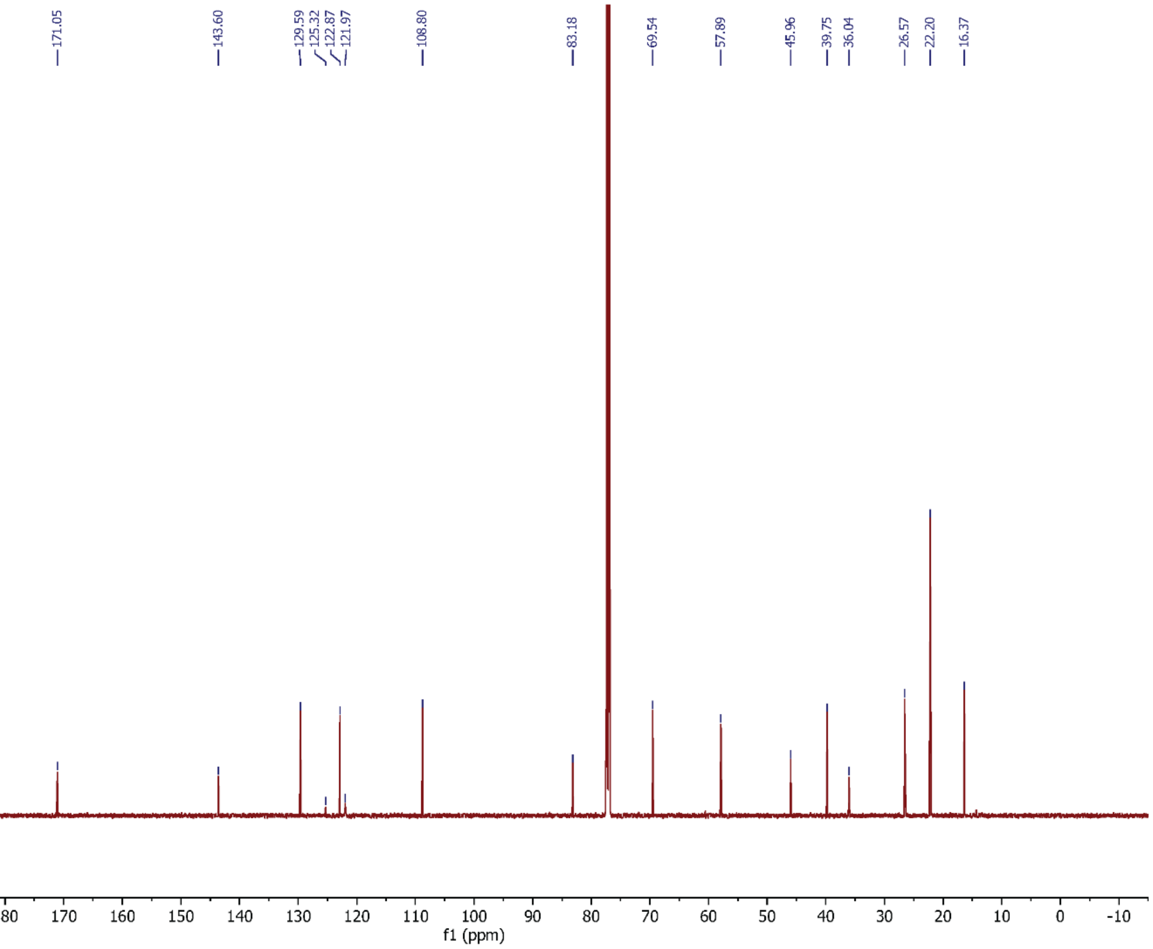

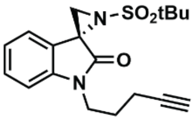


**KL2-168**
(700 MHz, $CDCl_3$)

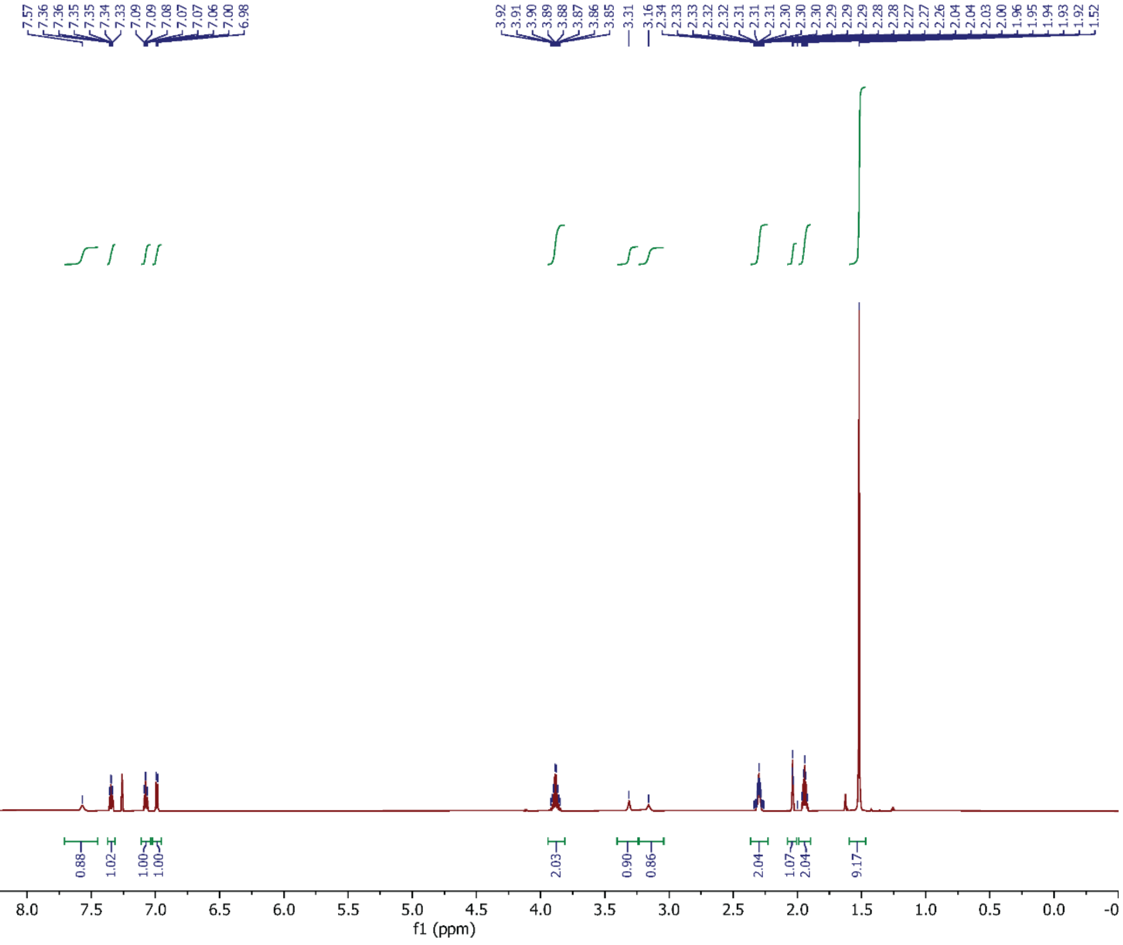


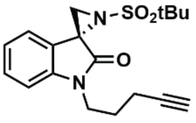


**KL2-168**
(151 MHz, $C_6D_6$)

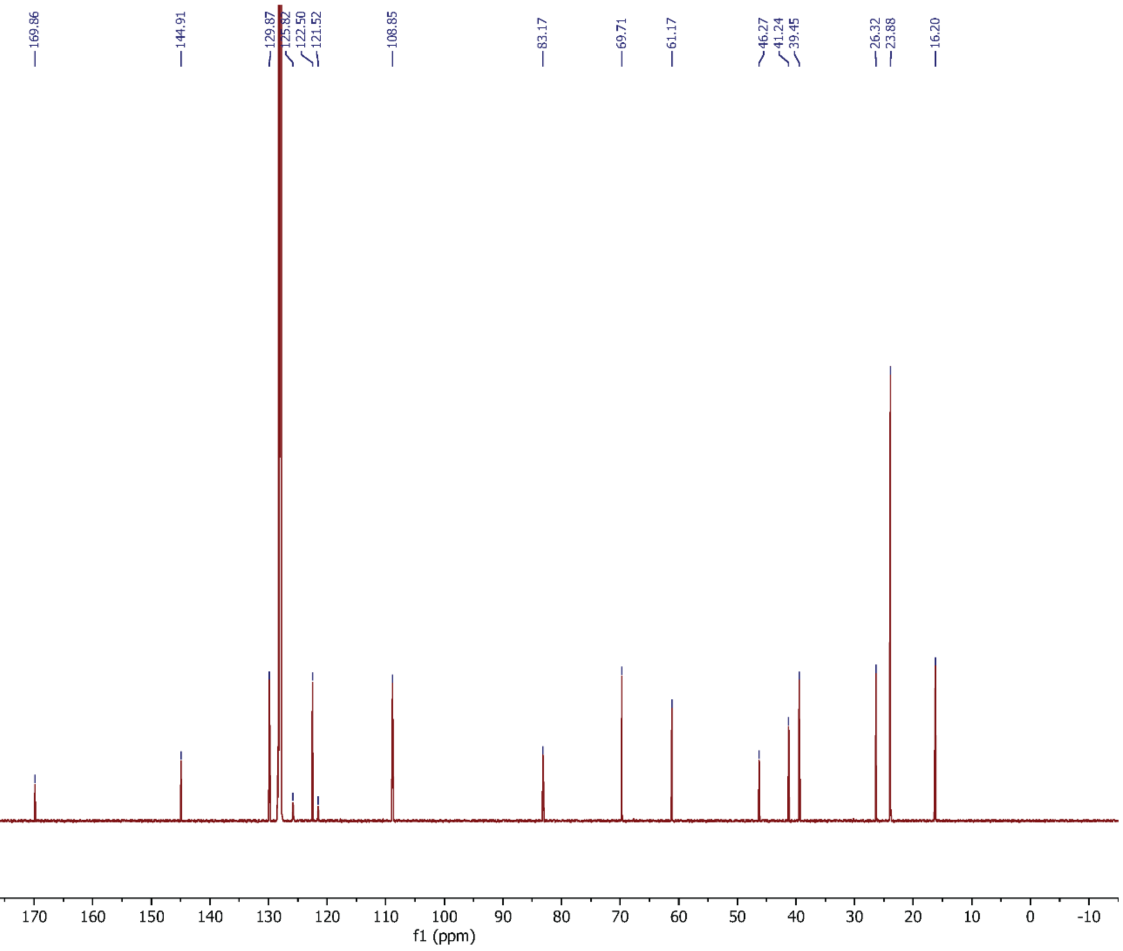

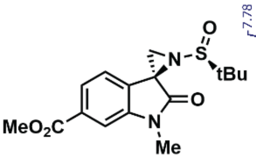

SI-2
(700 MHz, $CDCl_3$)

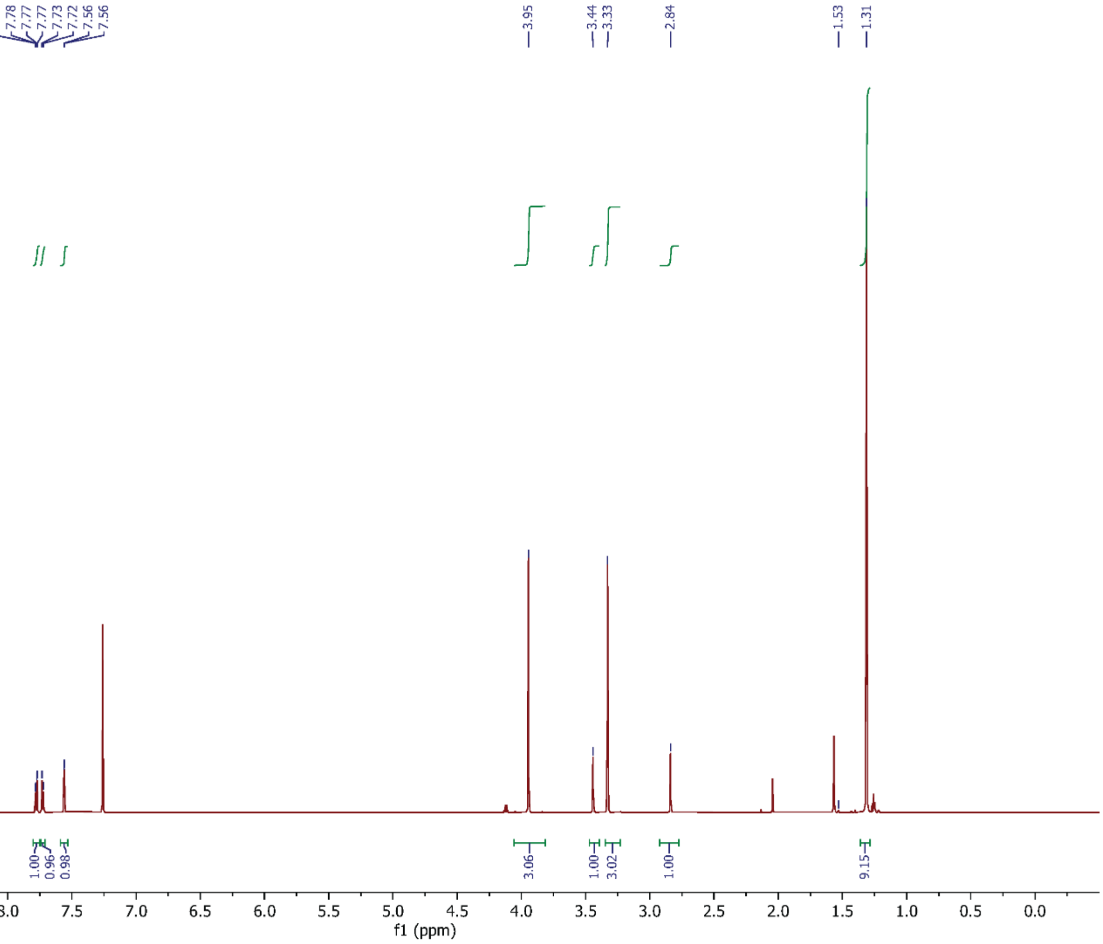


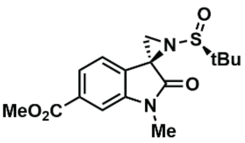

SI-2
(151 MHz, $CDCl_3$)

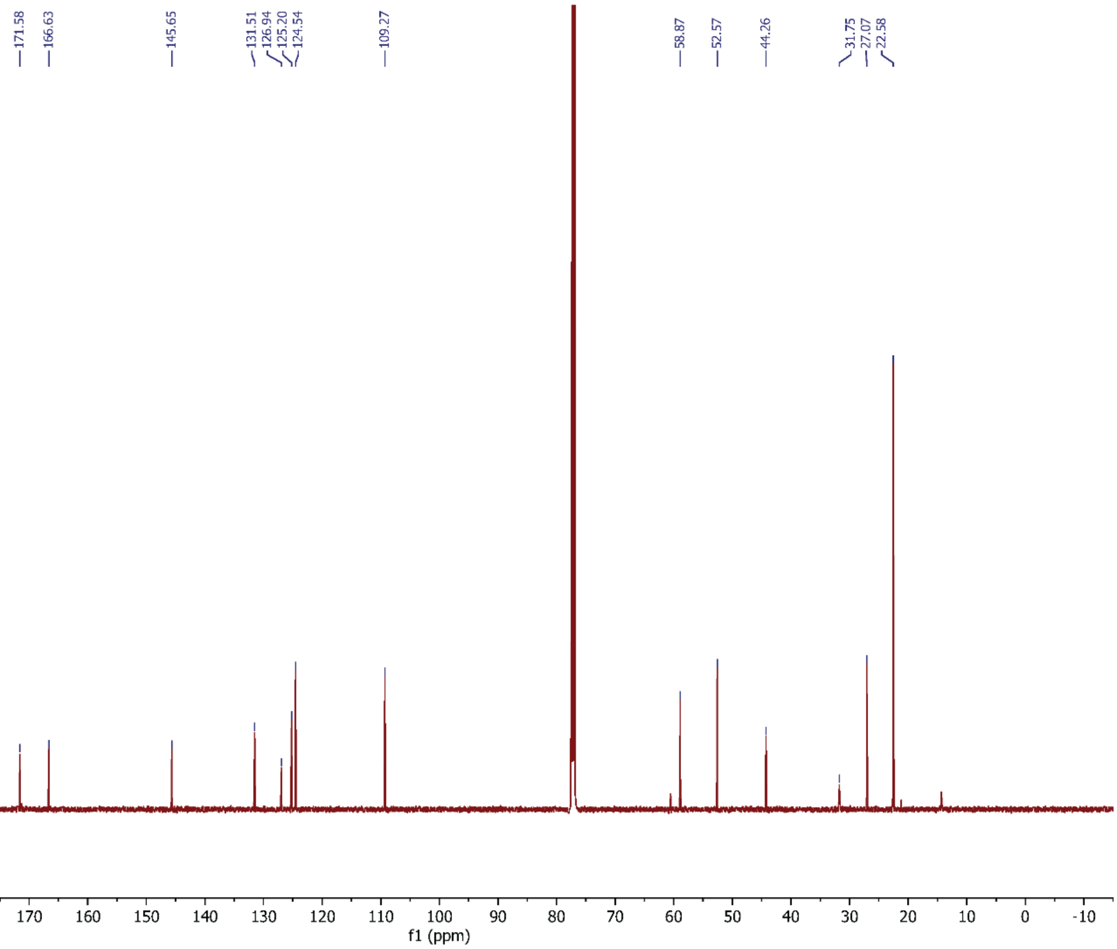

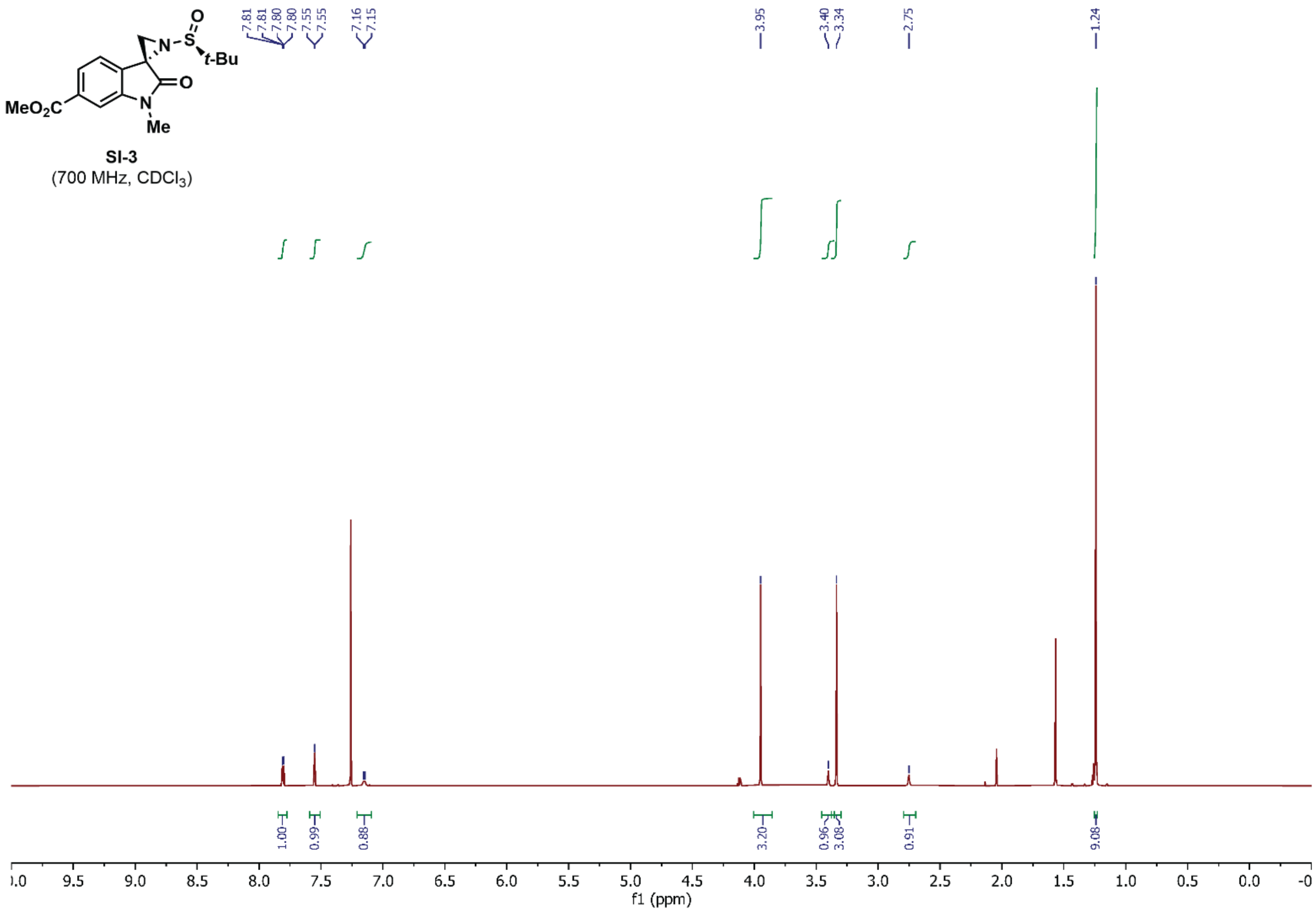

MeO2C
N
Me
O
N
S
O
t-Bu
SI-3
(700 MHz, CDCl3)
7.81 7.81 7.80 7.80 7.55 7.55 7.16 7.15
3.95
3.40 3.34
2.75
1.24
1.00 0.99 0.88 3.20 0.96 3.08 0.91 9.08
f1 (ppm)


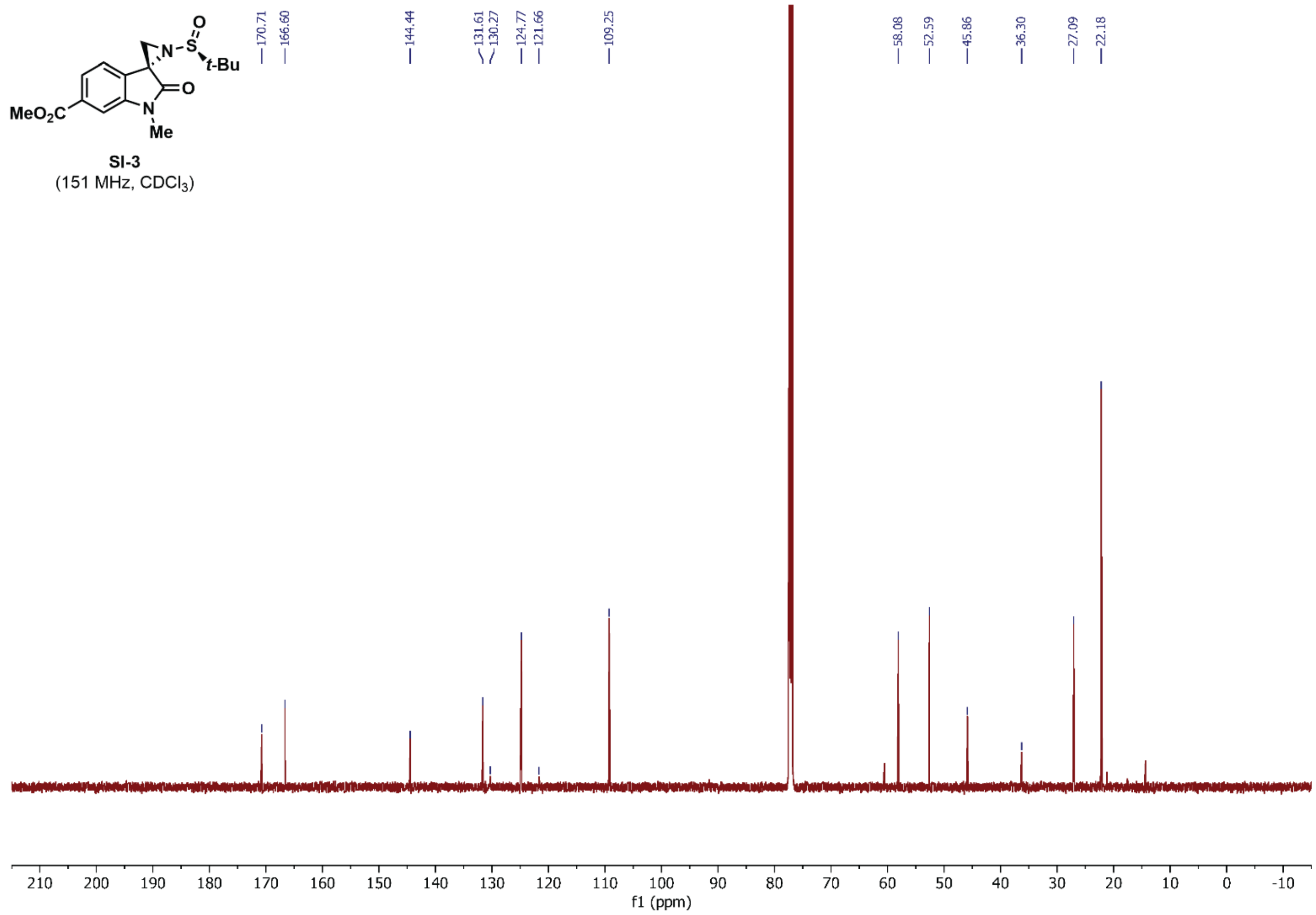

MeO2C
N
Me
O
N
S
O
t-Bu
SI-3
(151 MHz, CDCl3)
170.71 166.60 144.44 131.61 130.27 124.77 121.66 109.25
58.08 52.59 45.86 36.30 27.09 22.18
f1 (ppm)

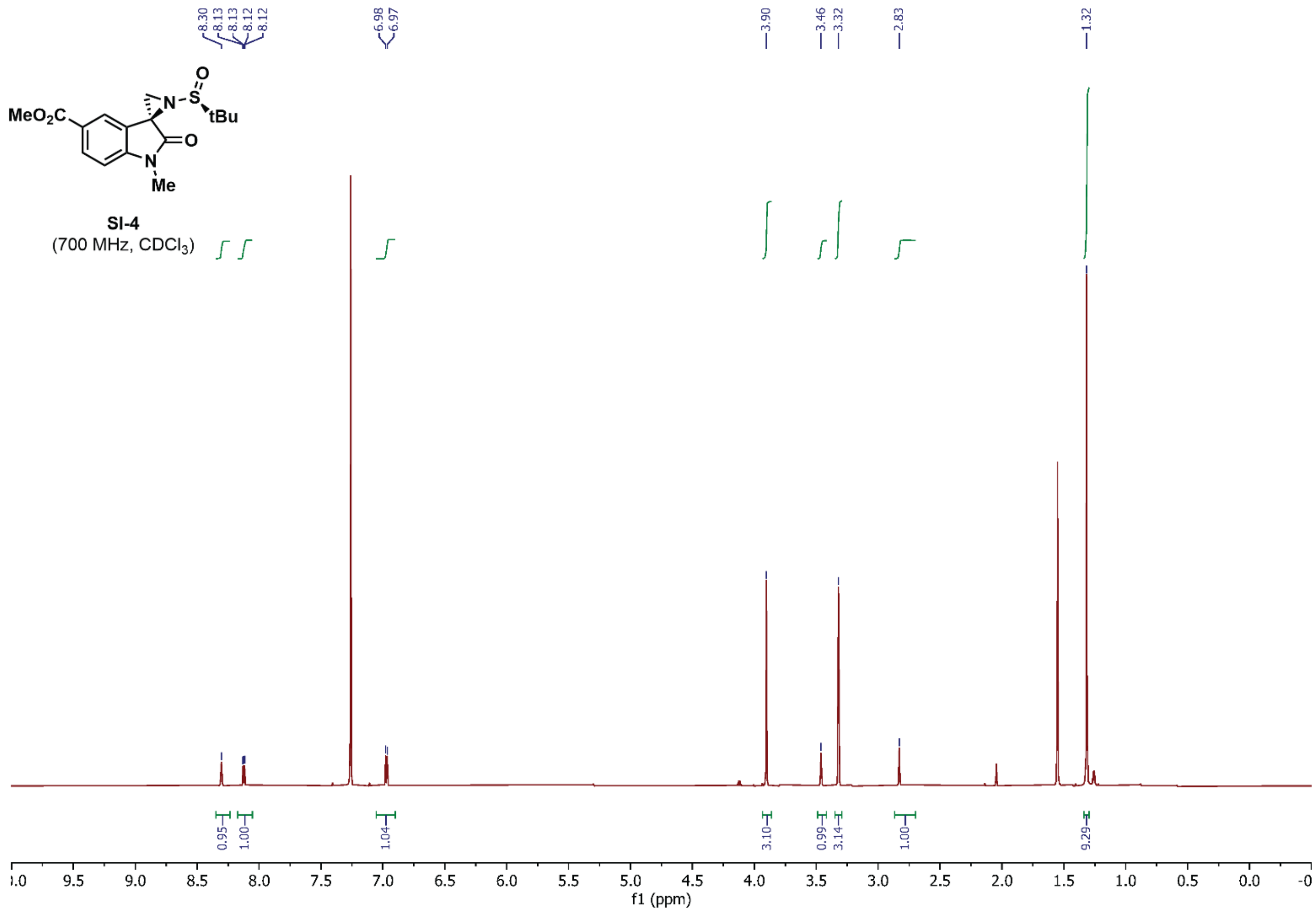


SI-4
(700 MHz, $CDCl_3$)

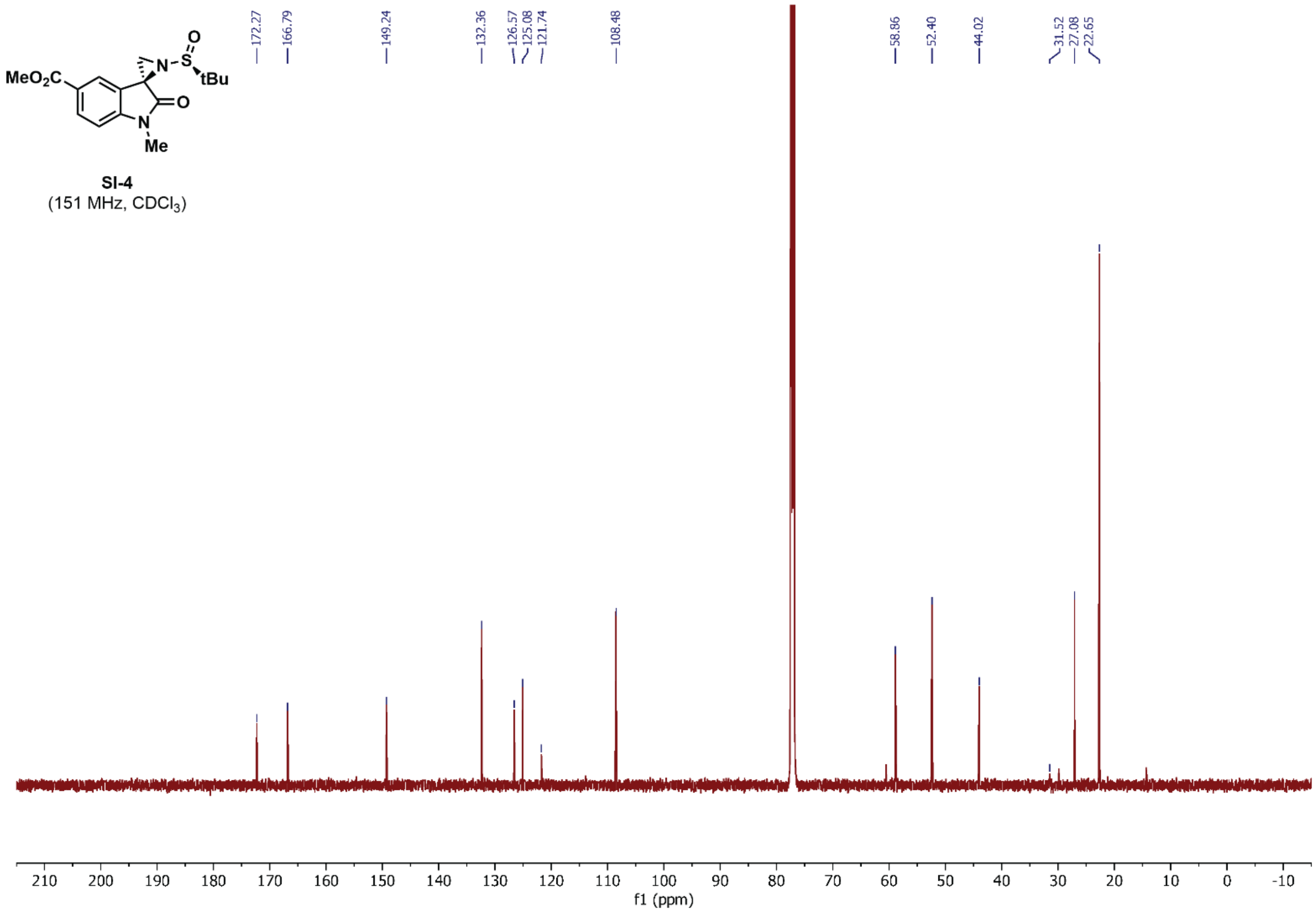


SI-4
(151 MHz, $CDCl_3$)

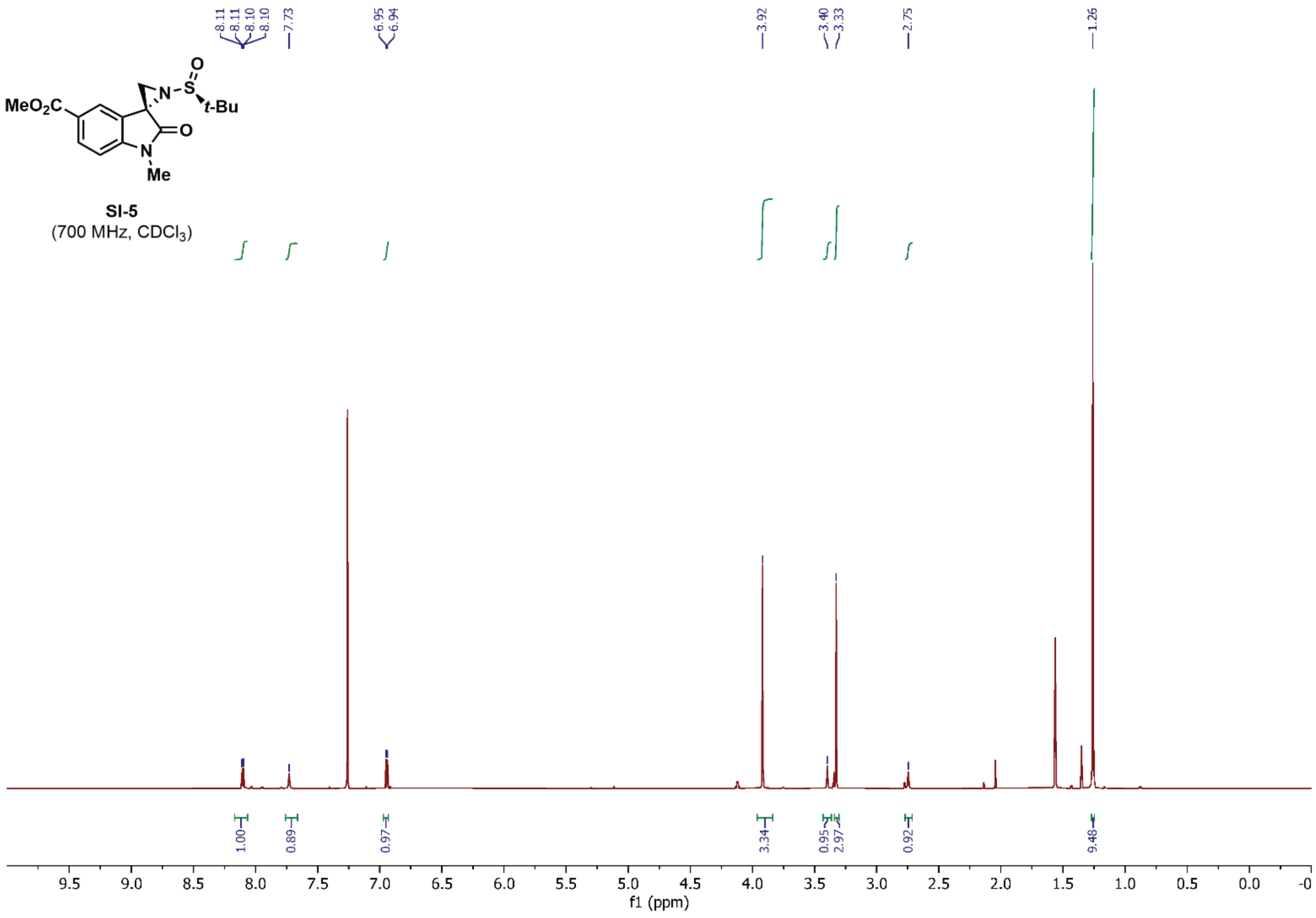

MeO2C
N
S
O
t-Bu
O
N
Me
SI-5
(700 MHz, CDCl3)
8.11
8.11
8.10
8.10
7.73
6.95
6.94
3.92
3.40
3.33
2.75
1.26
1.00
0.89
0.97
3.34
0.95
2.97
0.92
9.48
f1 (ppm)


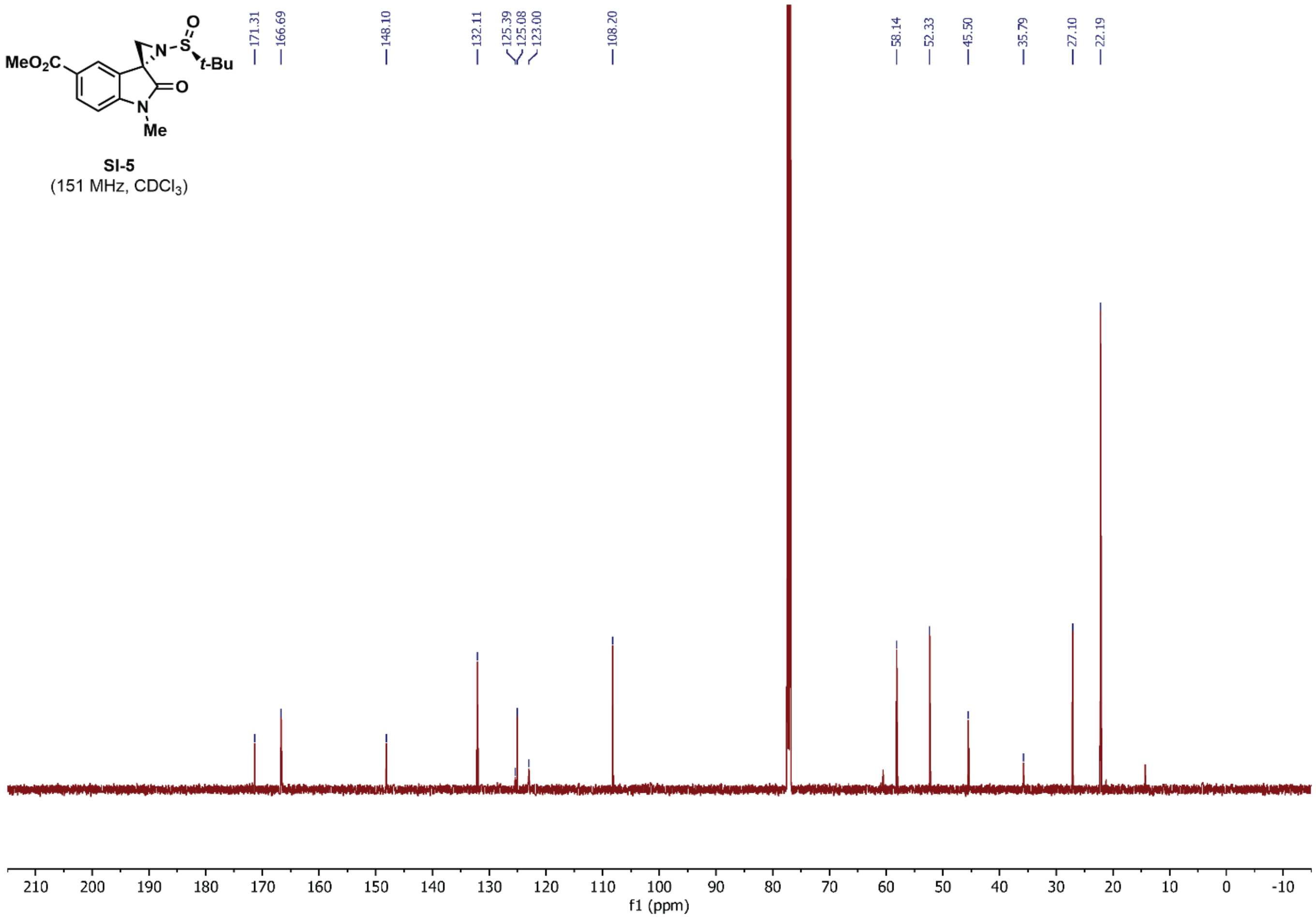

MeO2C
N
S
O
t-Bu
O
N
Me
SI-5
(151 MHz, CDCl3)
171.31
166.69
148.10
132.11
125.39
125.08
123.00
108.20
58.14
52.33
45.50
35.79
27.10
22.19
f1 (ppm)

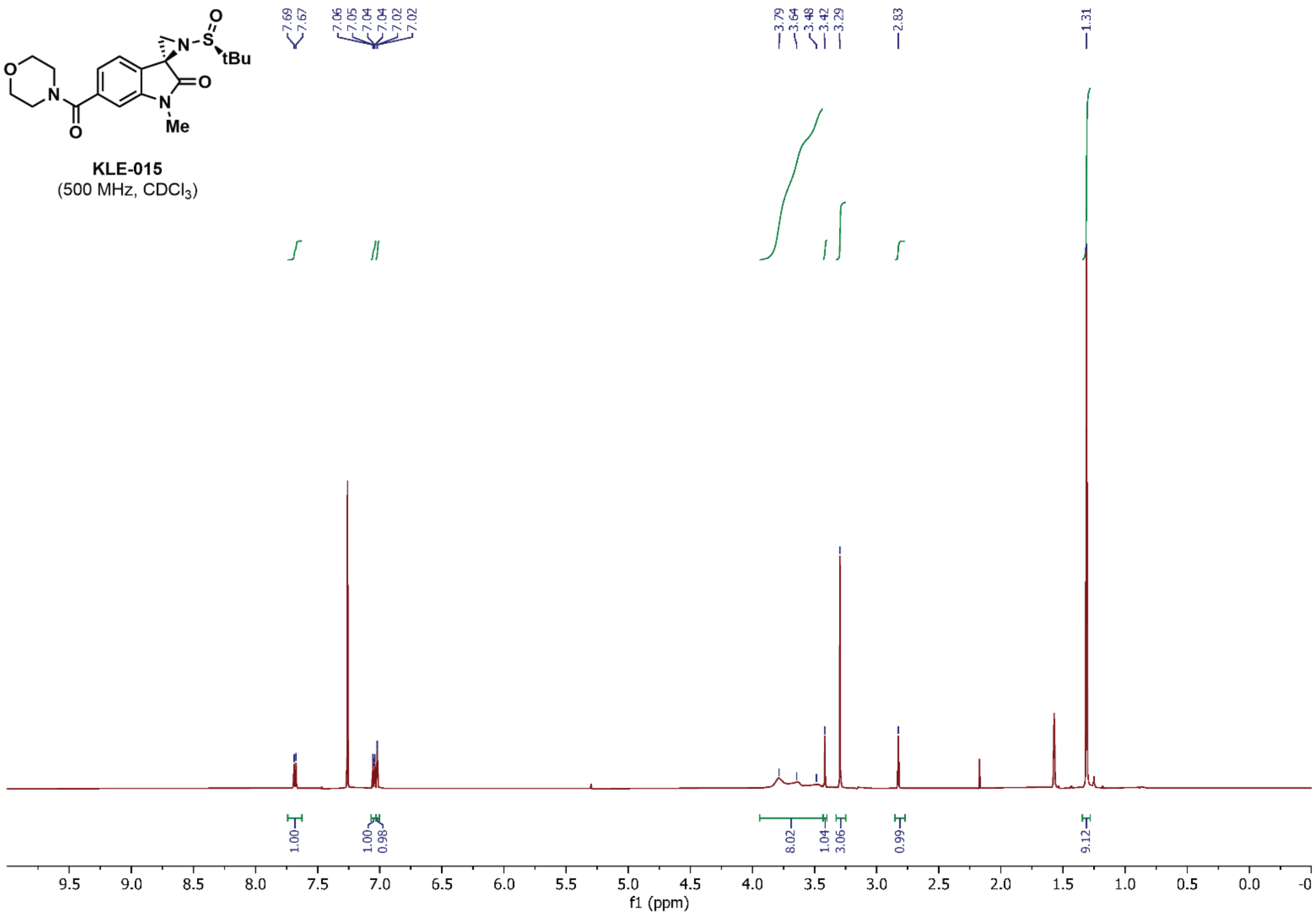


**KLE-015**
(500 MHz, $CDCl_3$)

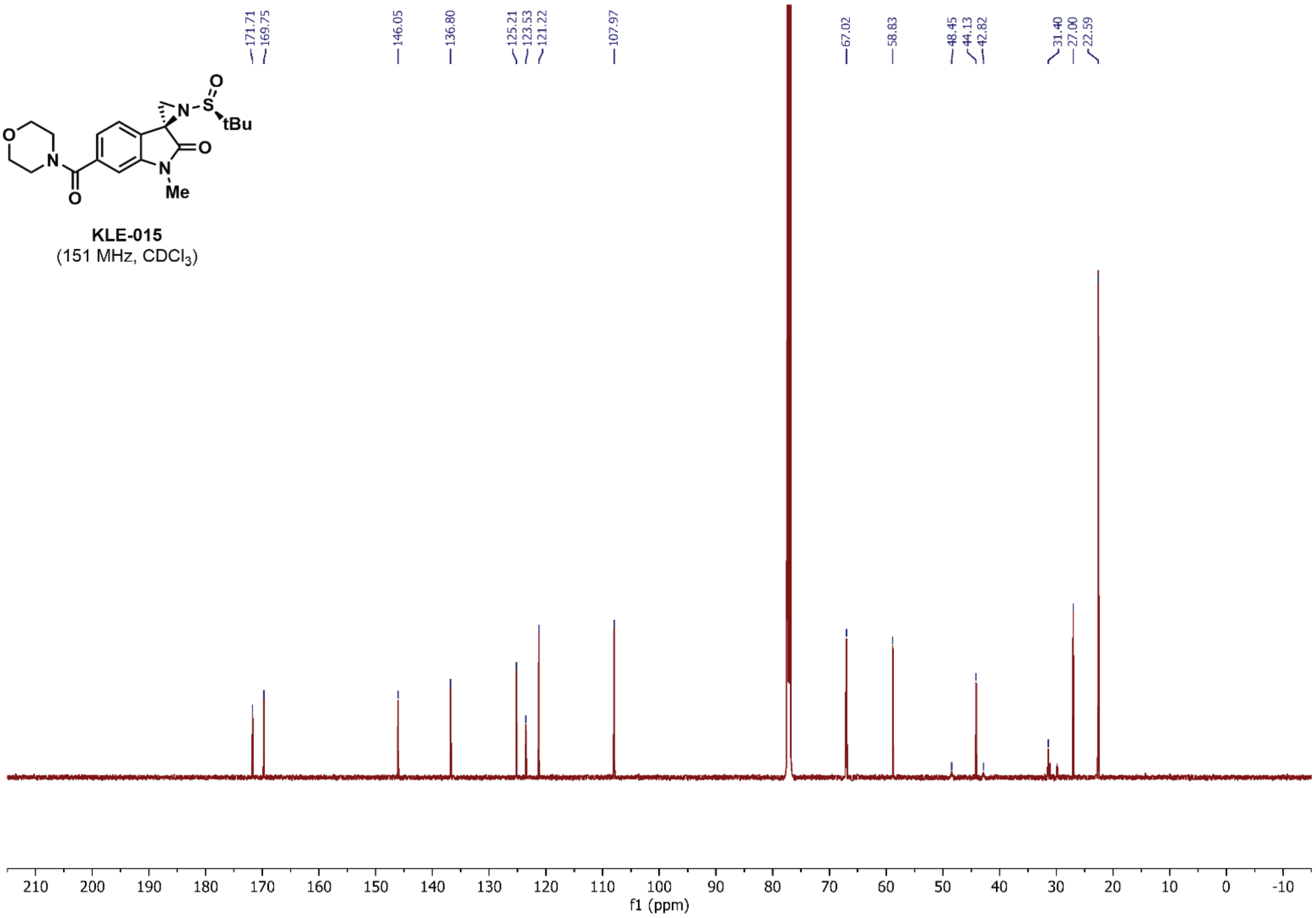


**KLE-015**
(151 MHz, $CDCl_3$)

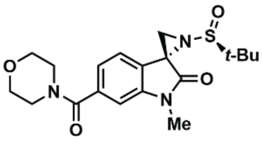

**KLE-016**
(500 MHz, $CDCl_3$)

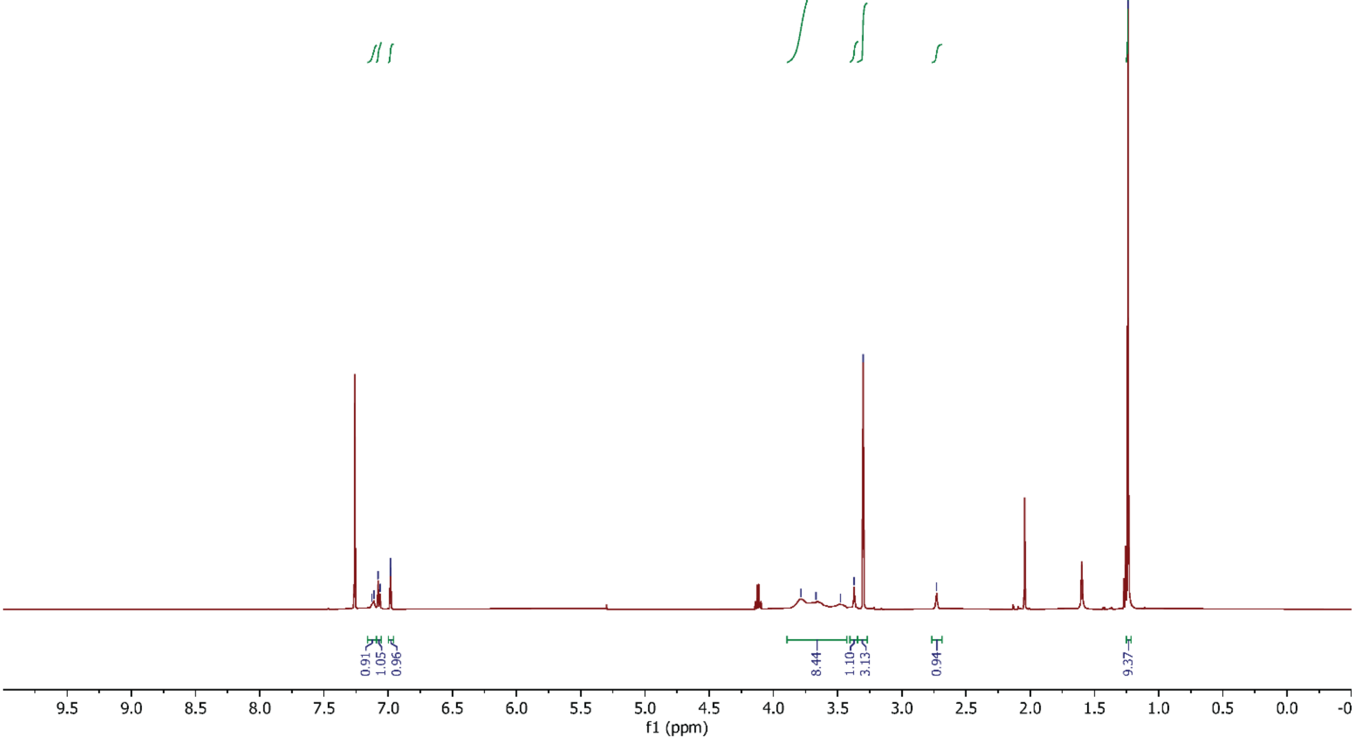


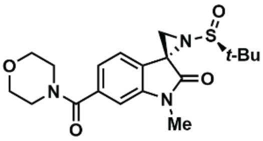

**KLE-016**
(151 MHz, $CDCl_3$)

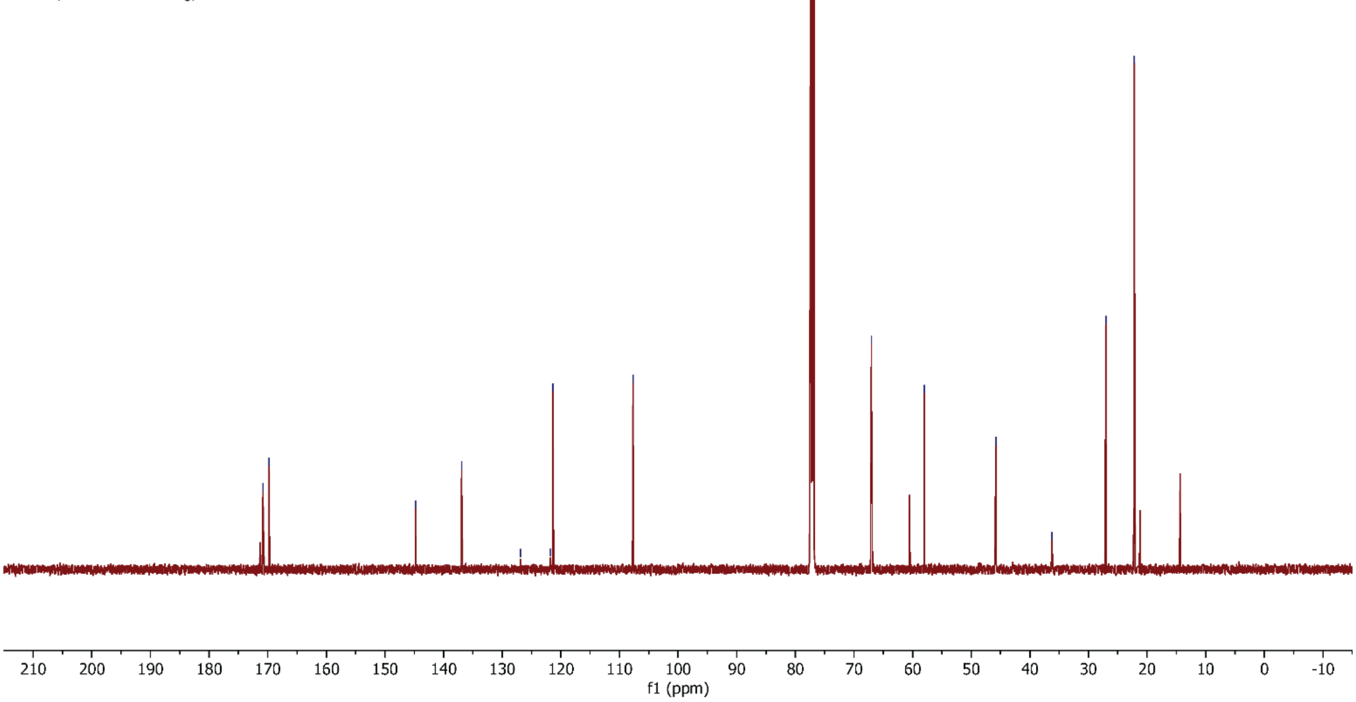

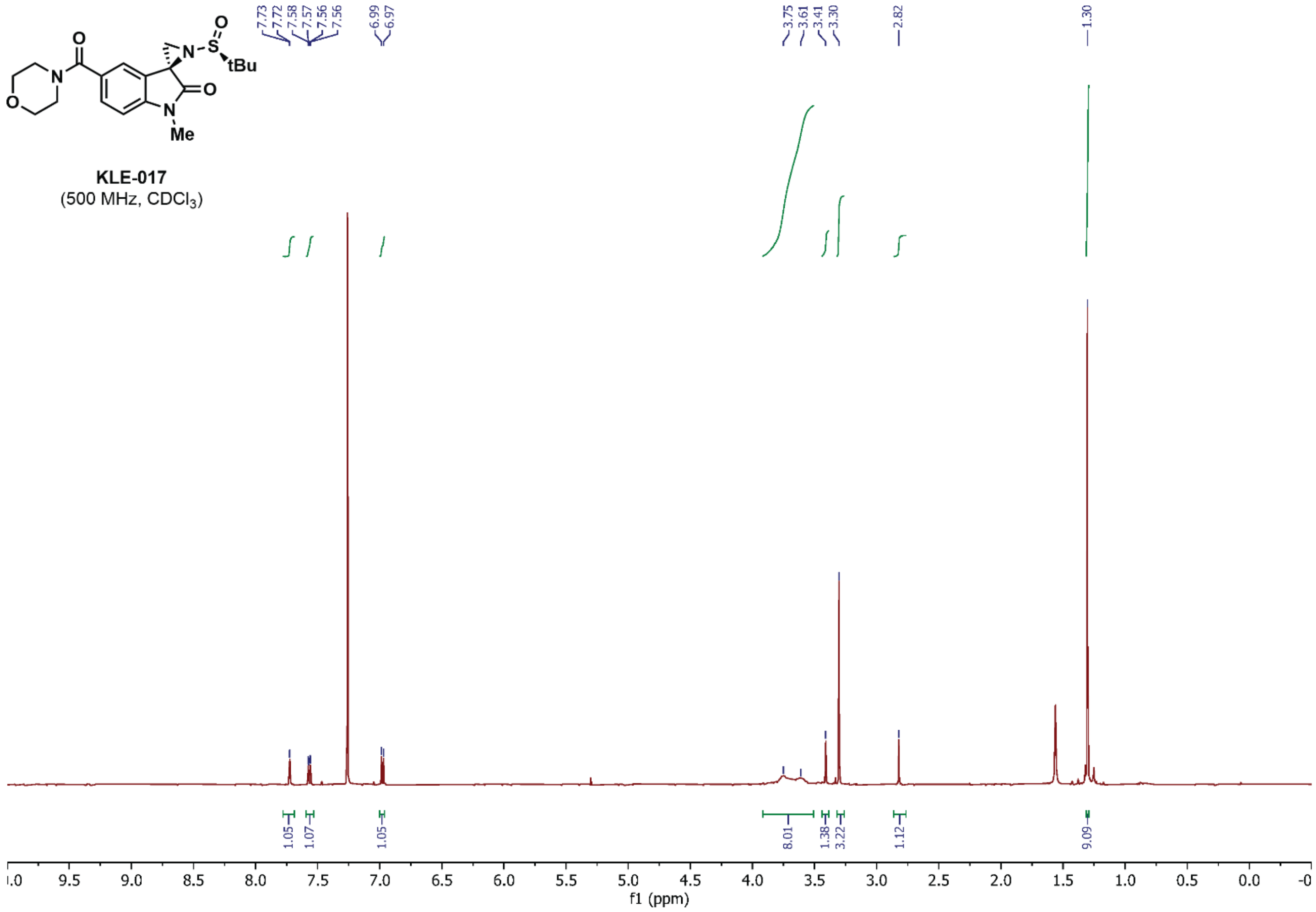

KLE-017
(500 MHz, $CDCl_3$)
7.73 7.72 7.58 7.57 7.56 7.56 6.99 6.97 3.75 3.61 3.41 3.30 2.82 1.30
1.05 1.07 1.05 8.01 1.38 3.22 1.12 9.09
f1 (ppm)


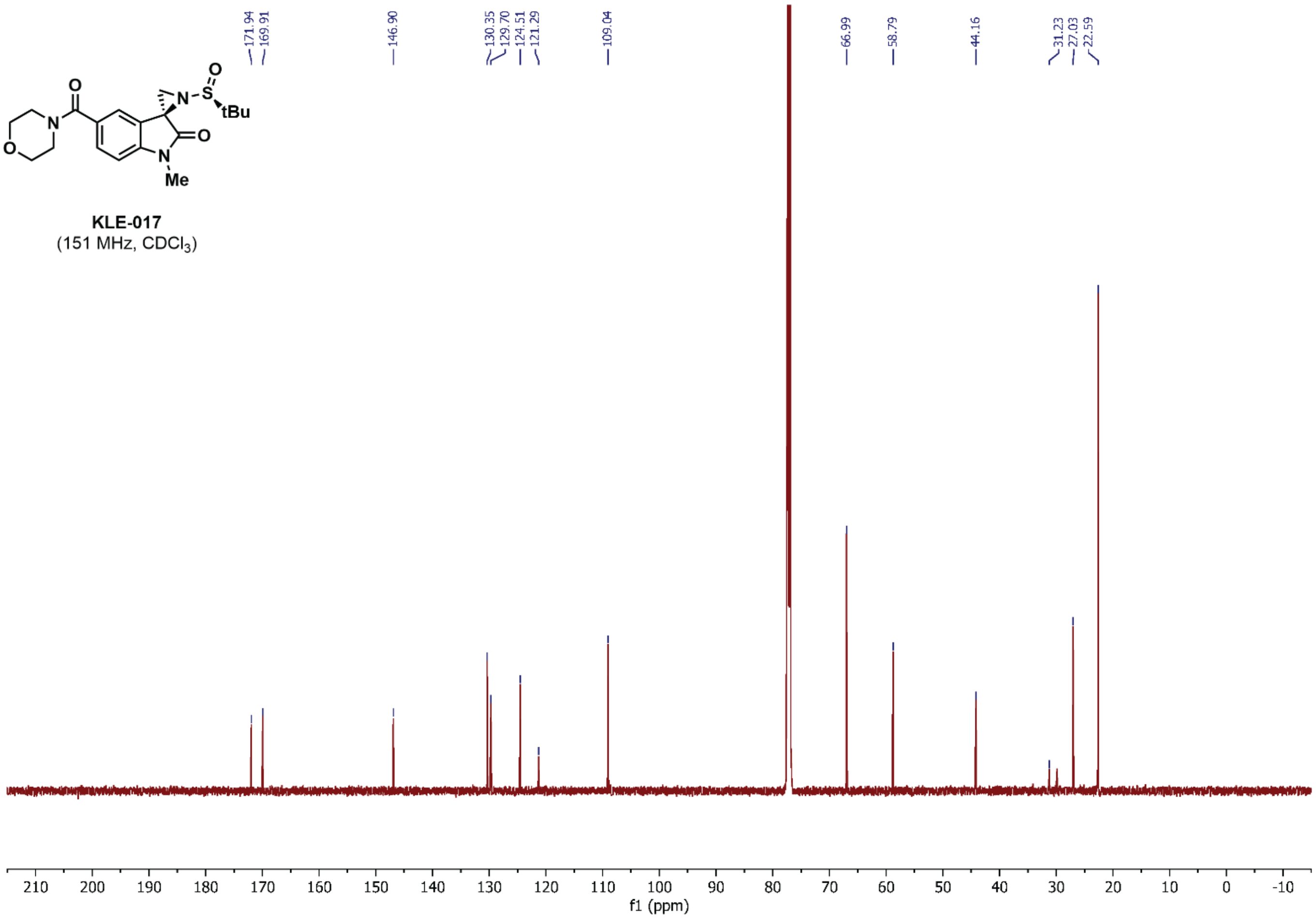

KLE-017
(151 MHz, $CDCl_3$)
171.94 169.91 146.90 130.35 129.70 124.51 121.29 109.04 66.99 58.79 44.16 31.23 27.03 22.59
f1 (ppm)

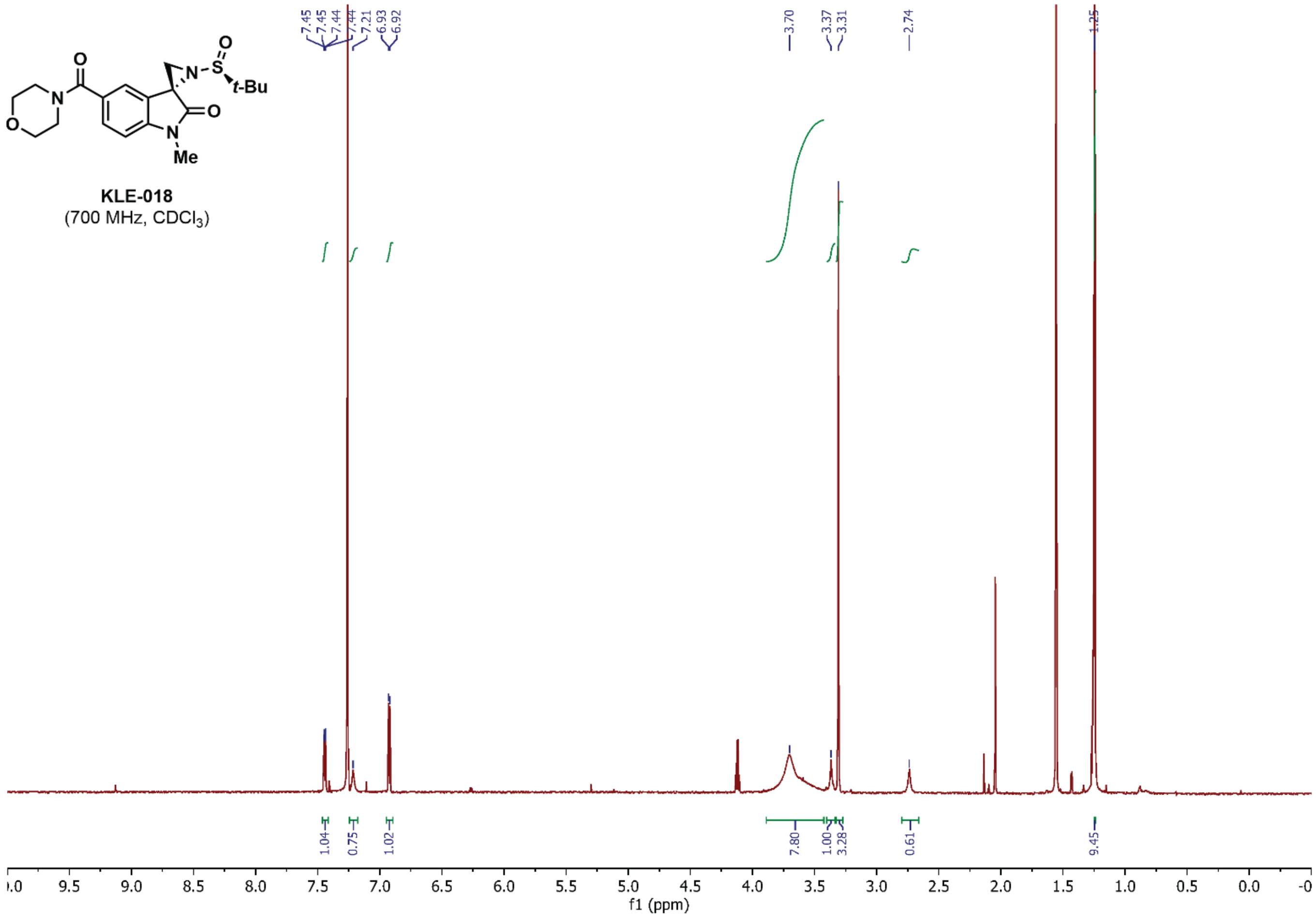
KLE-018
(700 MHz, $CDCl_3$)
7.45
7.45
7.44
7.44
7.21
6.93
6.92
3.70
3.37
3.31
2.74
1.25
1.04
0.75
1.02
7.80
1.00
3.28
0.61
9.45
f1 (ppm)

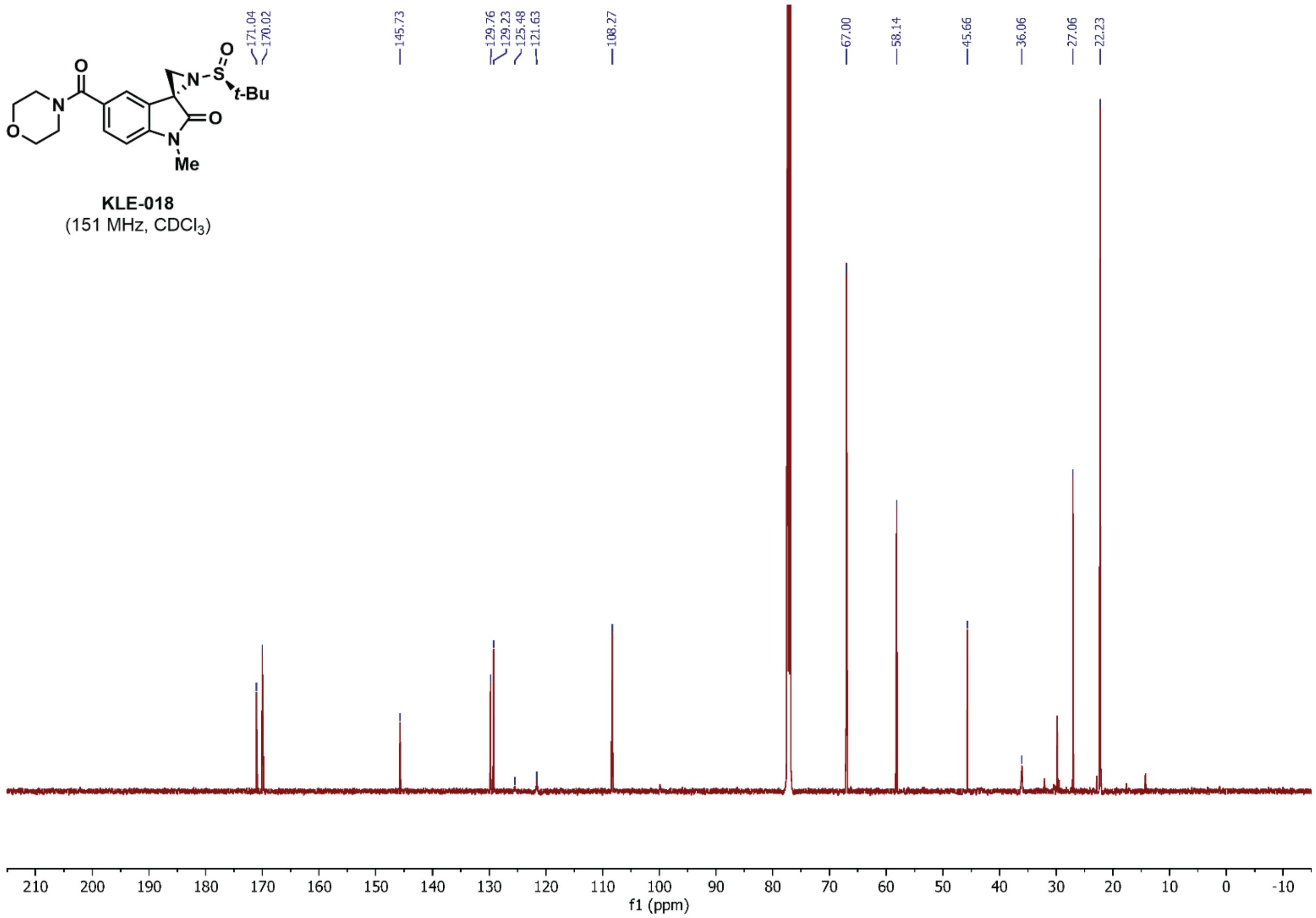
KLE-018
(151 MHz, $CDCl_3$)
171.04
170.02
145.73
129.76
129.23
125.48
121.63
108.27
67.00
58.14
45.66
36.06
27.06
22.23
f1 (ppm)

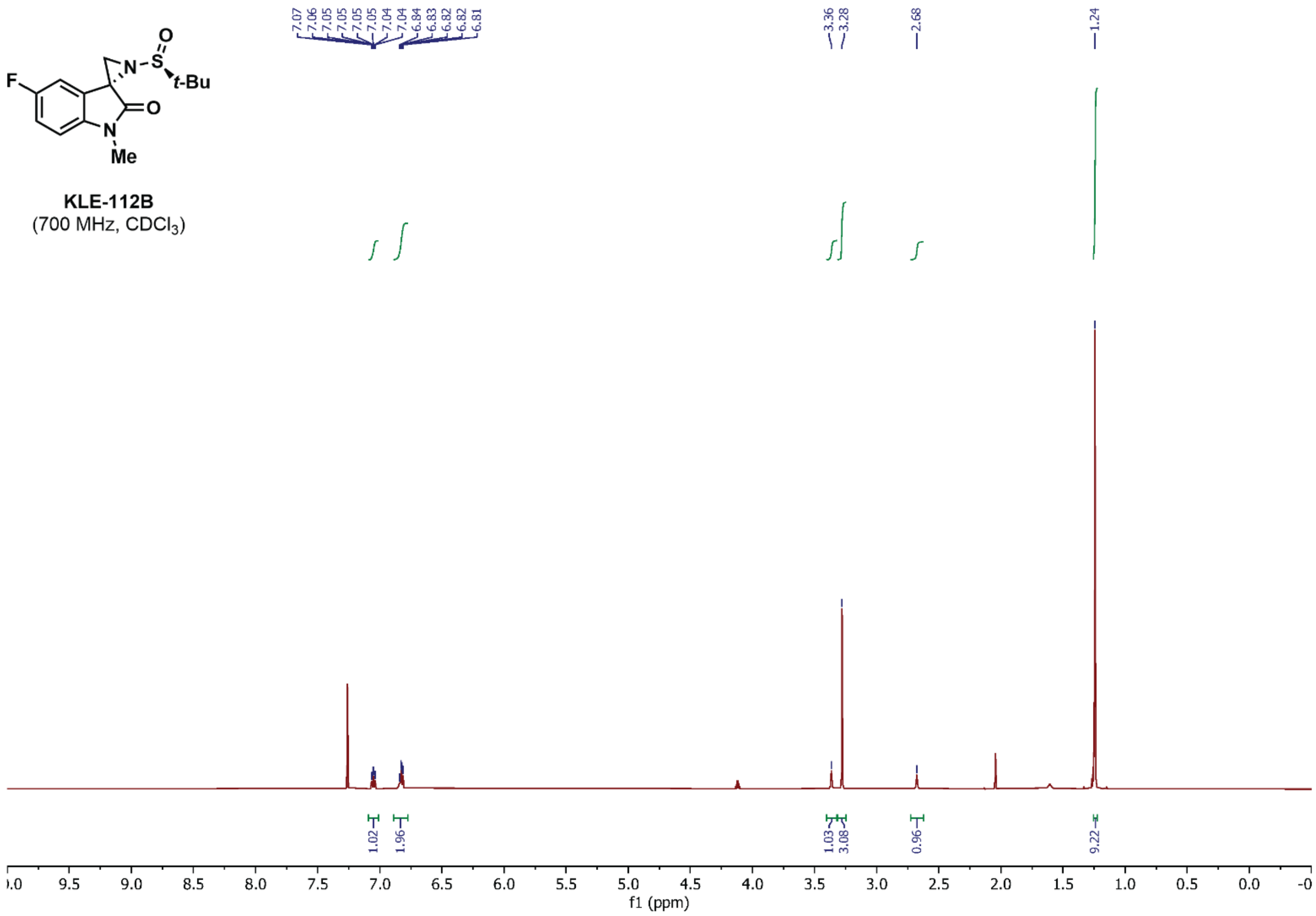
F
N
S
O
t-Bu
O
N
Me
KLE-112B
(700 MHz, $CDCl_3$)
7.07 7.06 7.05 7.05 7.05 7.05 7.04 7.04 6.84 6.83 6.82 6.82 6.81
3.36 3.28
2.68
1.24
1.02 1.96 1.03 3.08 0.96 9.22
f1 (ppm)

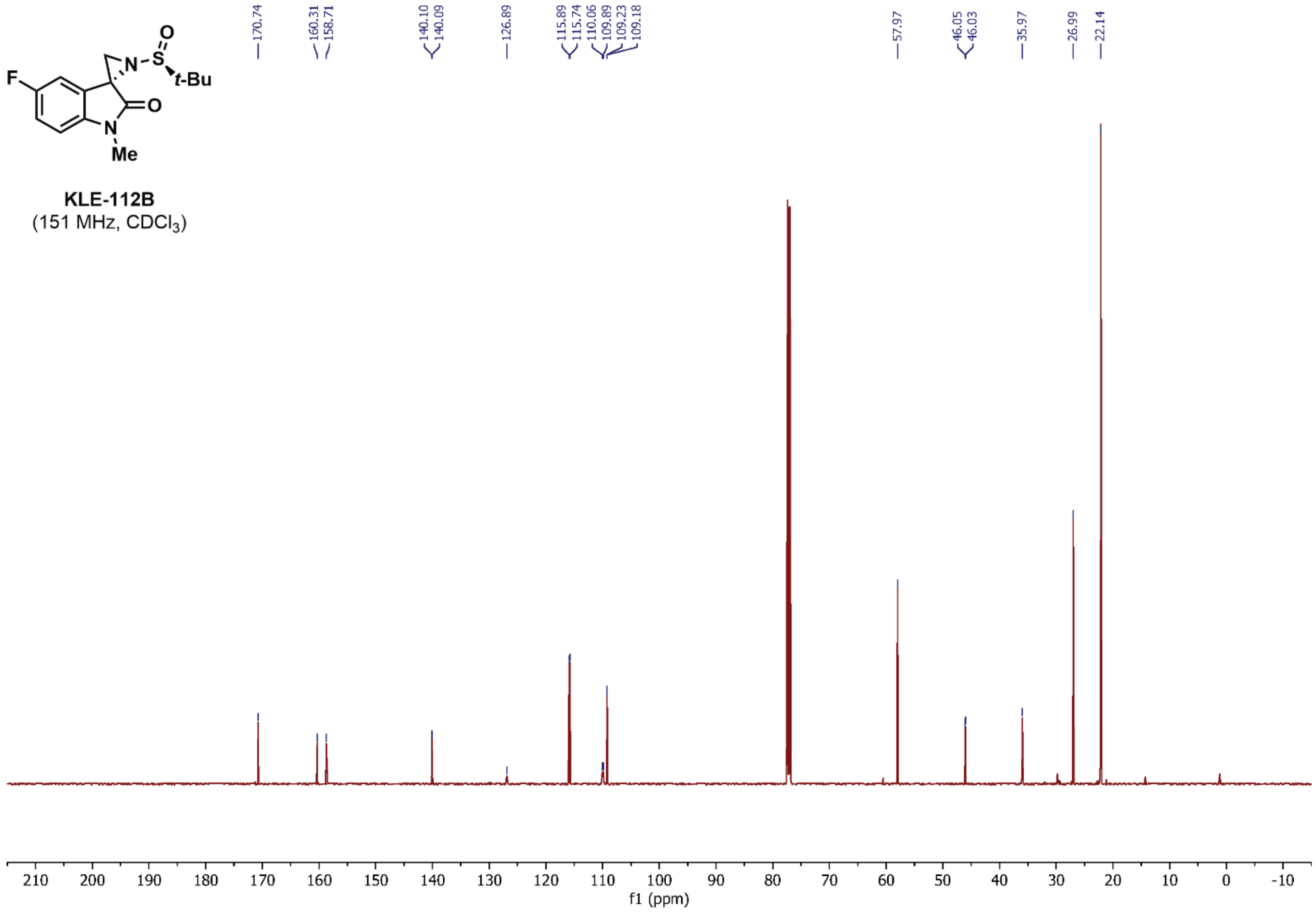
F
N
S
O
t-Bu
O
N
Me
KLE-112B
(151 MHz, $CDCl_3$)
170.74
160.31 158.71
140.10 140.09
126.89
115.89 115.74 110.06 109.89 109.23 109.18
57.97
46.05 46.03
35.97
26.99
22.14
f1 (ppm)

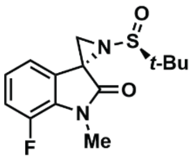


**KLE-113B**
(700 MHz, $CDCl_3$)

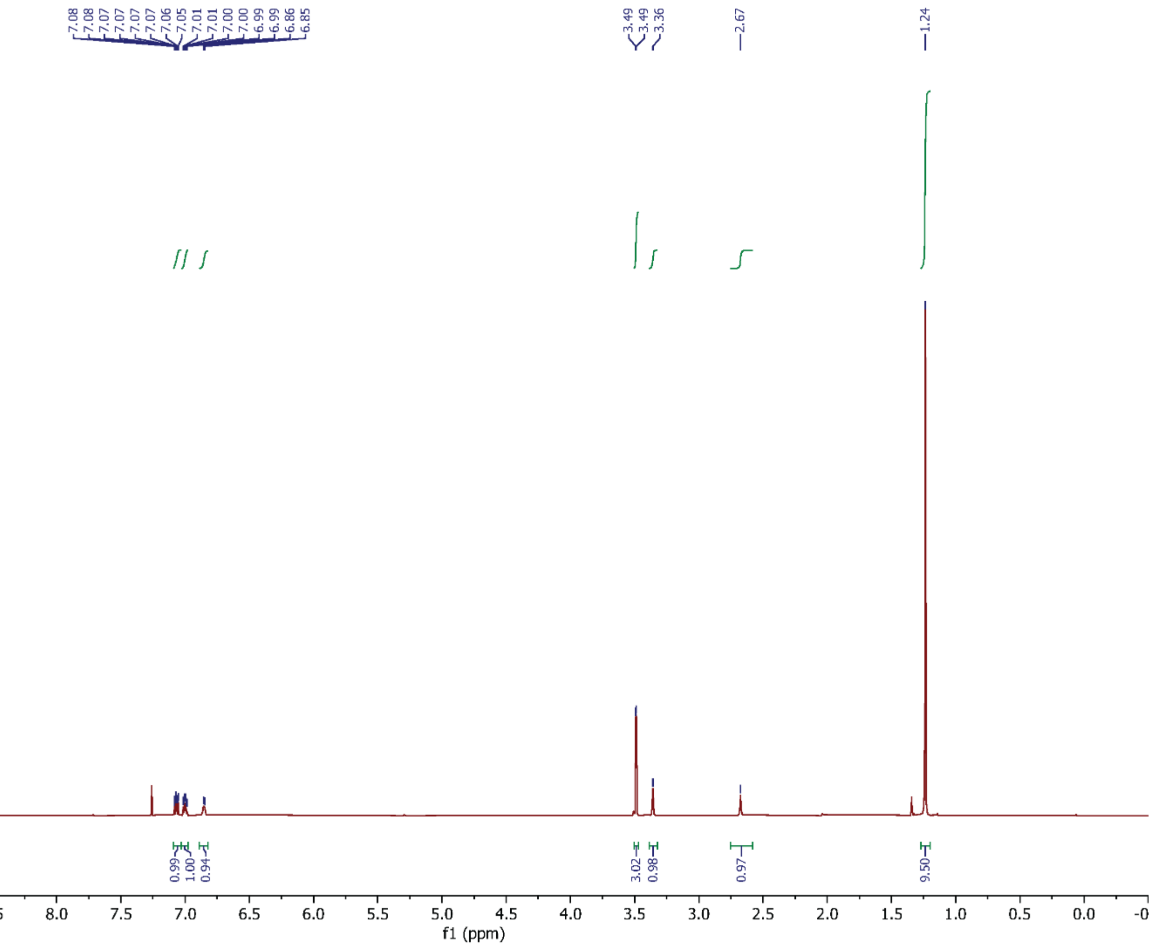


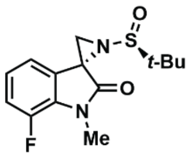


**KLE-113B**
(151 MHz, $CDCl_3$)

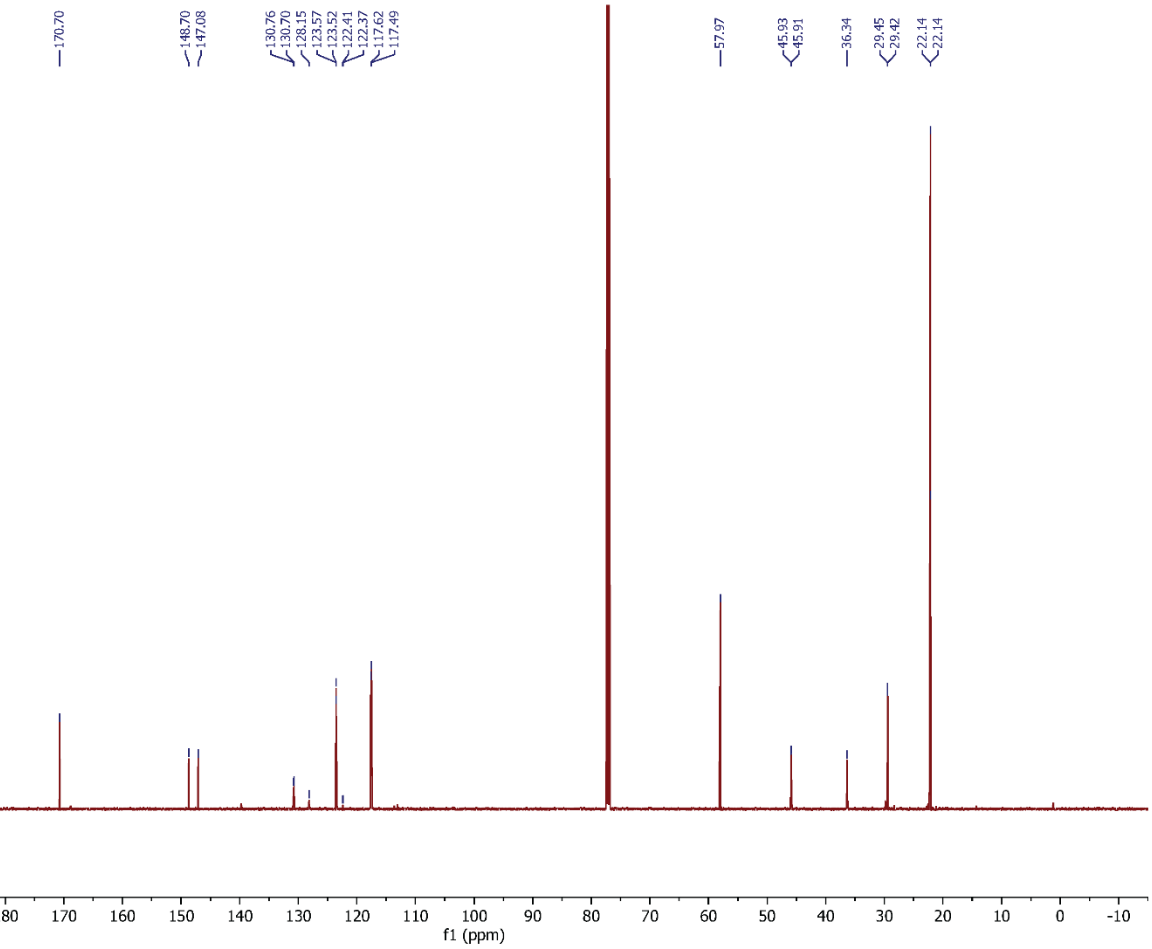

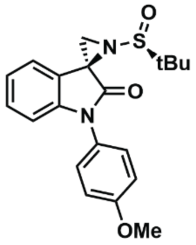


**KL4-219A**
(600 MHz, $CDCl_3$)

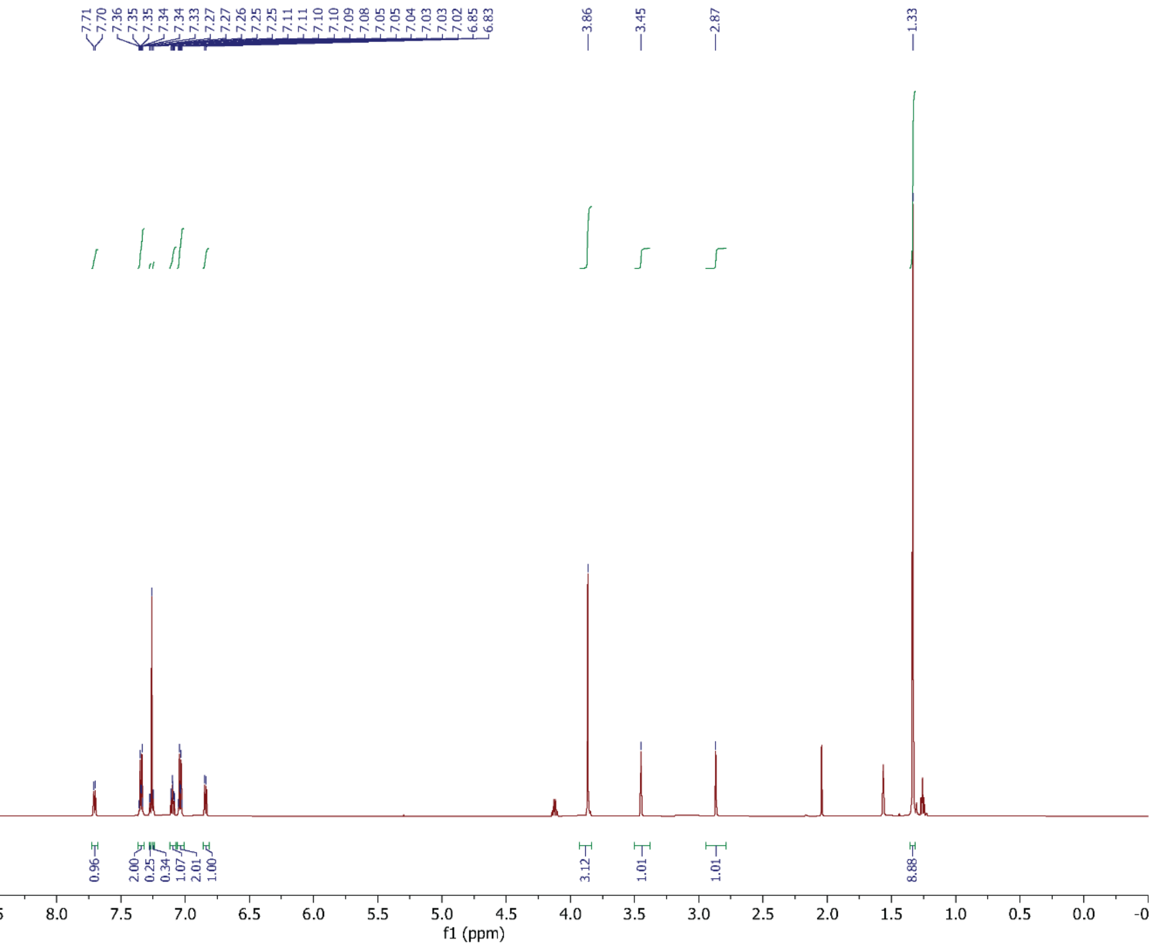


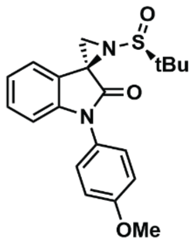


**KL4-219A**
(151 MHz, $CDCl_3$)

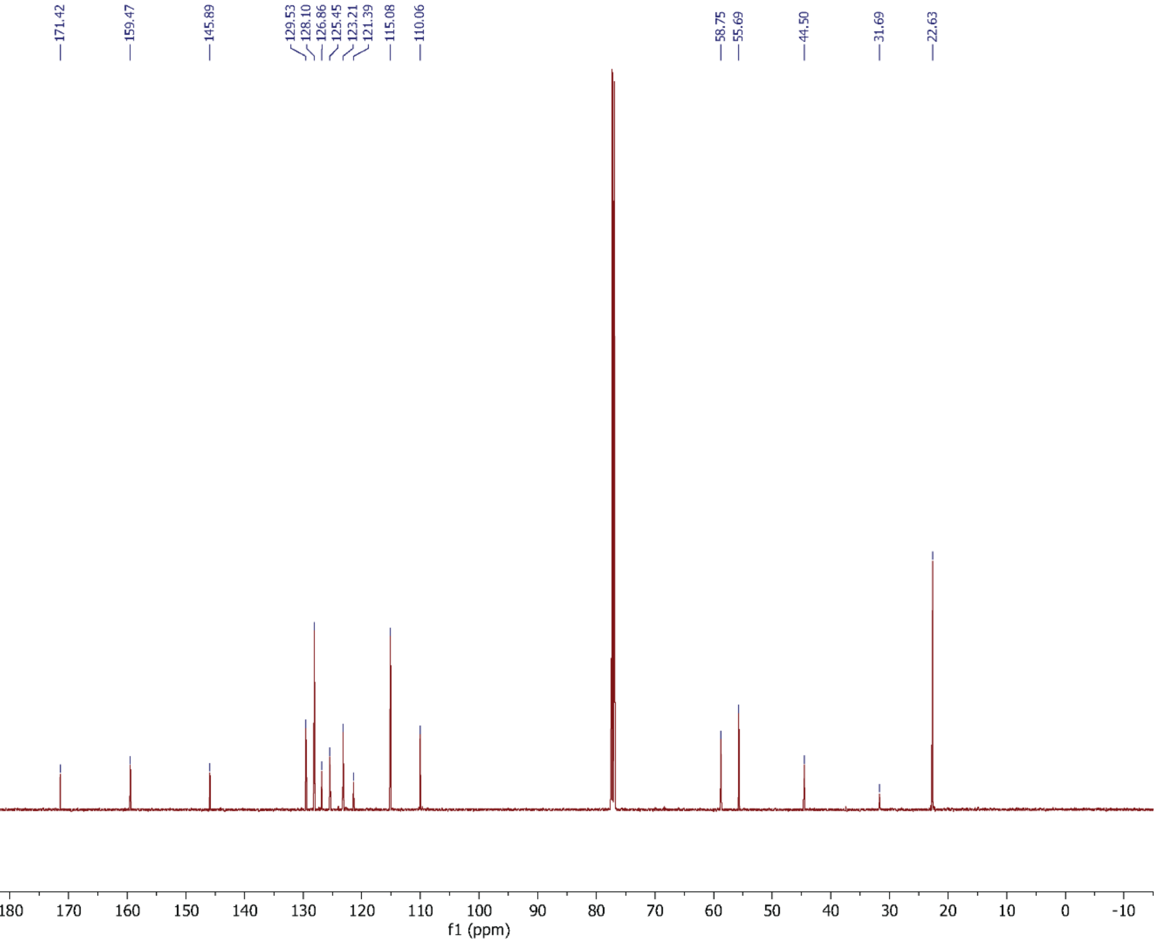

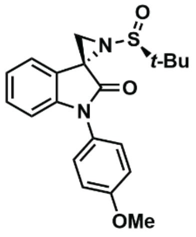


**KL4-219B**
(700 MHz, $CDCl_3$)

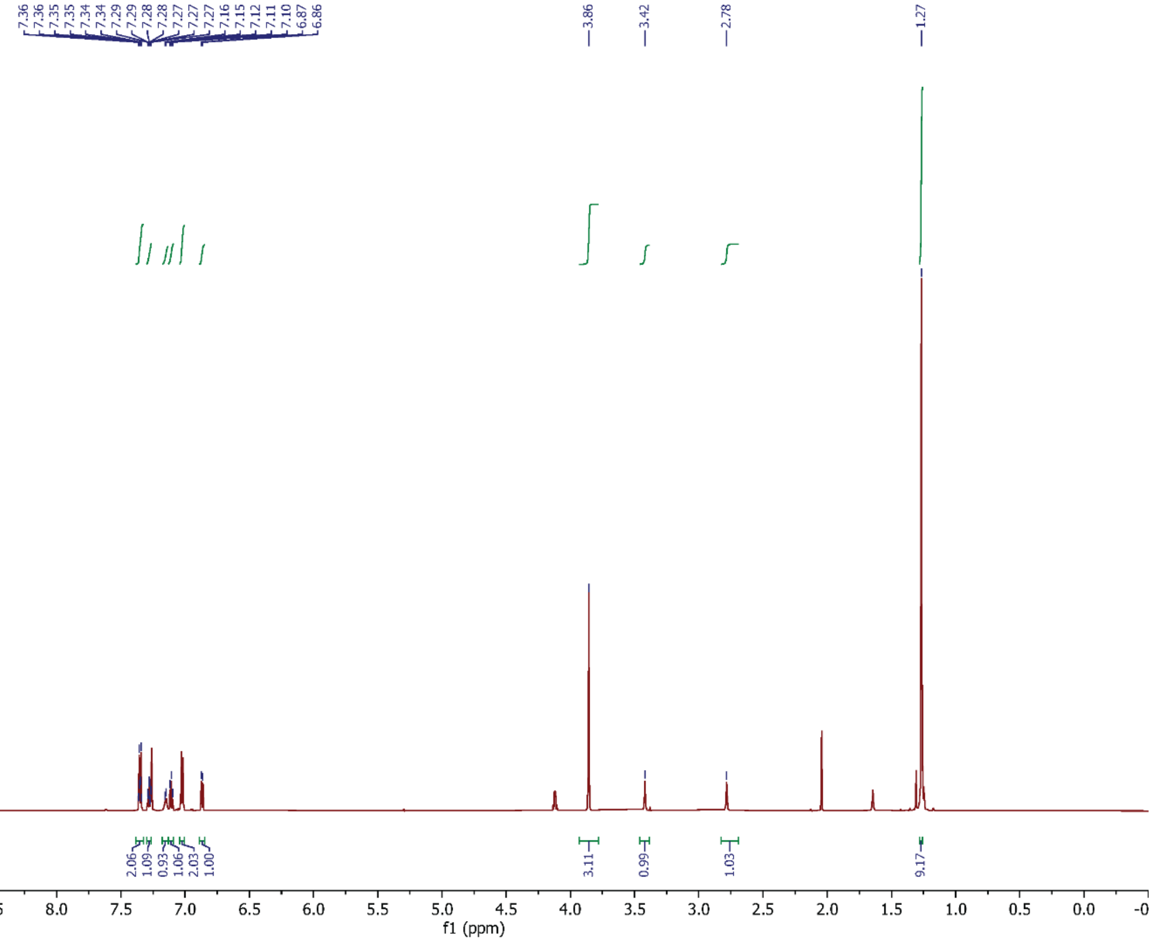


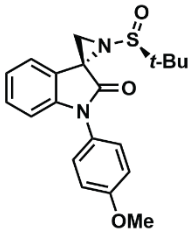


**KL4-219B**
(126 MHz, $CDCl_3$)

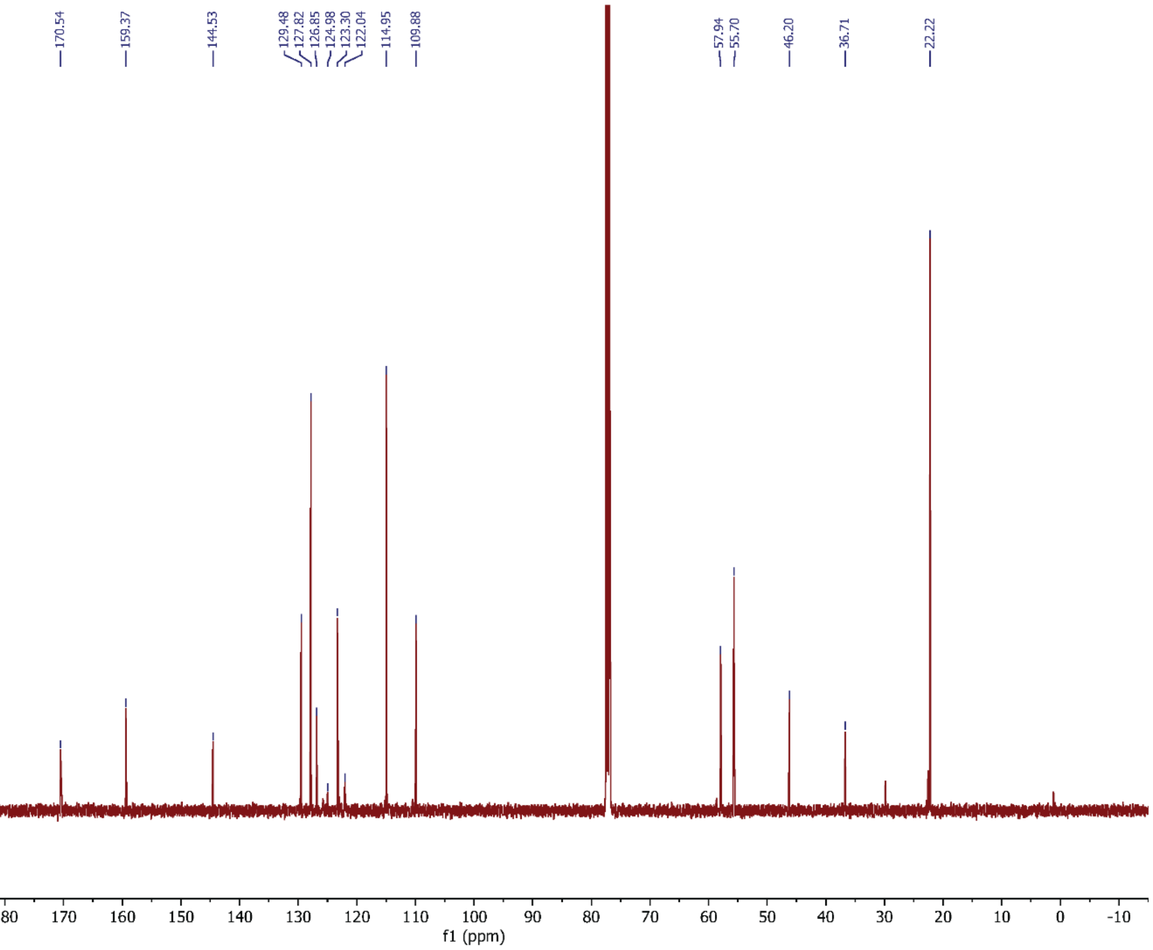
